# Genetic diversity and regulatory features of human-specific *NOTCH2NL* duplications

**DOI:** 10.1101/2025.03.14.643395

**Authors:** Taylor D. Real, Prajna Hebbar, DongAhn Yoo, Francesca Antonacci, Ivana Pačar, Danilo Dubocanin, Mark Diekhans, Gregory J. Mikol, Oyeronke G. Popoola, Benjamin J. Mallory, Mitchell R. Vollger, Philip C. Dishuck, Xavi Guitart, Allison N. Rozanski, Katherine M. Munson, Kendra Hoekzema, Jane E. Ranchalis, Shane J. Neph, Adriana E. Sedeño-Cortés, Benedict Paten, Sofie R. Salama, Andrew B. Stergachis, Evan E. Eichler

## Abstract

*NOTCH2NL* (*NOTCH2*-N-terminus-like) genes arose from ape-specific chromosome 1 segmental duplications implicated in human brain cortical expansion, including an incomplete *NOTCH2* gene. Genetic characterization of these loci and their regulation is complicated because they are embedded in large, nearly identical duplications that predispose to recurrent microdeletion syndromes. Using near-complete long-read assemblies generated from 70 human and 12 ape haploid genomes, we show independent recurrent duplication among apes with protein-coding copies emerging in humans 2.2-3.7 million years ago. We distinguish *NOTCH2NL* paralogs present in every human haplotype (*NOTCH2NLA*) from copy number variable ones. We also characterize large-scale structural variation, including gene conversion, for 28% of haplotypes leading to a previously undescribed paralog, *NOTCH2tv.* Finally, we apply Fiber-seq and long-read transcript sequencing to human dorsal forebrain organoids to characterize the regulatory landscape and find that the most fixed paralogs, *NOTCH2* and *NOTCH2NLA*, harbor the greatest number of paralog-specific elements potentially driving their regulation.

## INTRODUCTION

Notch signaling, a mechanism of cell communication conserved throughout the metazoan kingdom, is uniquely altered in humans due to a recent ape segmental duplication of the *NOTCH2* gene^1,2^. Segmental duplications (SDs) have both restructured primate genomes as well as led to the emergence of lineage-specific gene families resulting in potentially new functions, including distinct developmental fates^3,4^). In the case of *NOTCH2NL* (*NOTCH2-N-terminus-like*), the gene family is a ∼70 kbp SD encompassing the four N-terminal exons of *NOTCH2* and includes a unique final fifth exon exapted from the fourth intron of *NOTCH2*. While *NOTCH2NL*-like sequences exist in several primates, the NOTCH2NL protein appears to only be expressed in the developing human brain, suggesting it is functionally human specific^1^. In humans, there are three truncated paralogs, *NOTCH2NLA*, *NOTCH2NLB* and *NOTCH2NLC*; all are expressed and the predicted proteins only differ by a few amino acids^1^. Experimental work has shown that NOTCH2NL interacts with NOTCH2 and modulates the NOTCH2-signaling pathway^1,2^). Specifically, NOTCH2NL increases the number of self-renewal divisions of progenitor radial glia while delaying differentiation of these cells into neurons. Prioritizing self-renewal over differentiation has been proposed to enable the human brain to increase neuronal mass during cortical neurogenesis^1,2^). In addition to its potential role in the expansion of the human brain, mutations in the *NOTCH2/NL* gene family or their associated SDs underlie four distinct genetic disorders, including Alagille syndrome^5,6^), neuronal intranuclear inclusion disease^7,8^, and chromosome 1q21.1 distal duplication/deletion and TAR (thrombocytopenia-absent radius) syndromes^9,10,11^).

Historically, complex high-identity regions enriched in SDs, like the *NOTCH2NL* locus from chromosome 1p12-1q21, have been difficult to sequence and assemble with short-read technologies. Duplicated *NOTCH2NL* copies are embedded in much larger blocks (often hundreds of kbp to a few Mbp in length) of SDs associated with other genes such as the core duplicon *NBPF* (neuroblastoma breakpoint family)^12,13,14,15,1,16^. These regions, subject to different mutational processes such as interlocus gene conversion (IGC)^17^, make paralogous sequence variants unreliable as tags unless such common IGC patterns in diverse humans are characterized. As a result, studies of human genetic variation have frequently excluded these regions; standard genome-wide association studies and attempts to functionally characterize via ENCODE and GTEx are almost nonexistent due to their dependence on short-read sequencing platforms^18,19^.

In this study, we address these limitations by using long-read sequencing data and associated pangenome and telomere-to-telomere (T2T) resources generated from nonhuman primates (NHPs) and a diverse set of humans (Human Pangenome Reference Consortium [HPRC])^20,3,21^. Our goal was to annotate structural differences, expression, and regulatory changes in the context of human genetic variation—a feat only possible with completely resolved haplotype sequence. Understanding variation in patterns of the human haplotypes will have the added benefit of helping to define breakpoints associated with patients harboring chromosome 1q21 deletions and duplications in the future^10,11^).

## RESULTS

### Structure of the *NOTCH2NL* gene family in a complete human genome assembly

In the fully assembled T2T-CHM13 haploid genome^22^, *NOTCH2NLA*, *NOTCH2NLB*, and *NOTCH2NLC* map to the q-arm (Figure 1A) and define the functional human-specific duplications. All daughter duplications are approximately 10-11 kbp proximal upstream of an *NPBF* gene, which has been implicated as a partner in creating *NOTCH2NL* fusion transcripts (Figure 1A)^23,16^. In T2T-CHM13, *NOTCH2NLA* and *NOTCH2NLC* gene models are distinct from the other paralogs because their first exon is suggested by gene annotation to be untranslated (Figure 1B), due to unique mutations that remove the canonical *NOTCH2* initiator methionine and secretory signal^1^. We annotated all structural variants >500 bp among the paralogs (Table S1) and defined synteny stretches extending beyond the *NOTCH2NL* genes (Table 1, Figure 1C). We also used the order and orientation of duplicons flanking each *NOTCH2NL* copy as defined by DupMasker^24^ to generate a “barcode” of each locus to readily identify regions in other human genomes when compared to T2T-CHM13 (Figure 1B, Methods). Representing the surrounding sequence in terms of the higher-order duplication content over 1 Mbp regions helped define orthologous locations in the presence of the homogenizing effects of IGC^25^. Importantly, all copies of *NOTCH2NL* show one breakpoint with respect to the ancestral *NOTCH2* corresponding to the *NBPF* duplicon that demarcates the 3’ end of each derived-duplicated gene (Figure 1C). Additionally, the presumptive pseudogene *NOTCH2NLR* is missing the upstream nongenic ancestral *NOTCH2* sequence present in *NOTCH2NLA* and *NOTCH2NLB*.

**Figure 1.**
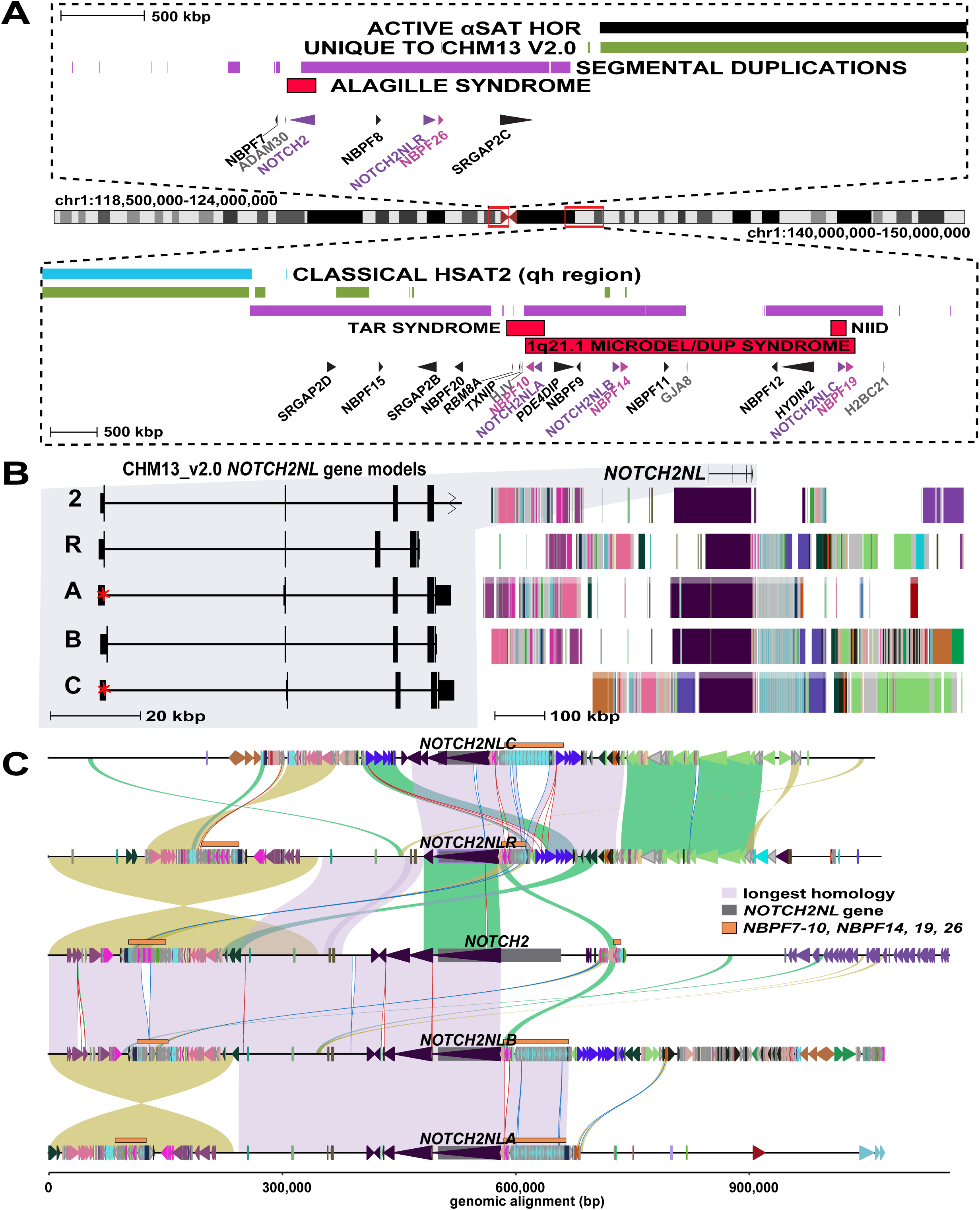
Genome structure and organization of the *NOTCH2NL* gene family. **A.** Long-range organization of *NOTCH2/NOTCH2NL* loci in the T2T-CHM13 reference genome, including centromere satellite annotations of active alpha satellite (*α*Sat) higher-order repeats (HORs) (black) and classical human satellite 2 (hsat2, secondary constriction [qh] region^57^) (blue), regions unique to the T2T-CHM13 assembly (green), intervals of segmental duplications (SDs) (purple), and Mendelian and genomic disorders associated with specific regions/paralogs (red). A subset of genes is depicted, including *NOTCH2NL* (purple), *NBPF* genes that are directly downstream of *NOTCH2NL* (pink), non-duplicated genes (gray) immediately adjacent to the SD blocks, and others (black). **B.** Duplicon organization as defined by DupMasker (Methods, Table S6) flanking the *NOTCH2NL* region and intron/exon structure of genes in T2T-CHM13 V2.0^58^ (http://genome.ucsc.edu). Red asterisks mark the nontraditional CTG start that the browser annotations do not take into consideration. **C.** Stacked SVbyEye plot of 1 Mbp regions flanking human *NOTCH2NL* genes (gray squares), contrasting syntenic regions in direct orientation (green/lavender) versus inverted alignments (yellow). Annotations include different *NBPF* genes in the region (orange). Note: the two large inversions between *NOTCH2/NOTCH2NLR* and *NOTCH2NLA*/*NOTCH2NLB*, respectively, are the result of proximity due to overlapping sequence. Duplicons as defined by DupMasker (colored triangles). See also Tables S1 and S6.

**Table 1.** Greater *NOTCH2NL* region pairwise homology matrix in T2T-CHM13.

| **Table 1. Greater *NOTCH2NL* region pairwise homology matrix in T2T-CHM13** |  |  |  |  |  |
| --- | --- | --- | --- | --- | --- |
|  | *NOTCH2* | *NOTCH2NLR* | *NOTCH2NLA* | *NOTCH2NLB* | *NOTCH2NLC* |
| *NOTCH2* | x | 115,690 | 326,052 | 556,583 | 118,987 |
| *NOTCH2NLR* | 99.7 | x | 342,476 | 328,624 | 217,296 |
| *NOTCH2NLA* | 99.3 | 99.3 | x | 416,088 | 182,193 |
| *NOTCH2NLB* | 99.3 | 99.3 | 99.7 | x | 265,598 |
| *NOTCH2NLC* | 99.2 | 99.0 | 99.1 | 99.2 | x |

### Independent *NOTCH2NL* duplications and large-scale restructuring of ape chromosome 1

We compared the extent of synteny of the corresponding *NOTCH2NL* loci among finished NHP genomes^20,3^ (Figure S1, Figure S2A). We identified 26 distinct *NOTCH2NL* SDs among nonhuman apes (NHAs) (average of 215 kbp) (Table S2). We find that the mean SD length between *NOTCH2NL* SDs in humans (311 kbp) and other NHAs (215 kbp) is not significantly different (p=0.42; t-test two-sided), indicating that this locus was unstable and began to duplicate in the common ancestor of the great apes. Consistent with Fiddes et al. (2018)^1^, all NHA homologs appear truncated with respect to *NOTCH2*. All are missing different canonical exons relative to the known human *NOTCH2NL* gene models (Figure 1B). Our comparative analysis (Figure 2) indicates that three distinct inversion events occurred during the evolution of the chromosome 1p21.2-q23.2 region. The first inversion (I) occurred in the ancestor of African great apes, flipping the region into its current orientation in gorilla, chimpanzee, and bonobo. Subsequently, in the human lineage, this region reverted to its ancestral configuration (inversion II). Additionally, an expansion of SDs in humans coincided with a human-specific pericentric inversion (III), corresponding to the rearrangement originally described by Yunis and Prakash^26^ and subsequently refined to 154 kbp and 562 kbp breakpoint intervals at chromosome 1p11.2 and 1q21.3, respectively^27^. We estimate that the region encompassing inversions II and III is 17 Mbp larger than the syntenic region in chimpanzee and orangutan. In humans this event uniquely positioned *NOTCH2NLA*, *NOTCH2NLB*, and *NOTCH2NLC* on the long arm of chromosome 1, splitting the *NOTCH2NL* locus across the centromere when it had previously always existed on a single chromosome arm (Figure S2B). The region encompassing *NOTCH2NL* paralogs has undergone significant restructuring via duplication and inversion among all great apes, but especially in the human genome.

**Figure 2.**
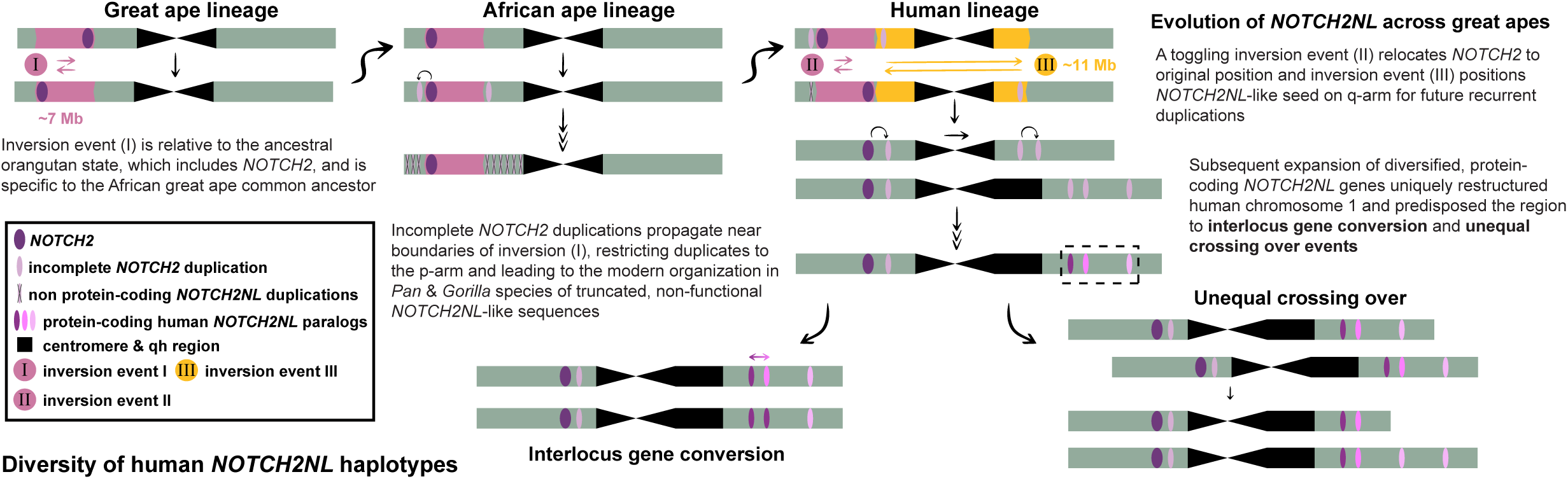
Summary of chromosome 1 rearrangement and *NOTCH2NL* evolution. *NOTCH2NL*-like genes have duplicated ancestrally and independently in three separate ape lineages, with protein-coding ability exclusively in the human lineage. Three inversion events, beginning in the African ape common ancestor, in addition to the subsequent duplication events, helped position *NOTCH2NL* to the current human genomic configuration. The organization and high homology maintained between human *NOTCH2NL* paralogs and surrounding regions promotes structural variation associated with gene conversion and unequal crossing over. See also Figures S1–S2 and Table S2.

Next, we constructed a maximum likelihood (ML) phylogeny using shared intronic sequence (intron 2) from a subset (8/29) of African ape *NOTCH2/NL* paralogs, the five human paralogs, and *NOTCH2* from Sumatran orangutan (Methods). We observe a distinct monophyletic clade populated only by the human T2T-CHM13-*NOTCH2NL* paralogs (Figure 3A, Figure S3), suggesting independent duplication or recent human-specific IGC. The topology of the tree further suggests independent expansions in the gorilla and *Pan* ape lineages, including lineage-specific expansions. In contrast, bonobo and chimpanzee share ancestral copies prior to their divergence (1-2 MYA)^3^. Using orangutan divergence and the species as an outgroup (Methods), our analysis predicts that the human lineage of *NOTCH2NL* copies emerged early in human evolution, around 4.9 MYA (4.3-5.7 MYA), after African ape speciation, and that such duplications were also occurring among the other ape lineages (albeit independently or subsequently derived from a larger initiating ancestral ape duplication). Approximately 3.0 MYA (2.2-3.7 MYA), the human-specific copies begin to diverge, distinguishing *NOTCH2NLC* from *NOTCH2NLA/B*. *NOTCH2NLA* and *NOTCH2NLB* appear to have diverged around 1.7 MYA (1.2-2.3 MYA), although once again IGC may have homogenized these loci since there is ample evidence of ongoing gene conversion (see Patterns of *NOTCH2NL* human genetic variation) in present-day humans for these two copies, which map in closest proximity to one another.

**Figure 3.**
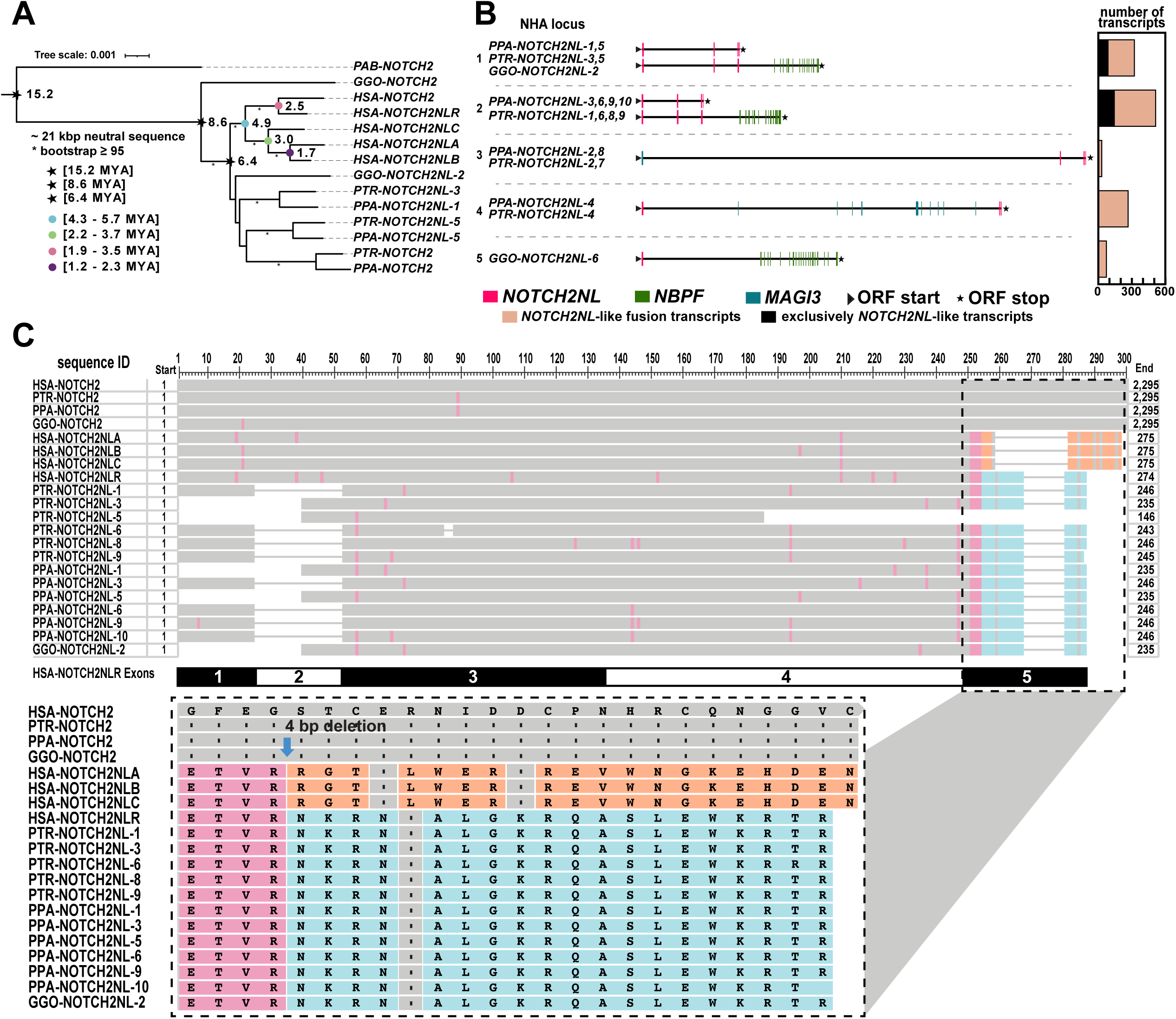
*NOTCH2/NL* duplications across multiple ape species. **A.** A maximum likelihood phylogeny based on a multiple sequence alignment (MSA) of 21 kbp of intronic *NOTCH2/NL* sequence from a subset of paralogs of five ape species, using Sumatran orangutan as an outgroup. Bootstrap support (>95%) is indicated (asterisk). Estimated divergence times of human paralogs and their confidence intervals are indicated (multicolored dots). Timings were based on human–orangutan divergence time of 15.2 MYA (Methods). **B.** Fusion transcript models expressed in different NHA grouped by species (GGO=Gorilla gorilla, PPA=Pan paniscus and PTR-Pan troglodytes; left), a schematic of the exon organization (middle) and transcript abundance (right) with the relative proportion of *NOTCH2NL*-like fusions (orange) compared to exclusive *NOTCH2NL*-like transcripts (black). These sets are representative of 20/26 *NOTCH2NL*-like loci in NHA from testis, fibroblast/lymphoblastoid cell lines, iPSCs, neuroepithelium, and neural progenitor cells. **C.** MSA of predicted protein sequences from 13/26 NHA NOTCH2NL-like loci, NOTCH2 from the NHAs, and all five NOTCH2/NL paralogs from human. Pop-out of exon 5 alignment shows that all NHAs possess the same unmodified carboxy terminus as NOTCH2NLR, which lacks a 4 bp deletion necessary for expression^1^. See also Figures S3–S5 and Table S3.

### NHA *NOTCH2NL* copies and fusion transcripts

Like humans, nearly all homologs in NHA map 11-12 kbp upstream of *NBPF* genes (25/26) (Table S3, Figure S4, Figure S5). We leveraged long-read RNA sequencing (RNA-seq) data primarily from testis and fibroblast/lymphoblastoid cell lines from chimpanzee, bonobo, and gorilla^3^ to annotate valid open reading frames (ORFs) across the majority of *NOTCH2NL*-like loci (23/26) in a set of NHA transcripts. While most transcripts predict fusions of *NOTCH2NL* with other genes (24/26), some transcripts (14/26) predict proteins most similar to human *NOTCH2NLR* sequence with ORF lengths ranging between 235-246 amino acids (AAs). The vast majority (19/26) of NHA *NOTCH2NL*-like loci create fusion transcripts where the site of fusion is at the 3’ end of *NOTCH2NL* and the 5’ end of *NBPF* (Figure 3B), which is similar to what is seen in all human paralogs (not including *NOTCH2*), but especially in human *NOTCH2NLR* (see Transcriptional expression and protein stability of *NOTCH2NL* paralogs). This included previously unreported gene fusions between *NOTCH2NL* and other genes, like *LRIG2* and *SORT1* (Table S3). All transcripts and predicted proteins similar in structure to *NOTCH2NLR* (specifically without additional gene fusion) in NHAs are distinct from human *NOTCH2NLR* because they have lost either exon 1 (containing the secretory pathway signal sequence) or exon 2 (Figure 3C). The phylogeny, different duplication architecture, and varying gene structures all support a largely independent evolutionary expansion among the great apes. Humans appear to be the only species with *NOTCH2NL* transcripts that are predicted to make a stable protein, likely because NHA copies lack the 4 bp deletion that was found to be essential for *NOTCH2NLA/B/C* protein expression^1^. We confirm this 4 bp deletion, which is after the fourth AA in the fifth exon, modifies the final 19-20 AAs of the carboxy terminus in not just a paralog-specific, but also human-specific, fashion (Figure 3C).

### Patterns of *NOTCH2NL* human genetic variation

To understand human *NOTCH2NL* genetic variation, including structural differences among human haplotypes, we initially selected 94 haploid genome assemblies recently generated by the HPRC^21^. We manually validated each assembly for these loci by assessing contiguity, annotated gaps, the presence of collapses, and verifying true SVs. Of these assemblies, 69 (73%) passed QC for sequence and structural accuracy; 54% of these genomes were of African origin while the remaining 46% were of non-African origin representing in total 14 distinct population groups (Table S4). Among these 69 genomes, we distinguished 11 different structural configurations operationally defining H1 based on the T2T-CHM13 reference configuration described above (70 haplotypes in total). Given the anticipated high degree of IGC^1,28^, we developed a tripartite workflow (Figure 4A) to assign *NOTCH2NL* identity. First, we examined the best transcript match by identifying which *NOTCH2NL* coding sequence best matches *NOTCH2NL* copies assigned in the T2T-CHM13 reference (Methods). Second, we used *NOTCH2NL* intronic sequence to construct a tree identifying a phylogenetic framework for each *NOTCH2NL* haplotype assigning different haplotypes to related clades. Third, we used the extended duplication organization as defined by the DupMasker barcode described above (Figure 1B) to examine the long-range organization of the region flanking *NOTCH2NL*. The combination of these results (Figure 4B) was used to delineate IGC events and to further define 11 distinct human haplotype configurations (H1-11) (Figure 5A).

**Figure 4.**
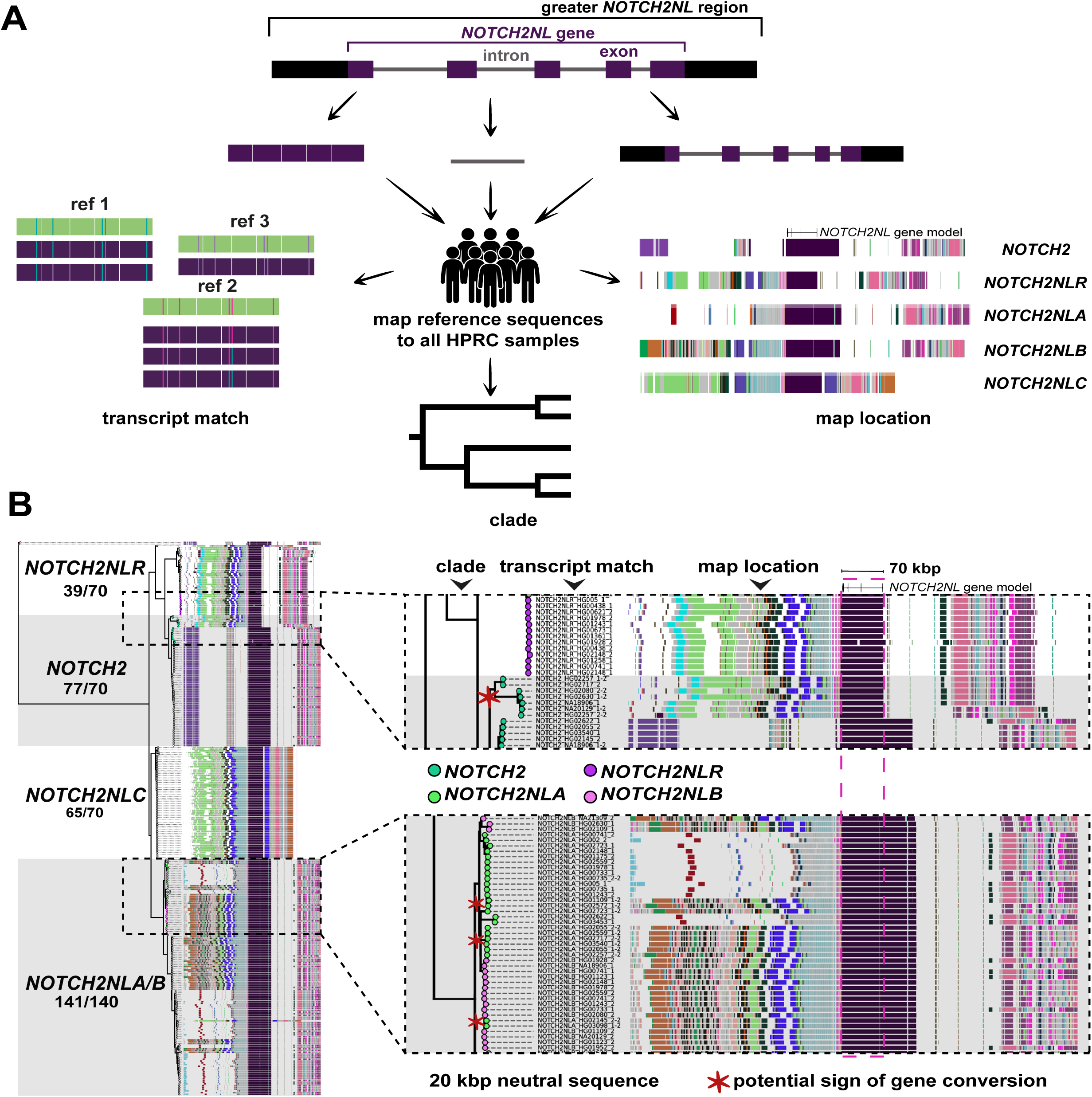
Patterns of human *NOTCH2NL* structural variation and gene conversion. **A.** Workflow to characterize *NOTCH2NL* paralog identity based on i) best transcript match (defined as the fewest mismatches with respect to T2T-CHM13 reference CDS annotation), ii) phylogenetic clade (assignment to nearest monophyletic grouping based on *NOTCH2* intronic ML tree), and iii) map location (defined here as the long-range genomic context based on DupMasker barcodes). **B.** Analysis of 70 human haplotypes depicts the clade assignment based on the phylogenetic tree, then the best transcript match, and finally the long-range duplicon organization based on the assembled HPRC genomes. Disagreements in paralog identity suggest potential gene conversion; examples marked with red asterisks. See also Figure S13 for phylogeny without gene conversion panels and Table S4.

**Figure 5.**
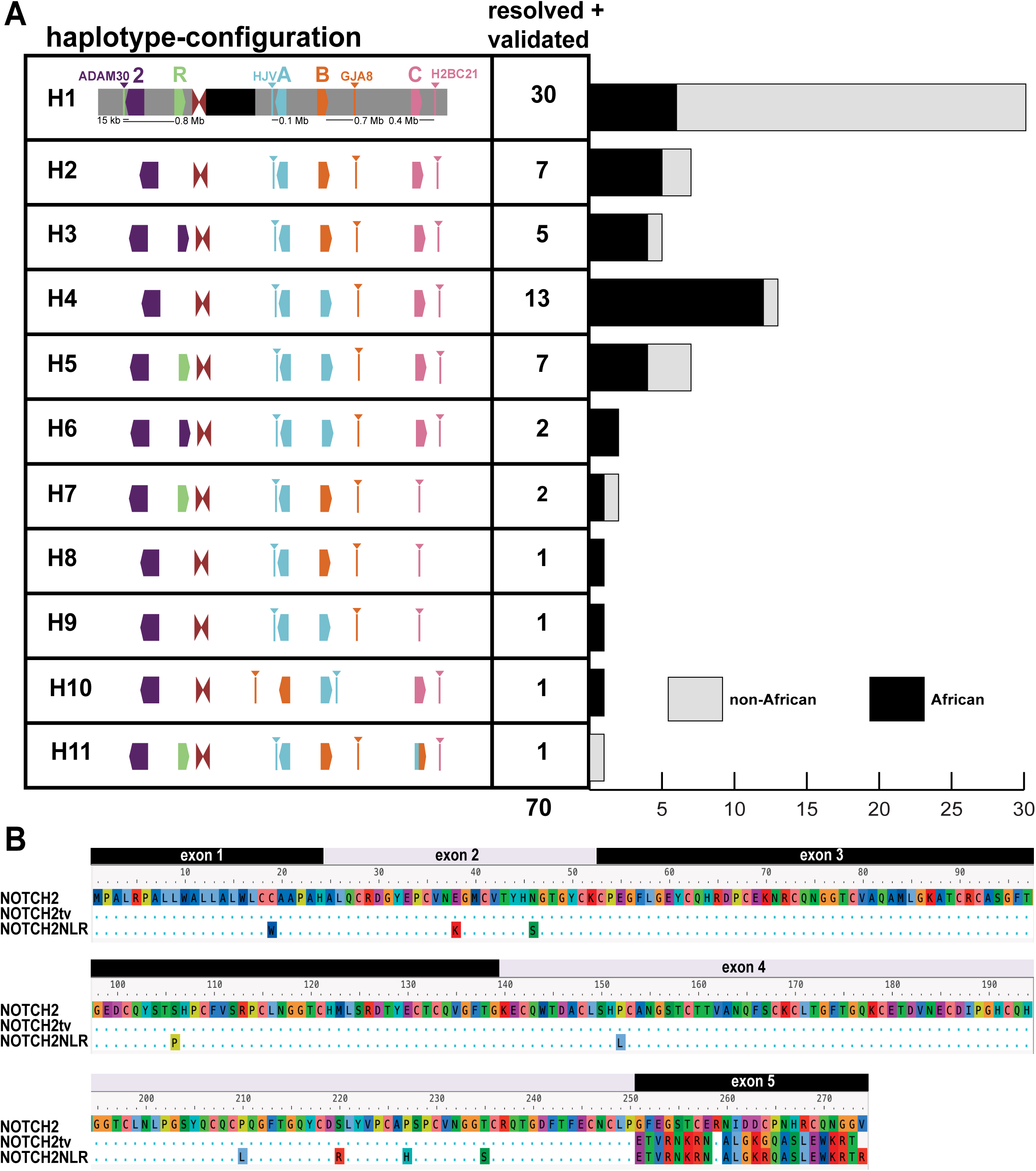
*NOTCH2NL s*tructural diversity and *NOTCH2tv.* **A.** A simplified schematic summary of the *NOTCH2NL* haplotype organization and frequency based on 69 sequence-resolved HPRC genomes and the T2T-CHM13 reference. The proportion of African and non-African samples associated with each haplotype configuration suggests some may be ancestral. **B.** Alignment of predicted AAs for the three paralogs suggests that *NOTCH2tv* arose as a result of an interlocus gene conversion (IGC) of *NOTCH2NLR* from *NOTCH2.* See also Figure S7 and Table S4.

Of the haplotype-resolved genomes, 42% share the canonical haplotype configuration (H1) observed in T2T-CHM13, thus representing the major human haplotype (Figure 5A). Among the remaining ten configurations, six are observed more than once in this subset from the HPRC (Figure 5A). Notably, the haplotype configuration currently represented in the standard human reference, GRCh38, is characterized by a nearly 2.5 Mbp inversion that reverses the orientation of *NOTCH2NLB* relative to T2T-CHM13 (Figure S6), yet it has not been observed in any other human haplotype. GRCh38, thus, either represents a minor variant or a misassembly. Our analysis indicates that the *NOTCH2NLR* pseudogene is not present in 43% (30/69) of all haploid assemblies (H2-4, H6, and H8-10). H2, H4, and H8-10 configurations strictly represent deletion events, which occur in 33% (23/69) of haploid assemblies. *NOTCH2*, is invariant with respect to copy number, however, so too is *NOTCH2NLA*, which is present in all sequenced human haplotypes but sometimes as more than one copy due to gene conversion. *NOTCH2NLC* is deleted in four haploid assemblies with three different configurations (H7-9) while in another (H11) it appears to have been converted to a *NOTCH2NLA/B* hybrid. Other than *NOTCH2NLR*, a likely pseudogene, *NOTCH2NLB* appears to be absent from human haplotypes based on sequence homology due to IGC between *NOTCH2NLA* and *NOTCH2NLB*. As a result, only 46 out of the expected 69 haplotypes actually carry it.

Overall, in this study, 33% (23/69) of haplotypes (H4, H5, H6, H9) appear to have a *NOTCH2NLB* to *NOTCH2NLA* conversion event (Figure 5A). This IGC event is shown by our workflow where both the phylogeny and duplication barcode are disrupted between *NOTCH2NLA* and *NOTCH2NLB* (Figure 4B) and has been confirmed by multiple sequence alignment (MSA) of predicted protein sequences from our samples. In addition, we observed a second gene conversion event that had not been previously characterized in-depth: direct conversion of the *NOTCH2NLR* pseudogene from the *NOTCH2* ancestral locus (Figure 4B), which is 654 kbp distant. Ten percent of haplotypes (H3 and H6) harbor a copy of the gene at this locus that resembles a truncated version of *NOTCH2* rather than *NOTCH2NLR*; this includes two H6 haplotypes that exhibit both gene conversion events (making the combined amount of IGC across haplotypes 42% instead of 43%). As a result, all eight AA changes associated with NOTCH2NLR now match the ancestral NOTCH2 (Figure 5B). When surveying gene conversion at the gene level, we see more >99% identity bins between *NOTCH2* and the gene conversion product than between the product and *NOTCH2NLR* (Figure S7). Notably, the H3 and H6 haplotypes are significantly enriched in African samples (p=0.007, Fisher’s exact test), suggesting *NOTCH2NLR* to *NOTCH2* could be an ancestral gene conversion event. Because of its sequence similarity to *NOTCH2*, we renamed this version of *NOTCH2NLR* to *NOTCH2tv* (*NOTCH2*-truncated-version), a sixth paralog in the gene family.

### Accessible chromatin architecture surrounding the *NOTCH2NL* paralogs

Having established the genetic architecture of *NOTCH2NL* paralogs and their surrounding loci, we next sought to determine how the structure of SDs influences the gene regulatory landscape surrounding *NOTCH2NL* paralogs. Gene regulatory landscapes are often defined using techniques like ATAC-seq and DNase-seq^29,30^, which can detect accessible chromatin elements. However, SDs have been historically excluded from these short-read-based techniques, as it is largely impossible to unambiguously assign short reads to large, highly identical SD regions. Fiber-seq, in contrast, is a long-read-based approach for mapping chromatin architecture^31^ and we previously demonstrated that this approach can be used to map chromatin accessibility to complex genomic regions, such as SDs^32^.

To determine whether there are any accessible chromatin elements within the vicinity of *NOTCH2NL* paralogs, we mapped CHM13 Fiber-seq data^33^ to the 300 kbp regions surrounding each *NOTCH2NL* paralog transcription start site (TSS) in T2T-CHM13 and compared the accessible chromatin maps from each paralog based on genetic synteny (Figure S8). Overall, this revealed that each *NOTCH2NL* paralog shares a promoter with accessible chromatin within CHM13 cells. Notably, each paralog shows a largely unique surrounding accessible chromatin landscape along regions that lack synteny, suggesting that non-syntenic sequence contributes to regulatory differences among the five *NOTCH2NL* paralog regions.

To further investigate the accessible chromatin landscape surrounding all the *NOTCH2NL* paralogs, including *NOTCH2tv*, in more relevant tissue, we generated Fiber-seq and long-read full-length transcript sequencing data from brain organoids of an HPRC sample that contains *NOTCH2tv* in addition to the other four copies of *NOTCH2NL* (Figure 6A-C, Methods). Specifically, lymphoblast cell lines from HG02630 were reprogrammed into iPSCs, which were then differentiated towards dorsal forebrain tissue. Such organoids of the cerebral cortex are used to model early fetal brain development, when *NOTCH2NL* is most highly expressed^1^. HG02630 has both a *NOTCH2tv* haplotype and a canonical *NOTCH2NLR* haplotype, enabling the evaluation of the gene regulatory landscape of all the *NOTCH2NL* paralogs within the same individual. Overall, like CHM13, we observe that *NOTCH2*, as well as all the *NOTCH2NL* paralogs, including *NOTCH2tv*, have accessible promoter elements that show a similar degree of chromatin accessibility (Figure 6B). In contrast to the promoter, accessible elements within the vicinity of these genes, once again, exhibit marked paralog-specific patterns both in terms of their location and accessibility.

**Figure 6.**
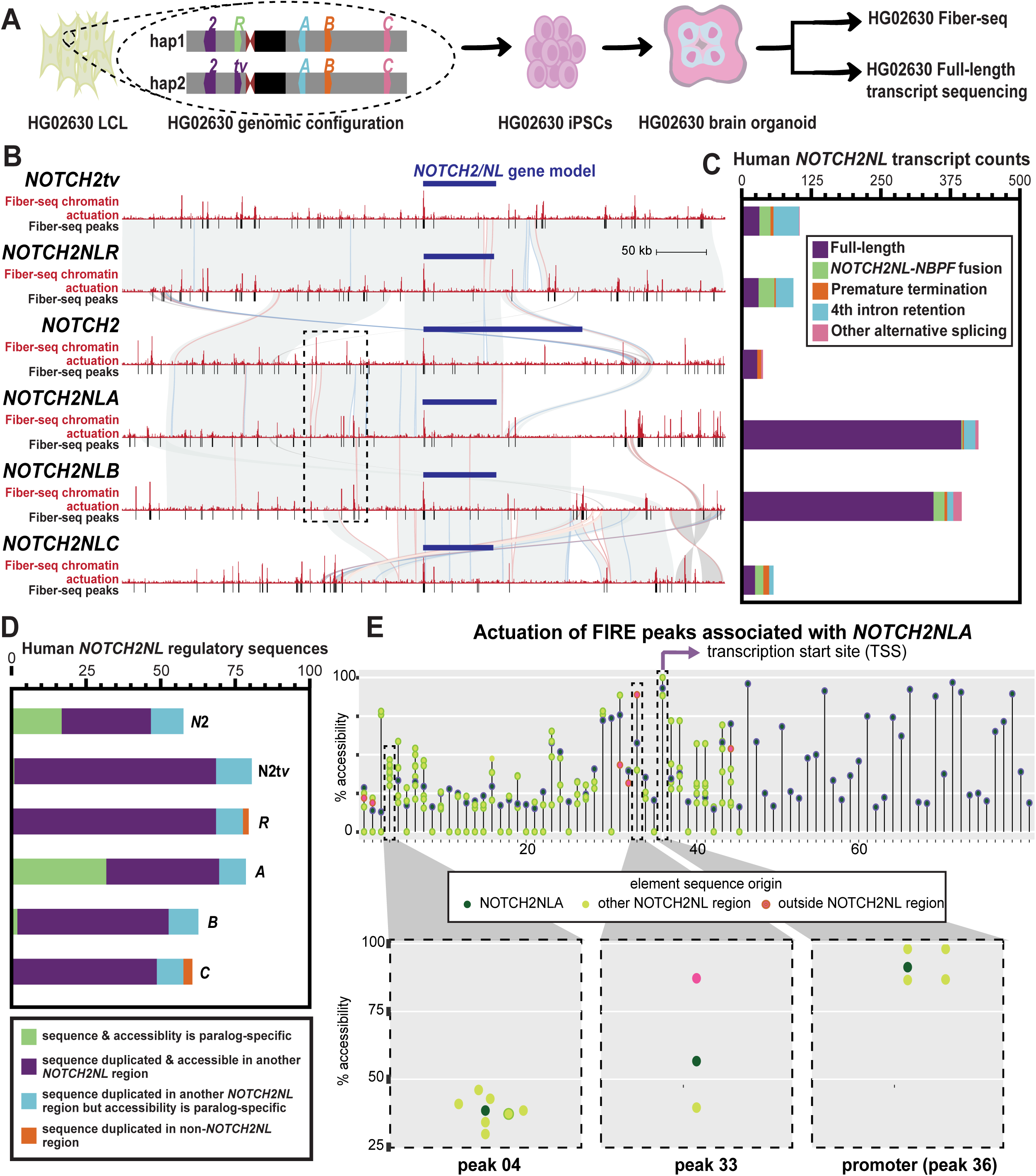
Regulatory architecture and transcription of *NOTCH2NL* in brain organoids. **A.** HG02360 was reprogrammed into iPSCs, differentiated into brain organoids, and then subjected to Fiber-seq and Iso-Seq to define putative regulatory elements and generate full-length transcripts. **B.** Fiber-seq peaks and chromatin actuation sites for each *NOTCH2/NL* paralog in the context of homology (gray), gene model, and transcription start site (TSS). Dotted black boxes are around elements specific to a region only shared across *NOTCH2*, *NOTCH2NLA*, and *NOTCH2NLB*; though the underlying sequence is nearly identical, we show paralog-specific actuation signals. **C.** The absolute abundance of full-length transcripts compared to other premature termination, fusion, and intron retention products in brain organoids. **D.** Bar graph showing categorization of accessible elements surrounding each *NOTCH2NL* paralog based on the presence of duplicate sequence and accessibility at that sequence on the different paralogs. Note that *NOTCH2* and *NOTCH2NLA* have the greatest proportion of paralog-specific sites (dark green). **E.** Percent actuation of each accessible regulatory element surrounding the *NOTCH2NLA* paralog (dark green) as well as the percent actuation of duplicate sequences for each element that are present surrounding other *NOTCH2NL* paralogs (light green) or outside of the *NOTCH2NL* paralogs (pink). See also Figures S9–S11.

Comparison of these accessible Fiber-seq-based chromatin maps (Methods) with underlying synteny maps reveals that nearly 87% of all accessible chromatin elements within 300 kbp of the *NOTCH2NL* TSSs share duplicated sequence with at least one other *NOTCH2NL* paralog region. This suggests prevalent reuse of genomic sequence with putative gene regulatory potential across these SDs (Figure 6D). Furthermore, 84% of those elements near multiple *NOTCH2NL* paralogs also show some level of accessible chromatin at more than one of their paralogous sites, demonstrating that by and large, these elements retain their ability to form accessible chromatin when rearranged in different genomic positions. Overall, the accessible chromatin landscape surrounding each *NOTCH2NL* paralog appears predominantly populated by these multi-paralog accessible chromatin elements. However, these paralogs often diverged in their magnitude of chromatin accessibility (Figure 6B,E) as well as their rearrangement relative to the *NOTCH2NL* promoters, indicating that position effects within these different SDs may impact these putative regulatory elements in a quantitative manner as opposed to simply abrogating their chromatin accessibility.

We also observe that 10% of all accessible chromatin elements within 300 kbp of the *NOTCH2/NL* TSSs map to duplicated sequence on a different chromosome. Five elements exclusively share duplicated sequence with regions on a different chromosome (about 1% of all elements) (Figure 6D). This suggests that the creation of the *NOTCH2NL* SDs was associated with the potential repurposing of accessible chromatin elements from elsewhere in the genome.

In total, we find that 12% of all accessible chromatin elements surrounding *NOTCH2/NL* paralogs are specific to only one paralog. These paralog-specific unique elements are concentrated in the *NOTCH2* and *NOTCH2NLA* regions. Notably, these paralogs are the most fixed for copy number variation within the human population.

To further refine the scope of regulatory elements more directly associated with each *NOTCH2NL* paralog, or at least their respective promoters, we applied FiberFold^34^ to our brain organoid data to predict topologically associated domains (TADs) in *NOTCH2NL* paralog regions in a haplotype-aware manner. FiberFold predicts each *NOTCH2NL* paralog occupies a distinct 3D genomic environment despite their highly identical underlying sequence, with each *NOTCH2NL* TSS surrounded by a unique 3D landscape (Figure S9A). These paralog-specific 3D landscape patterns are largely consistent across haplotypes, with differences between paralogs more pronounced than differences between haplotypes (Figure S9B). Consistent with this, we also find that chromatin actuation differences between paralogs are more pronounced than differences between haplotypes (Figure S10), despite the high sequence identity between paralogs. Limiting this analysis to accessible chromatin elements that fall within the predicted TADs for each of the *NOTCH2NL* paralogs confirmed the above findings (Figure S11) and exposed a cluster of elements 63 kbp upstream of the *NOTCH2*, *NOTCH2NLA*, and *NOTCH2NLB* TSSs, which are selectively active on the *NOTCH2NLA/B* SDs but inactive on the *NOTCH2* SD (Figure S9C). Overall, this analysis uncovers marked SD-specific accessible chromatin architectures and nominates specific elements that may be resulting in the paralog-specific regulation of the *NOTCH2NL* genes. However, further experiments will be needed to characterize the functional role of elements within TAD boundaries.

### Transcriptional expression and protein stability of *NOTCH2NL* paralogs

We found distinct differences within the transcript abundance of each of the *NOTCH2/NL* paralogs (Figure 6C), indicating that these paralog-specific accessible chromatin elements may be creating unique gene regulatory environments for each of the *NOTCH2/NL* paralogs. Specifically, *NOTCH2NLA* and *NOTCH2NLB* had ∼3-fold higher steady-state transcript abundance than the other *NOTCH2/NL* paralogs. Ancestral *NOTCH2* has a low abundance of full-length transcripts; however, *NOTCH2* also has very few other transcript types, including fusions. In fact, NHAs appear to show higher levels of fusion transcripts with the ancestor than seen in humans (Figures 3B, 6C). Furthermore, we observed that although the promoter and transcript sequence of *NOTCH2tv* mirrors that of *NOTCH2*, the transcript abundance and composition of *NOTCH2tv* appeared to mirror most closely that of *NOTCH2NLR*. Specifically, *NOTCH2tv* and *NOTCH2NLR* had 102 and 91 transcripts, respectively. However, only ∼30% of these transcripts represented canonical full-length transcripts, with the majority arising from fusion transcripts, and an incorrectly spliced exon. Surprisingly, the fusion transcripts of all *NOTCH2NL* copies do maintain ORFs predicted to be 1179-1574 AA long. Overall, this indicates that despite the transcript identity of *NOTCH2tv* matching the first four exons of *NOTCH2*, the surrounding gene regulatory architecture in fact mirrors that of *NOTCH2NLR*, potentially impacting the overall function of *NOTCH2tv.* This is likely a result of the gene conversion event being bounded by a 75 kbp syntenic block between *NOTCH2* and *NOTCH2NLR* that spans from just upstream of their promoters to their fourth introns.

Although the gene conversion event results in NOTCH2tv adopting the exact same protein sequence as NOTCH2 for its first 250 AAs, NOTCH2tv ends in a distinct 23 AA sequence arising from its terminal fifth exon. NOTCH2NLR was previously shown to form an unstable protein product, which is thought to be driven by its carboxy-terminal sequence. As such, we sought to evaluate whether NOTCH2tv similarly forms an unstable protein product, as its C-terminal sequence shares 91% AA similarity to NOTCH2NLR. We transfected HEK293 cells with a constitutive reporter system containing either *NOTCH2tv*, *NOTCH2NLR*, *NOTCH2NLB*, or a negative control and demonstrated that despite the ability of *NOTCH2tv*, *NOTCH2NLR*, and *NOTCH2NLB* to produce sufficient transcripts in this reporter, only *NOTCH2NLB* resulted in a stable protein product (Figure S12). Together, these data indicate that although gene conversion has generated a paralog of *NOTCH2NL* that contains sequence and promoter features consistent with *NOTCH2*, this paralog retains the overall gene regulatory architecture and transcript patterns of *NOTCH2NLR* and is similarly unable to form a stable protein product and, thus, likely represents a pseudogene.

## DISCUSSION

The rapid expansion of interspersed SDs in the ancestral genome of African apes around 8-15 MYA^35^ provided the substrate for the human genome to evolve both ape and species-specific genes. *NOTCH2NL* is one of at least five human- and ape-specific SD gene families that have been implicated in expansion of the human frontal cortex. This includes genes associated with delayed maturation of synapses and increasing synaptic density (*SRGAP2C*)^36,37^, genes like *NOTCH2NL*^1,2^ implicated in cortical progenitor self-renewal, and genes directly promoting cortical and basal progenitor amplification (*TBC1D3*, *ARHGAP11B*, and *CROCCP2*)^38,39,40,41^. Like *NOTCH2NL* it is noteworthy that most of these human-specific gene innovations originated from an incomplete duplication that truncated the ancestral gene model, leading to human-specific isoforms. In fact, the incomplete duplication appears to have been a first critical step in either neofunctionalization (*ARHGAP11B*) or dominant negative effects (*SRGAP2C*, *NOTCH2NL*) where shorter, derived proteins either interfere or modulate ancestral protein function through protein-protein interactions^39,40,36,42,1,2,37^.

We demonstrate that *NOTCH2NL*, like other recently characterized primate gene families, likely independently expanded in human, chimpanzee, and gorilla (*TBC1D3*, *LRRC37*, and *NPIP*)^43,44,45,46^. The basis for this recurrence or genomic instability is unknown, but it is interesting that most *NOTCH2NL* ape copies are also associated with the *NBPF* duplicon—an association that is postulated to have co-evolved both in terms of structure and transcriptional regulation, especially in humans^16^. *NBPF* is one of about a dozen core duplicons (along with *TBC1D3*, *LRRC37*, and *NPIP*) implicated as a potential driver of interspersed SDs in the primate lineage^13,14,15^. Of note, a comparative analysis of *NBPF* associated with *NOTCH2NL* reveals species-specific expansions of different portions of the *NBPF* DUF1220 in different apes (Figure S5), so it is possible that *NBPF* plays a more general role in gene innovation in ape species other than human.

Notwithstanding this proclivity to duplicate in the common ancestor of great apes, the apparently functional human copies of *NOTCH2NL* arose much later in human evolution. We estimate that the human-specific expansions (or IGC events) occurred around 4.9 MYA and diversified over a range of 3.0-1.7 MYA (Figure 3A). It is worth noting that other duplicate gene families implicated in the expansion of the human frontal cortex (*SRGAP2C* and *TBC1D3*) show similar evolutionary trajectories beginning to emerge 2-3 MYA^47,43^. This is significant in the context of fossil record evidence, which suggests divergence of the genus *Homo* from *Australopithecus* ∼2 MYA and a subsequent initial increase in archaic hominin cranial volume. There is also evidence of subsequent increases in cranial volume taking place between 2.0-1.5 MYA consistent with the diversification of *NOTCH2NL* genes in humans^48^.

Among human haplotypes, we report polarized signals of gene conversion, which seems especially significant in the case of *NOTCH2NLA*, the only paralog present in all assemblies and that has expanded, seemingly at the expense of *NOTCH2NLB*. This may be a consequence of either an unknown mechanistic or selective bias favoring *NOTCH2NLA* as a donor. In Fiddes et al. (2018)^1^, it was also postulated that having a combined dosage of A/B was more important than having two of each paralog. Like *SRGAP2C*^47^, *NOTCH2NLA* represents the most fixed paralog suggesting functional constraint. Gene birth can be accomplished through duplication^49^, but there is a gap in the research on how IGC may influence this process. When first investigating the understudied paralog *NOTCH2tv*, we hypothesized it was a case of gene conversion reviving a nonfunctional gene since so far *NOTCH2tv* has acquired a promoter and 4-exon N-terminus identical to *NOTCH2*. This in theory could enable *NOTCH2tv* to regulate expression similar to *NOTCH2*^23^. However, whereas *NOTCH2NLA/B/C* contain a protein-stabilizing 4 bp deletion in their terminal exon (Figure 3C), neither *NOTCH2NLR* nor *NOTCH2tv* has this same 4 bp deletion in its terminal exon (Figure 5B). Consistent with this, we found that like *NOTCH2NLR*, *NOTCH2tv* does not produce a stable protein (Figure S12) and, thus, is not a fully functional paralog.

By leveraging Fiber-seq and FiberFold, we identify marked paralog-specific gene regulatory patterns surrounding each *NOTCH2NL* SD and paralog. Overall, we find that the different *NOTCH2NL* regions frequently retain duplicated sequences that encompass an accessible chromatin element on at least one paralog region. However, ∼14% of elements present within these duplicated sequences exclusively show chromatin accessibility in only one paralog. Furthermore, even for those elements that do show some chromatin accessibility across two or more duplicate sequences, we find that the degree of chromatin accessibility can vary quite substantially between the two duplicates (Figure 6E). This suggests that putative regulatory elements within SDs are being subjected to positional effects, with the predominant effect being quantitative differences in chromatin accessibility as opposed to drastic changes to on/off actuation.

In summary, we hypothesize that the dramatic restructuring of the *NOTCH2NL* loci during human evolution led to the only ape lineage with protein-coding copies. This was made possible by a dynamic set of large- and small-scale changes associated with NAHR, recurrent duplications/deletions^50^, and IGC^25^. Many genes embedded in these regions, including *NOTCH2NL*, are associated with neurologic and developmental phenotypes, including copy number variation syndromes, such as 1q21.1 distal duplication/deletion syndrome^11^ or TAR syndrome^9^. The fact that this region is among the most frequently rearranged regions of the human genome^51^ is a testament to the evolutionary instability that continues to persist in the human population. It is well established that the presence of these human-specific duplications has phenotypic consequences associated with recurrent rearrangement and developmental delay^10,11^). Consistent with the core duplicon hypothesis^52^ and population studies that suggest large duplications should be under purifying selection unless some other type of selective force is acting^53,54,55,56^, the mutational lability of chromosome 1q21.1 and the emergence of *NOTCH2NL* genes likely represents a significant trade-off between positive and negative selection during human evolution. In the case of *NOTCH2NL*, we hypothesize that the benefits of expanding the cortex must have outweighed the mutational burden of increasing the proportion of high-identity duplicated sequences in the genome. Our findings suggest that this trade-off is still ongoing. The biased gene conversion that is potentially driving the fixation of *NOTCH2NLA* as well as the high level of fourth intron retention transcripts of *NOTCH2tv* and *NOTCH2NLR* may be examples of ongoing evolution in this gene family, such as specification of the functional NOTCH2NL protein and potential acquisition of a novel carboxy terminus.

### Limitations of the study

#### Gene conversion biases

Gene conversion exists at varying degrees throughout this locus, even in T2T-CHM13, which is used as a reference and may be a source of error for analysis of human genetic variation. Although we attempted to identify the largest gene conversion events, both reference effects and smaller events may remain undetected. IGC obscures the timing of duplication events and signals of selection. Additionally, homogenization between highly similar loci where only a few SNV differences distinguishes paralogous segments complicates breakpoint definition.

#### Phylogeny and duplication timing

Although we assess robustness of the phylogenies using bootstrap support, several sources of error remain. The largest source is the timing of speciation events used to calibrate the molecular clock, which are estimates themselves. As noted previously, IGC is also a concern, as any duplication timing estimates we report may capture time since gene conversion versus time since the original duplication event.

#### Gene regulation and Fiber-seq

Fiber-seq is a measure of chromatin accessibility, which is only a proxy for noncoding regulatory regions; it does not distinguish between element types, such as repressors and enhancers, nor address which genes these elements would be regulating. Further work is needed to resolve how paralog-specific regulatory architectures diverge relative to SD sequence identity. Although FiberFold will generate predictions of which genes may be regulated by these accessible chromatin peaks, further efforts are needed to functionally characterize these predictions.

#### Iso-Seq analyses

Iso-Seq is a measure of steady state transcripts, which is influenced by both the potency of regulatory elements as well as the stability of the full-length transcript. Given the marked divergence between *NOTCH2NL* paralogs in terms of the surrounding regulatory elements as well as their 3’ UTR structure, it is anticipated that both of these are contributing to the final steady state measurements of transcript abundance.

## RESOURCE AVAILABILITY

### Lead Contact

Requests for further information and resources should be directed to and will be fulfilled by the lead contact, Evan Eichler.

### Materials Availability

DNA sequence of plasmids generated in this study for NOTCH2NL protein expression have been deposited to GitHub (https://github.com/tdreal/NOTCH2NL-0325/tree/main) and Zenodo (https://zenodo.org/records/15022214).

### Data and Code Availability

PacBio HiFi Fiber-seq and Kinnex full-length Iso-Seq from HG02630 brain organoids generated for this study have been made available on NCBI with the BioProject ID: PRJNA1236375. Original western blot images are deposited to GitHub (https://github.com/tdreal/NOTCH2NL-0325/tree/main) and Zenodo (https://zenodo.org/records/15022214). There is no original code reported by this study. Any additional information required to reanalyze the data reported in this paper is available from the lead contact upon request.

## ACKNOWLEDGMENTS

We would like to thank T. Brown for editing this manuscript. We would like to thank Nicolas Altemose at Stanford University for additional support in generating and validating the FiberFold tool. We would like to thank the UW ISCRM facility for performing HG02630 iPSC generation. We would also like to thank the HPRC and Primate T2T Consortium for providing numerous high-quality assemblies for analysis. A.B.S. holds a Career Award for Medical Scientists from the Burroughs Wellcome Fund and is a Pew Biomedical Scholar. This work was supported, in part, by US National Institutes of Health (NIH) grants R01MH120295 to S.R.S., 1DP5OD029630 and 1U01HG013744 to A.B.S., and R01HG010169 and R01HG002385 to E.E.E. E.E.E. is an investigator of the Howard Hughes Medical Institute.

This article is subject to HHMI’s Open Access to Publications policy. HHMI lab heads have previously granted a nonexclusive CC BY 4.0 license to the public and a sublicensable license to HHMI in their research articles. Pursuant to those licenses, the author-accepted manuscript of this article can be made freely available under a CC BY 4.0 license immediately upon publication.

## AUTHOR CONTRIBUTIONS

This work was conceptualized by T.D.R., A.B.S., and E.E.E. Chromosome 1 visual alignment of *NOTCH2NL* region between human and NHP and NHA *NOTCH2NL* SD analyses were done by D.Y. with additional visualization and genomic rearrangement analysis by F.A. Annotation of *NOTCH2NL* gene fusions in NHA and *NBPF* DUF1220 domain sequence analysis was performed by P.H., assisted by M.D. A subset of HPRC assemblies were re-run with Verkko by X.G. NucFreq validation of HPRC assemblies was processed by A.N.R. Human *NOTCH2NL* phylogeny was constructed and visualized by T.D.R. with support from P.C.D. Human SD and IGC analyses were performed by T.D.R. and assisted by M.R.V. Dorsal forebrain organoid generation of HG02630 was performed by I.P. FiberFold TAD prediction analysis was done by D.D. NOTCH2NL protein-expression cloning, tissue culture, and western blot experiments were done by G.J.M., O.G.P., and B.J.M. Genomic and transcriptomic data for this study were generated by K.M.M., K.H., and J.E.R. Data were processed by K.M.M., S.J.N., and A.E.S-C. The manuscript was written by T.D.R., P.H., F.A., D.Y., I.P., D.D., B.J.M., A.B.S., and E.E.E. All additional analyses not specifically listed in this section were done by T.D.R. Project was advised by B.P., S.R.S., A.B.S., and E.E.E.

## DECLARATION OF INTERESTS

E.E.E. is a scientific advisory board (SAB) member of Variant Bio, Inc. A.B.S. is a co-inventor on a patent relating to the Fiber-seq method (US17/995,058). All other authors declare no competing interests.

## STAR⍰METHODS

### Key resources table

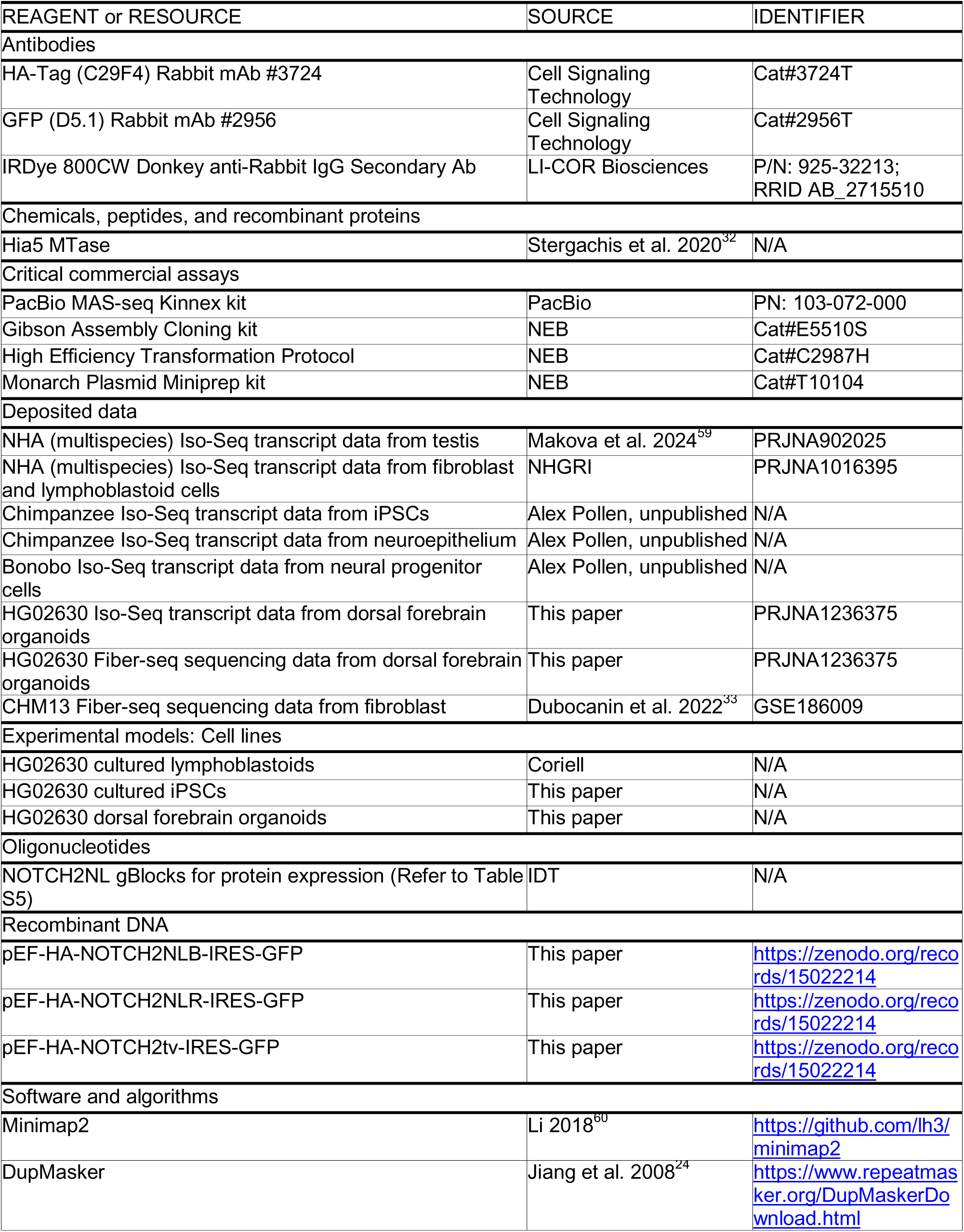
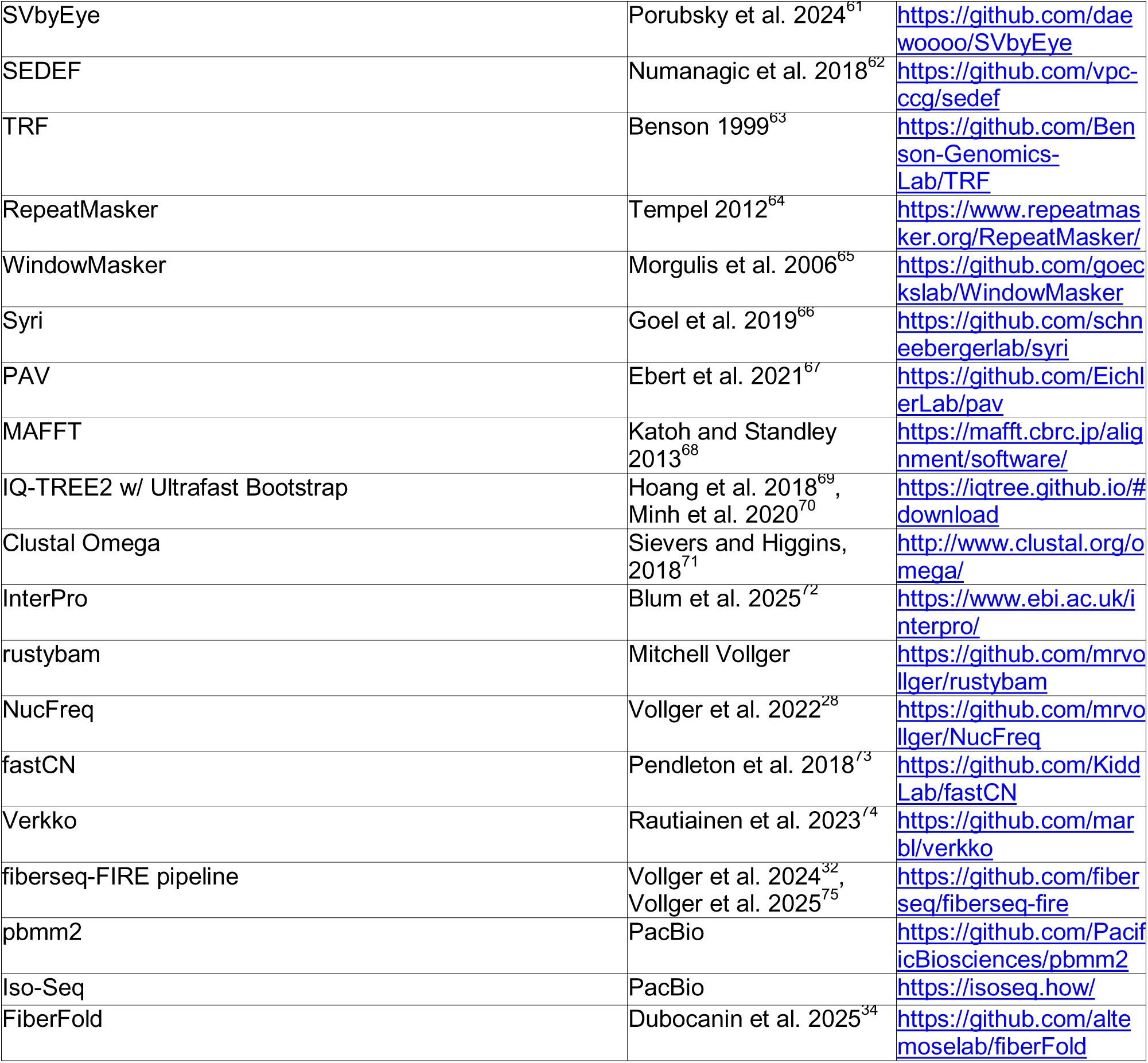

## METHOD DETAILS

### *NOTCH2NL*-CHM13 region identity matrix

*NOTCH2NL* paralog sequences from T2T-CHM13 V2.0^58^ (http://genome.ucsc.edu) plus 1 Mbp surrounding each gene were all aligned to each other simultaneously and allowing for secondary alignments (paralog pairs with less than 1 Mbp between them had overlapping sequence removed), using minimap2^60^ and the parameters

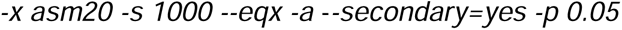

The longest nonoverlapping stretch of syntenic sequence that can be aligned between each paralog pair is represented in the matrix. The percent identity shared between each paralog pair is calculated by the number of base matches.

### *NOTCH2NL*-CHM13 region duplication barcodes

The duplicon locus barcoding is a computational approach that can readily be used to assess the content and organization of the SDs for any given paralogous locus. It takes advantage of the fact that all SDs can be decomposed into smaller evolutionary units that have accumulated into larger higher-order structures based on their juxtaposition and accumulation over time^13^. Based on this repeat graph annotation, the tool DupMasker^24^ will encode any SD region into these evolutionary subunits—the orientation and juxtaposition of each becomes a unique identifier that can be color-coded to create a visual track of the pattern of duplications in a given region. Even though *NOTCH2NL* paralogs are in SD regions that share high identity, the accumulation pattern is effectively unique allowing assignment to specific copies based on the majority-rule consensus for this DupMasker “barcode”.

In this study *NOTCH2NL* paralog sequences plus 1 Mbp surrounding each gene were used as input for DupMasker. The .duplicons output file was processed into a .txt file to create visual tracks of duplicon barcodes for each region.

### Visual alignment of *NOTCH2NL*-CHM13 regions

*NOTCH2NL* paralog sequences plus 1 Mbp surrounding each gene were aligned to each other one by one and allowed for secondary alignments, using minimap2 parameters

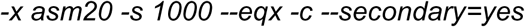

The output of each gene pair alignment was compiled into a single output .paf file that was used as input for SVbyEye^61^. The ladder plot alignment figure was visualized using the plotAVA function. Each *NOTCH2NL* region’s corresponding duplication barcodes are included as an additional annotation track. The output .paf CIGAR string was also used to return a table of insertion/deletion SVs for each paralog.

### Chromosome 1 visual alignment of *NOTCH2NL* region between human and NHPs

NHP assemblies used in this study from previous research can be found by the following GenBank accession IDs^3,20^: GCA_028858775.2 (chimpanzee), GCA_029289425.2 (bonobo), GCA_029281585.2 (gorilla), GCA_028885625.2 and GCA_028885655.2 (Bornean and Sumatran orangutans) and GCA_030222085.1 (macaque). Homologous sequence of the *NOTCH2NL* region in human (chr1:110,000,000-160,00,000) was identified by aligning human sequence against each of the NHP genomes. The alignment was performed using minimap2 with the parameters

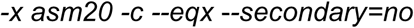

Alignment blocks equal to or larger than 10 kbp were retained. After locating the corresponding sequences of NHPs, the alignment was performed allowing for secondary alignment using the parameter

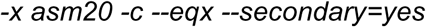

progressively in the order from human to macaque. The alignment was visualized using SVbyEye. Similar alignments between T2T-CHM13 and the previous reference genome, GRCh38, were performed using the same command (Figure S6).

### NHA SD and *NOTCH2NL* analysis

SD tracks generated by Yoo et al. 2025^3^ were used. Briefly, the SD track was annotated using SEDEF (v1.1)^62^, after masking the repeats using TRF (v4.1.0)^63^, RepeatMasker (v4.1.5)^64^, and WindowMasker (v2.2.22)^65^. The SDs were filtered for length >1 kbp, pairwise sequence identity >90%, and satellite content <70%.

### Simulation test for enrichment of SVs and SDs at inversion breakpoints

Coordinates of inversions and SVs identified by a previous study were utilized^3^, which used Syri (v1.6.3)^66^ and PAV (v2.3.2)^67^ pipelines. We first quantified SVs/SDs at the inversion breakpoints found within 110-160 Mbp regions by intersecting 100 kbp of inversion coordinates with SVs/SDs to get observed bp of SVs/SDs at the inversion breakpoints. To test enrichment of variants at the inversion breakpoints, we generated random distribution of SVs/SDs by shuffling the breakpoints, while restricting the region to 110-160 Mbp, and excluding centromere satellites. The null statistics distribution was generated by quantifying the SVs/SDs at the randomly shuffled regions. The one-sided simulation p-value was computed by quantifying the proportion of null distribution that show more extreme statistics (greater than observed SVs/SDs).

### *NOTCH2NL* NHA phylogeny and duplication timing

Multiple intronic sequences of *NOTCH2* were mapped to each NHA primate assembly using minimap2. The coordinates of all the *NOTCH2NL*-like regions found were used to pull out the sequence and construct a multiple sequence alignment (MSA) using MAFFT^68^ with the parameters

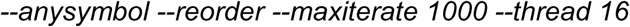

The MSA was then processed with IQ-TREE2, an ML phylogeny building program, and bootstrapped using the ultrafast bootstrap^69,70^. IQTREE estimated phylogenetic dating using LSD2 to build a time tree based on orangutan–human divergence time being 15.2 MYA, gorilla–human divergence time being 8.6 MYA, and chimpanzee–human divergence being 6.4 MYA. IQ-TREE2 was run with the parameters

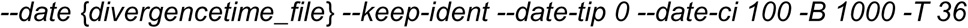

The most robustly bootstrapped tree was used in the main text.

### Annotation of *NOTCH2NL* gene fusions in NHA

NCBI RefSeq and CAT2.0. annotations from the T2T NHA genomes were used to study copies of *NOTCH2NL*-like genes across the NHA assemblies. First, the genomic annotations for any gene from the *NOTCH2NL* family were extracted from both these annotation sets. Next, any copies that were missed by either of the two methods were extracted with BLAT, using the genomic sequence of the first four exons of *NOTCH2* on the T2T NHA genomes. The full set of copies were then manually investigated on the UCSC Genome Browser with the added NHA Iso-Seq transcript data from testis, fibroblast/lymphoblastoid cell lines (available under NCBI BioProject IDs: PRJNA902025 and PRJNA1016395), as well as iPSCs, neuroepithelium, and neural progenitor cells (Pollen Lab, unpublished). The exact boundaries of the genes and fusions were determined. Blat was used to describe the exon structures of these fused genes. The predicted proteins for these copies were aligned using Clustal Omega^71^ and visualized using the NCBI Multiple Sequence Alignment Viewer (1.25.3).

### NBPF DUF1220 domain sequence analysis

The NBPF protein sequence for all the gene copies downstream of *NOTCH2NL* in human and NHA were extracted and InterPro^72^ was run to annotate all protein domains on the sequences. DUF1220 domains were then extracted and matched against the canonical sequences (UniProt) for HLS1, HLS2, HLS3, CON1, CON2, and CON3 using BLAT.

### Assembly validation of *NOTCH2NL* regions in HPRC assemblies

Except for T2T-CHM13, the human assemblies used in this study were originally released as a part of HPRC year 1^21^ without complete validation of every region. These assemblies are available under BioProject ID: PRJNA730822. To validate our set of assemblies, we first confirmed the correct assembly of contiguous sequence between *NOTCH2NL* paralogs on the same chromosome arms. This was done by evaluating the number of copies in each assembly and how many contigs they covered using rustybam (mrvollger.github.io/rustybam/). Assemblies with copies across three or more contigs were removed and those with no more than two contigs (allowing for gaps across the centromere) were assessed with NucFreq^28^. Assemblies with collapses (excess of secondary bases in a single haplotype assembly) within *NOTCH2NL*, collapses greater than 10 kbp outside *NOTCH2NL*, gaps (low-quality, N-based sequence) within *NOTCH2NL*, and gaps greater than 10 kbp outside *NOTCH2NL* over the 1 Mbp region surrounding each *NOTCH2NL* paralog were considered incorrectly assembled. Finally, assemblies were also removed from the completed set if they were missing specific unique gene marks outside *NOTCH2NL* SD regions, which are used as anchors for rearrangement boundaries. Deletions were further validated using fastCN^73^ read depth. A subset of incorrectly assembled samples was attempted to be rescued using Verkko (1.1 and 1.2)^74^ and added to the validated set: 69/94 haploid assemblies in total were correctly assembled and validated.

### Determining *NOTCH2NL* identity and uncovering IGC in HPRC assemblies

To determine *NOTCH2NL* identity we took a three-pronged approach: (1) identifying which T2T-CHM13 *NOTCH2NL* CDS reference or ‘transcript’ best matches the *NOTCH2NL* sequence being queried using BLAT; (2) identifying the phylogenetic clade the *NOTCH2NL* sequence being queried best groups with—sequences from intron 2 of *NOTCH2/NL* from all samples queried were used to construct an MSA using MAFFT, from which an ML phylogeny was built using IQ-TREE2; and (3) identifying the mapping location of the *NOTCH2NL* sequence being queried using the greater *NOTCH2NL* region DupMasker barcode—*NOTCH2NL* intron 2 sequences plus 1 Mbp surrounding each intron were used as input for DupMasker and duplicon barcodes for each sequence were added to the tree. IGC events were first found through conflicts between the coding sequence identity, clade grouping, and/or mapping location. A final visualization of protein-coding changes caused by gene conversion can be confirmed using MSA software.

### iPSC generation of HG02630

iPSCs were generated from HG02630 cultured lymphoblastoid cells obtained from Coriell using the method described in Vollger et al. (2025)^75^: StemFlex media (Thermo Fisher Scientific, A3349401) on Matrigel (Corning, CLS354277)-coated plates at 37 °C in 5% CO_2_ was used to maintain iPSCs and ReLeSR (STEMCELL Technologies, 100-0484) was used for passage of cell colonies.

### Making brain organoids of HG02630

HG02630 iPSC line maintenance and cerebral organoid generation until day 21 were done using the methods described in Seiler et al. (2022)^76^. To achieve uniform basal ECM coating on days 6 and 7, a combination of 0.2% alginate and 0.6mg/ml Geltrex LDEV-Free Reduced Growth Factor Basement Membrane Matrix (ThermoFisher) coating was used (Hoffman et al., in preparation), and the alginate was crosslinked with CaCl2.

On day 21, the organoids were broken apart using Trypsin-EDTA (0.25%), placed in the centrifuge at 250g for 5 minutes, resuspended in 0.5 ml of PBS, and centrifuged for 5 more minutes at 250g. The supernatant was aspirated, and the cells were resuspended in 180ul of Buffer A (components missing). The sample was transferred to a PCR tube, and 180ul of 2X lysis buffer was added. Cells were spun at 350g for 5 minutes, after which the supernatant was removed. The remaining nuclei pellets were resuspended in (Buffer A, 32mM SAM, Hia4 (200U/ul)) at 25°C for 10 min. Finally, 9ul of 1% SDS was added to the sample and transferred to 1.5ml tubes using wide-bore pipette tips.

### Fiber-seq and identification of chromatin accessibility in HG02630 brain organoids

PacBio HiFi Fiber-seq data were generated from HG02630 brain organoid nuclei pellets treated with Hia5 enzyme using the method described in Vollger et al. 2025^75^. The data were analyzed using the fiberseq-FIRE pipeline^32,75^ (https://github.com/fiberseq/fiberseq-fire), which identifies single-molecule sites of chromatin actuation as well as peaks of chromatin actuation with a false discovery rate 5% threshold. Percent actuation was calculated as the percentage of fibers mapping to a given location that were classified as having a Fiber-seq Inferred Regulatory Element (FIRE).

For proof-of-concept unique mapping at this locus before HG02630 data was generated, previously published T2T-CHM13 Fiber-seq data was mapped to the NOTCH2NL region (Figure S8).

### Long-read RNA-seq in HG02630 brain organoids

PacBio MAS-seq^77^ (PN: 103-072-000) data were generated from HG02630 brain organoid nuclei pellets using the method described in the RNA preparation section of the Methods in Vollger et al. (2025)^75^. The data were processed and mapped to the HG02630 diploid assembly using pbmm2 (https://github.com/PacificBiosciences/pbmm2) and isoforms were defined using the Iso-Seq pipeline (https://isoseq.how/) and annotated using Pigeon.

### Protein expression of NOTCH2tv in HEK293 cells

gBlocks were designed to contain an HA tag, *NOTCH2NL* CDS, IRES sequence, and *E-GFP* CDS (Table S5) and ordered using IDT (https://www.idtdna.com/page). Gibson Assembly (NEB, E5510) was used to clone constructs into a pEF-GFP vector. Each vector construct was cloned using NEB High Efficiency Transformation Protocol (C2987H/C2987I) with NEB 5-alpha Competent E. coli. Colonies were picked, inoculated, and plasmid DNA was extracted using the Monarch Plasmid Miniprep kit (NEB, T10104).

Four wells of a 6-well plate were seeded with 6.25×10^^5^ HEK293 cells and grown in DMEM (Gibco) supplemented with 10% FBS and 1% Pen-Strep at 37°C in a humidified incubator with 5% CO_2_. 24 hours after seeding, cells were transiently transfected with 2.5ug of the *NOTCH2tv*, *NOTCH2NLR*, *NOTCH2NLB*, or pEF-GFP (for GFP Ab control) expression plasmid construct using a 3:1 μl/μg ratio of Lipofectamine LTX Reagent (ThermoFisher) according to the manufacturer’s protocol. Cells were harvested 48 hours after transfection, washed in 1mL of cold PBS, resuspended in 250 ul of cold RIPA buffer (5M NaCl, 1M Tris-HCl pH 8.0, 1% NP40, 10% sodium deoxycholate, 10% SDS, 1mM PMSF, 1x Protease Inhibitor tablet (Pierce)) and incubated in a thermomixer at 4°C and 500 rpm for 20 minutes. The lysis was then spun down at 16,000 rpm for 20 minutes and the supernatant collected. 12μL of the cleared lysate was supplemented with 4μL of 4x LDS Sample Buffer (Invitrogen) and boiled at 70°C for 10 minutes. A 4-12% Bis-Tris gel (Invitrogen) was loaded with 15μL of each sample in duplicate and run in MOPS buffer at 200V for 50 minutes. The gels were then transferred onto a 0.45 μm nitrocellulose membrane (Bio-Rad) using a genie transfer apparatus (Idea Scientific) at 12V for 90 minutes. The membrane was incubated in blocking buffer (5% milk in TBST) at room temperature for 1 hour before cutting the membrane in half and incubating with a 1:1000 dilution of either primary anti-HA (Cell Signaling Technology, 3724T) or primary anti-GFP (Cell Signaling Technology, 29565) in blocking buffer and incubated at 4°C overnight. The following day membranes were washed 3x with 10mL of blocking buffer followed by a 1-hour incubation in a 1:20,000 dilution of IRDye 800CW secondary Ab (LI-COR, 926-32213) in blocking buffer. The membrane was washed 3x in TBST and imaged on an Odyssey imaging system (LI-COR) (Figure S12).

### Predicting TADs associated with *NOTCH2NL* paralogs using FiberFold

To generate predicted contact maps surrounding *NOTCH2/NL* promoters, we used the recently released tool FiberFold^34^. We first collapsed single-molecule Fiber-seq data into one-dimensional bigWig tracks. These single-base-pair resolution bigWigs, which represent FIRE density, CpG methylation state, and CTCF footprinting state, were generated following the protocol described at https://github.com/altemoselab/fiberFold. Phased Fiber-seq data were used to produce haplotype-specific bigWigs, and all maps were oriented to maintain consistent directionality relative to the *NOTCH2* promoter. To compute mean squared error (MSE) plots, we calculated the element-wise difference between the two contact maps and squared the resulting values to generate the distribution of differential contacts between the loci.

## SUPPLEMENTAL INFORMATION

Document S1. Tables S1–S5, Figures S1–S13, and supplemental references Table S6. Duplicon spreadsheet, related to Figure 1

---

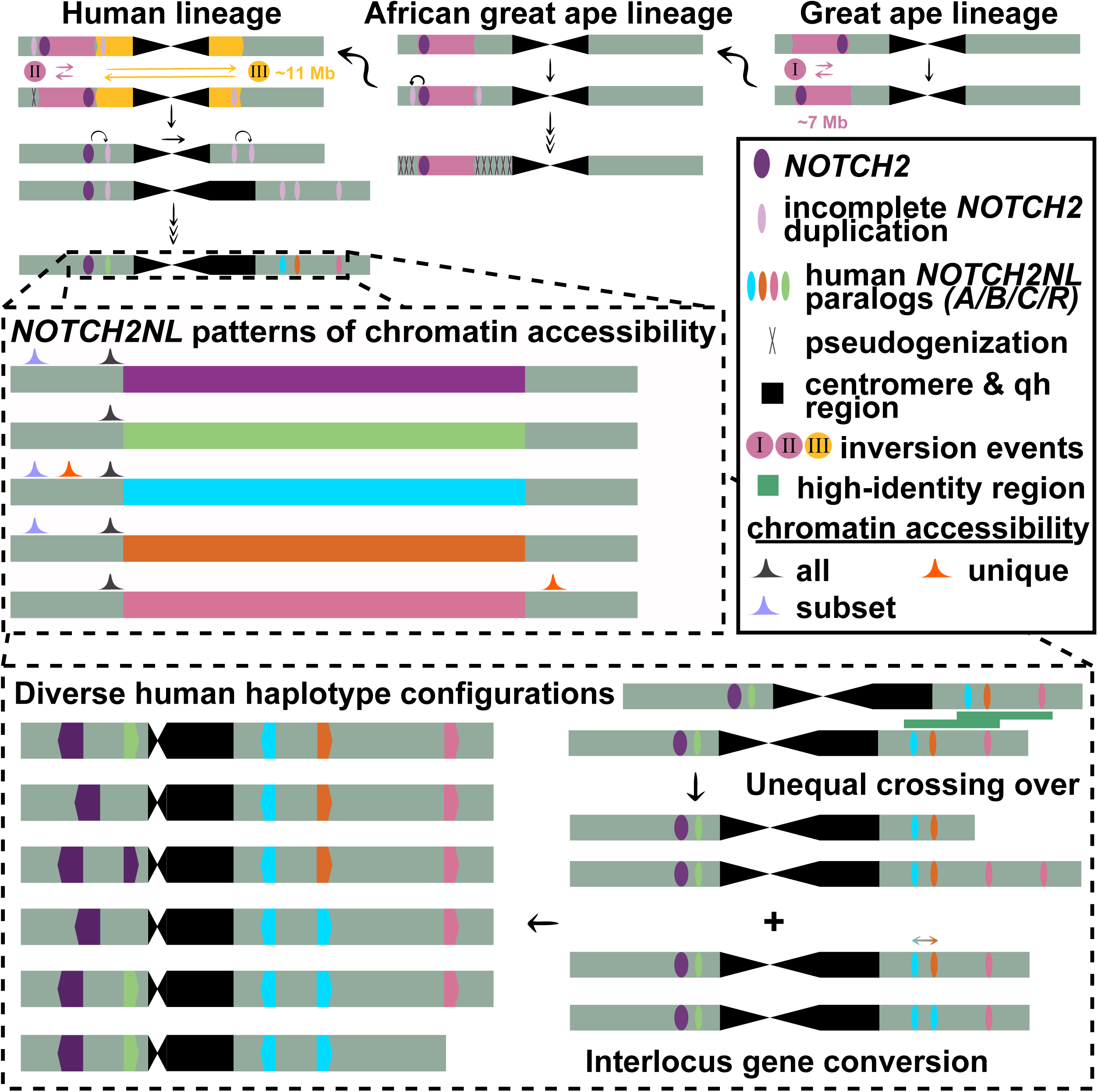

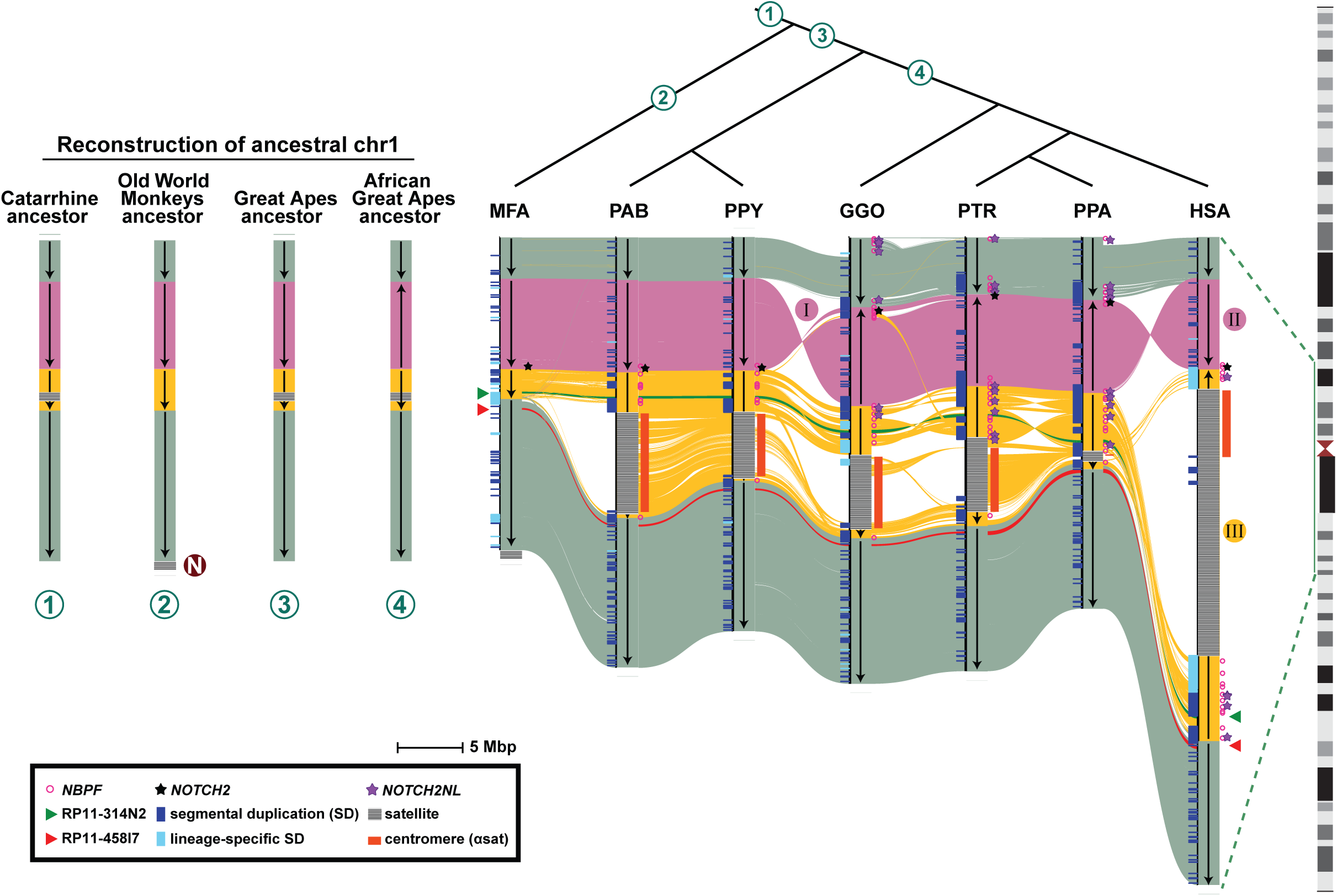

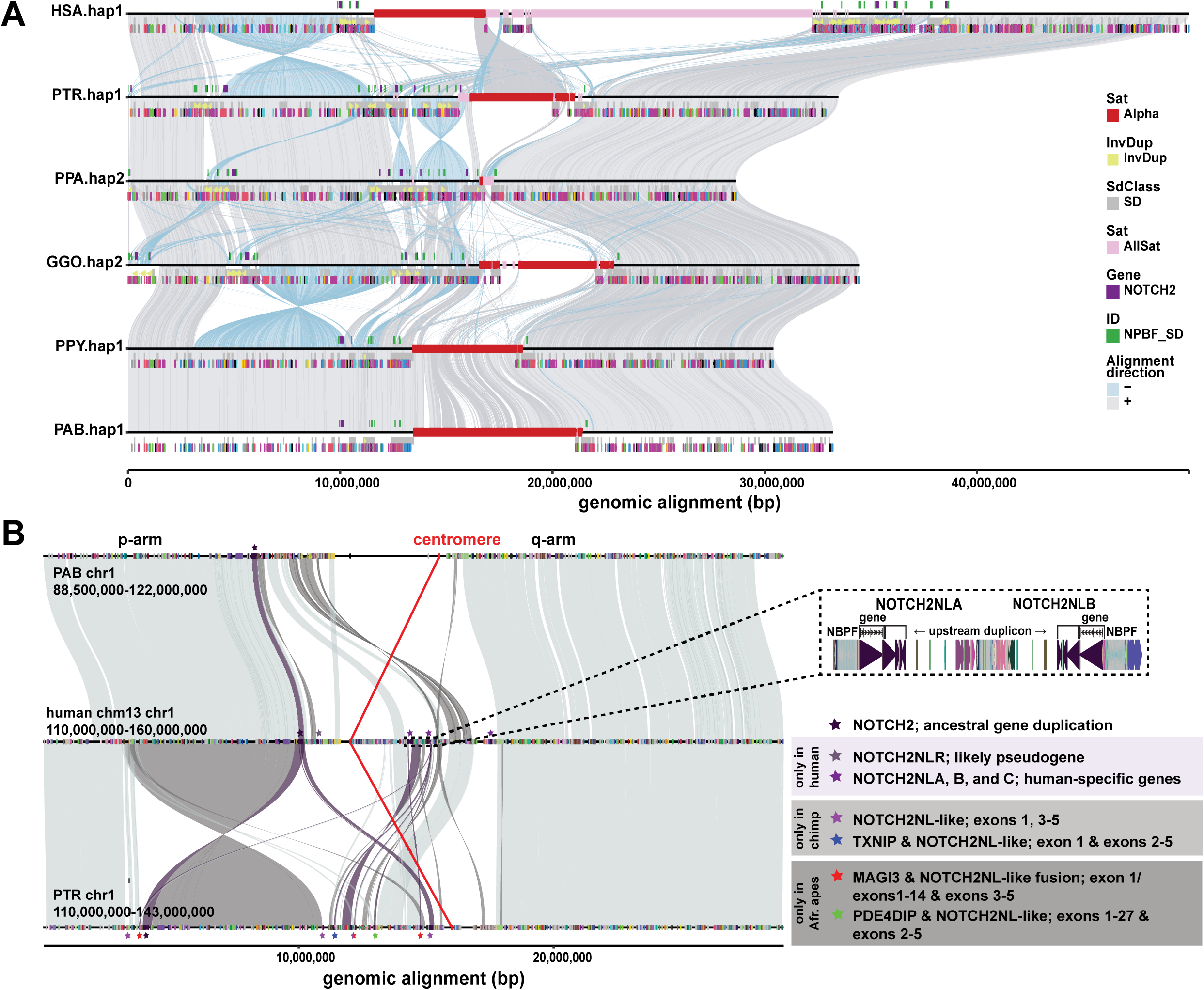

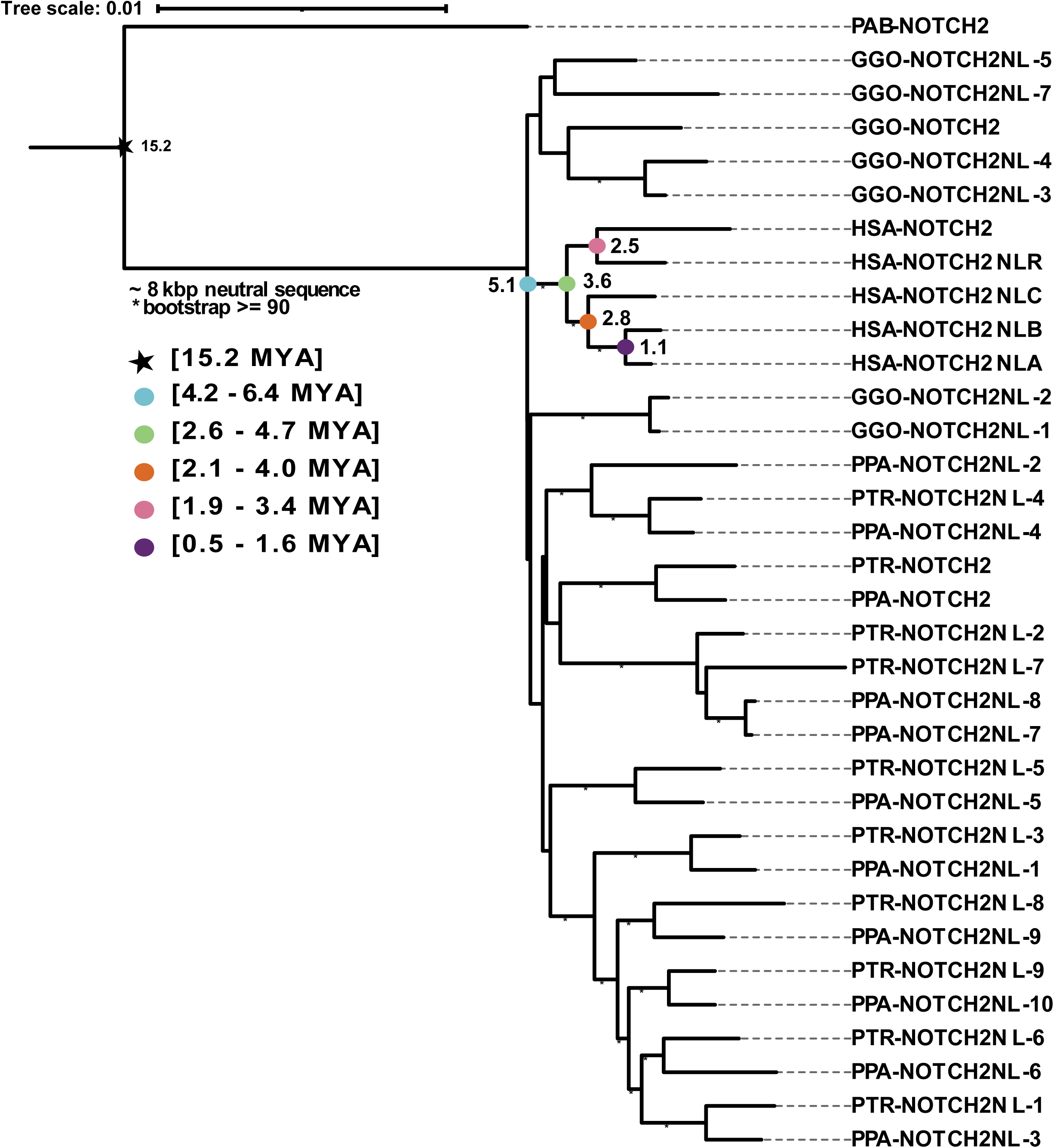

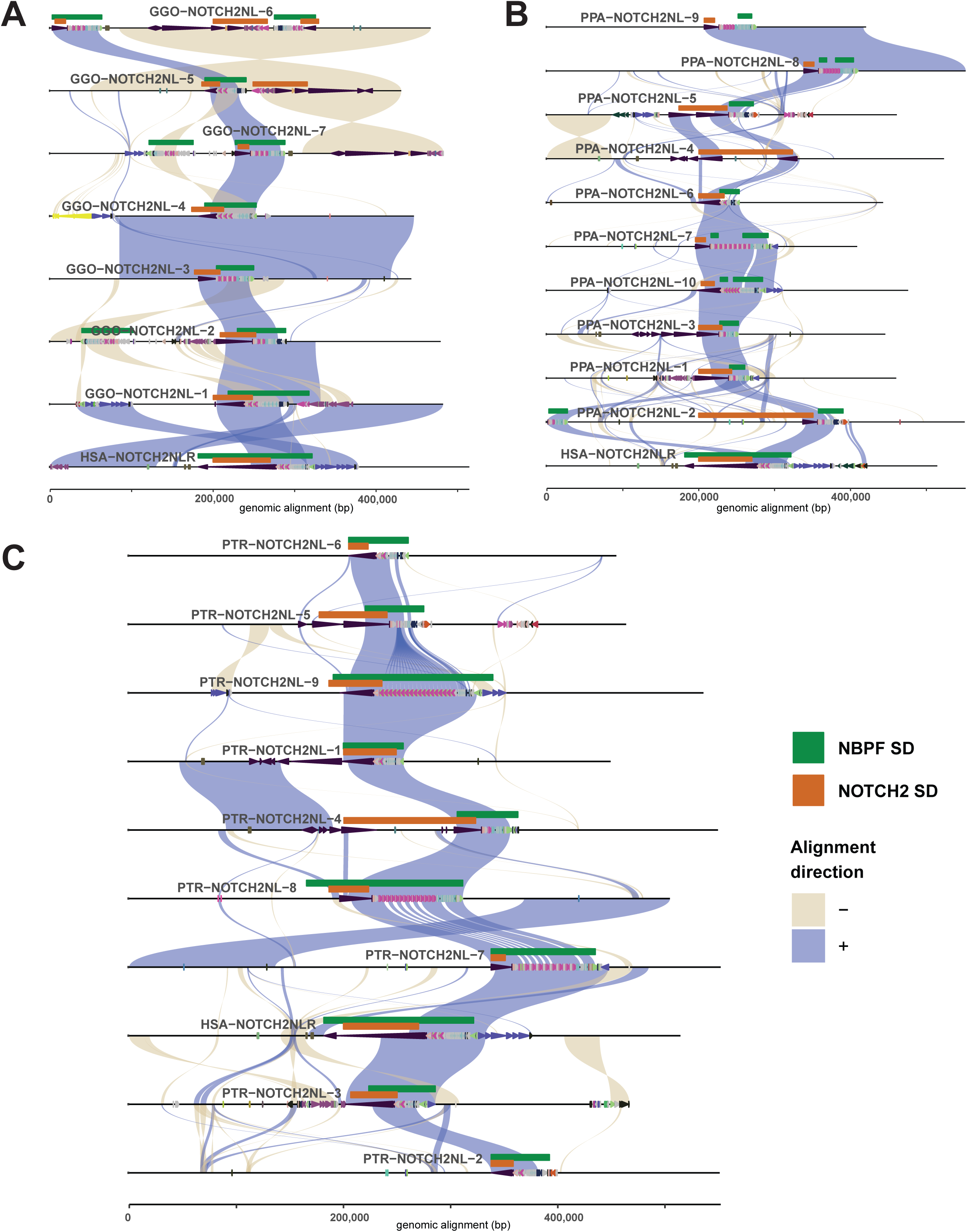

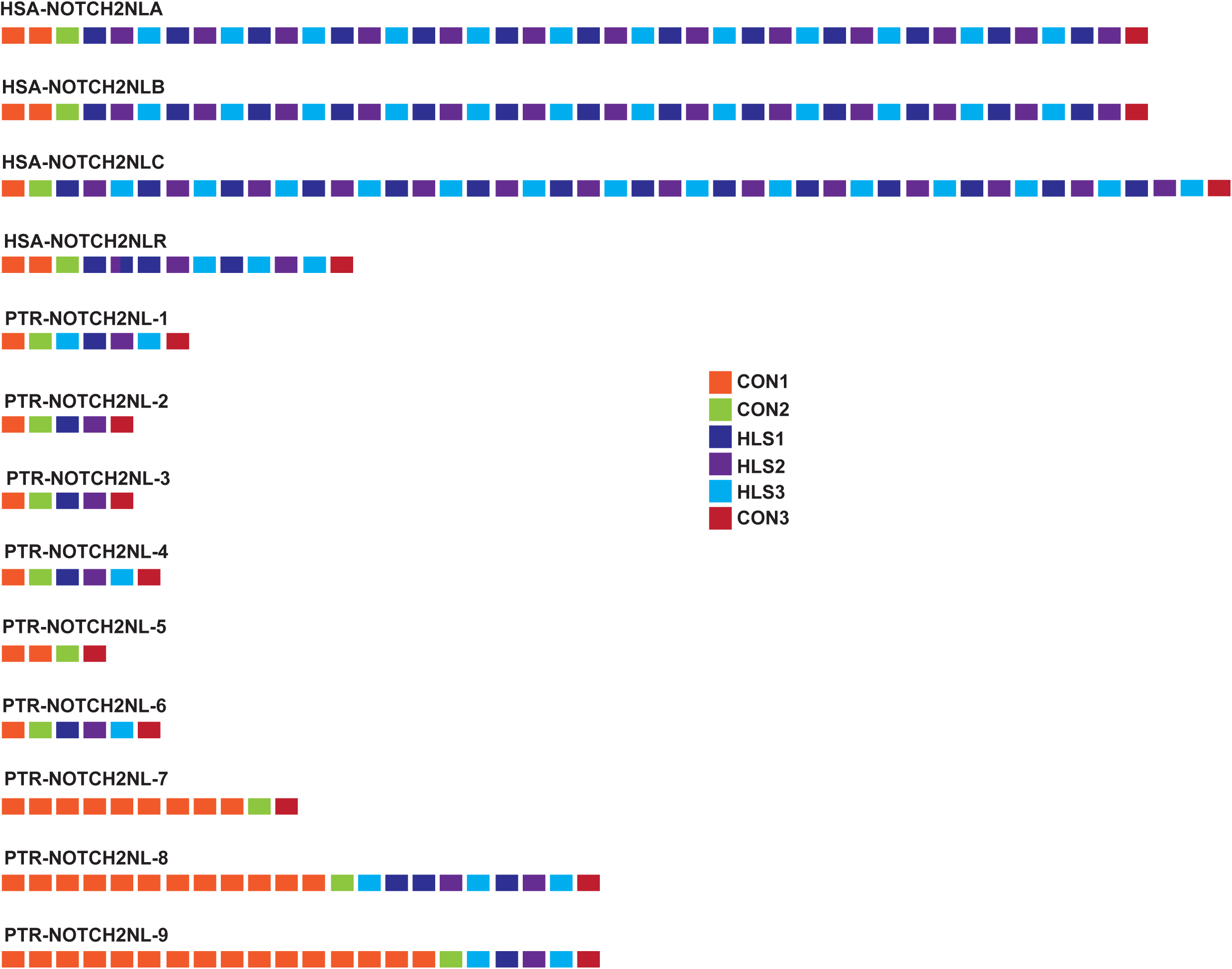

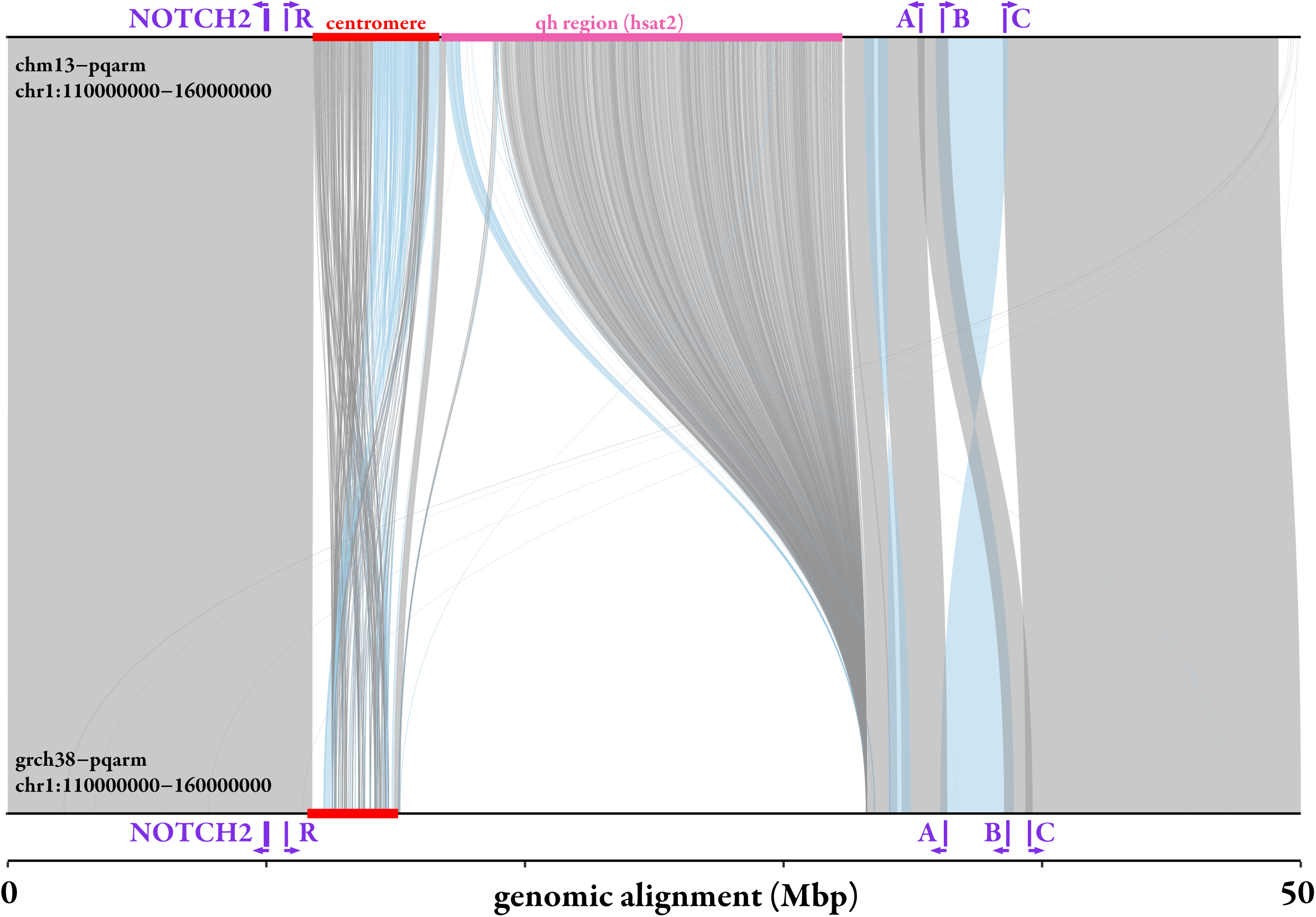

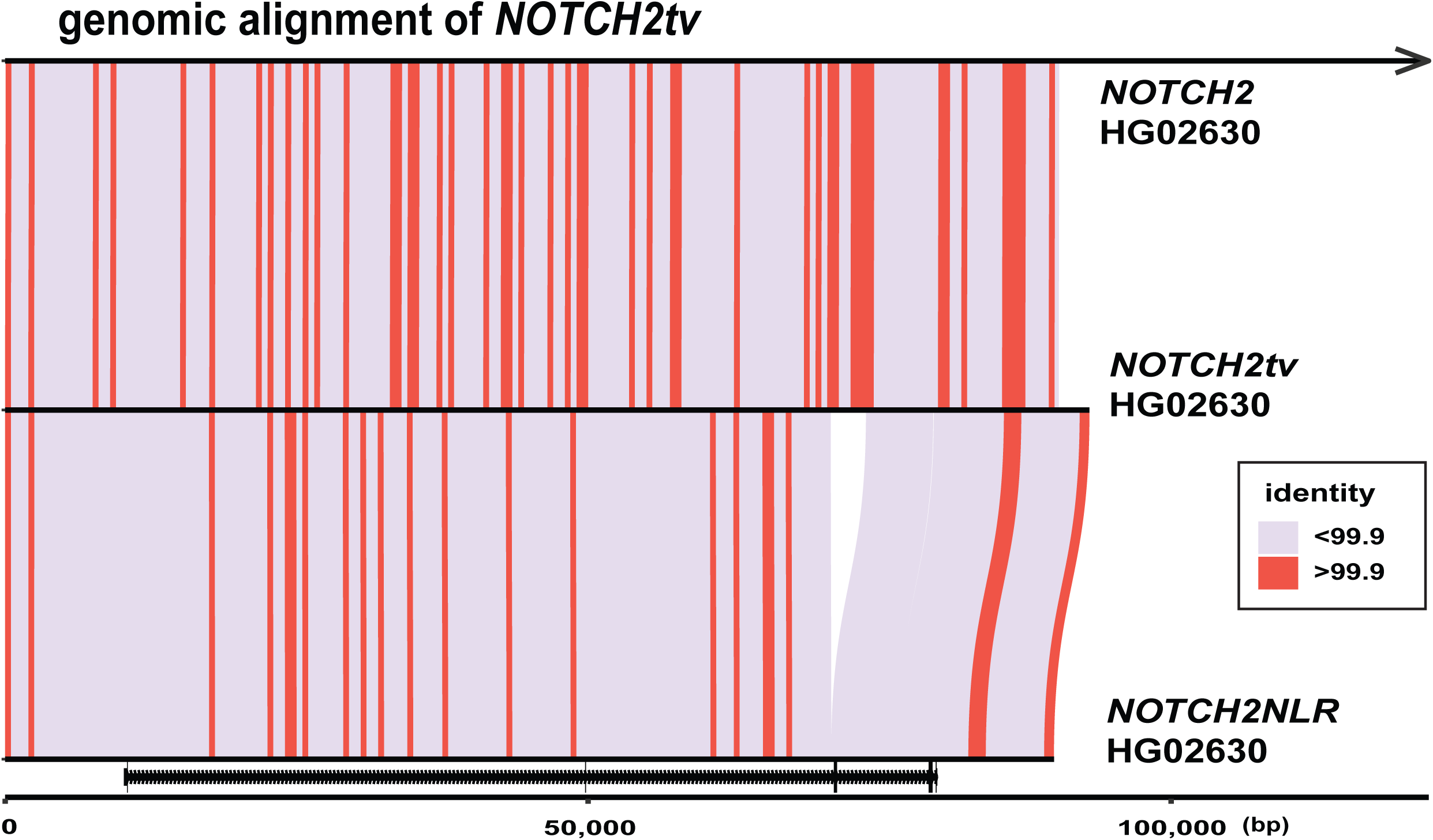

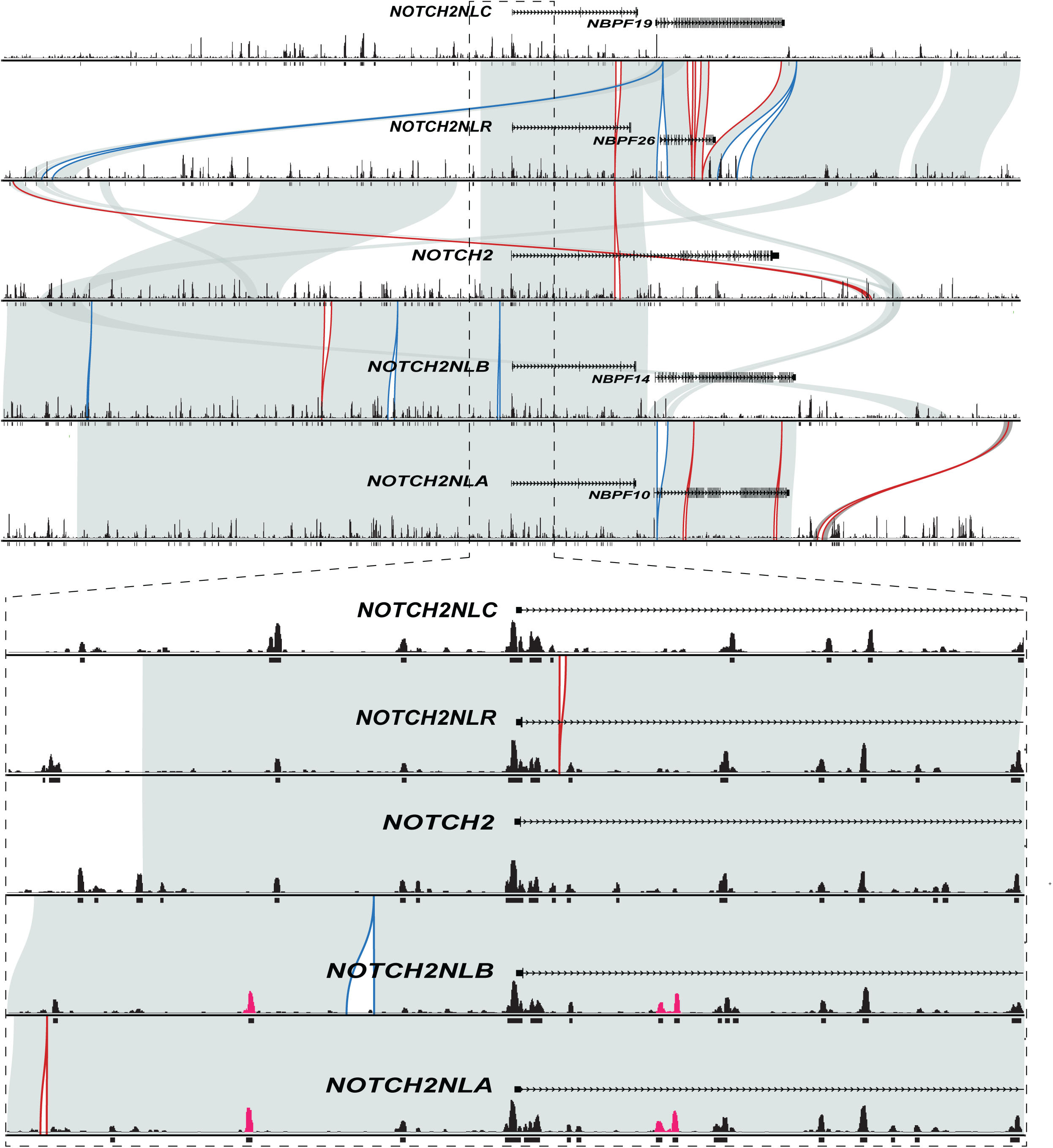

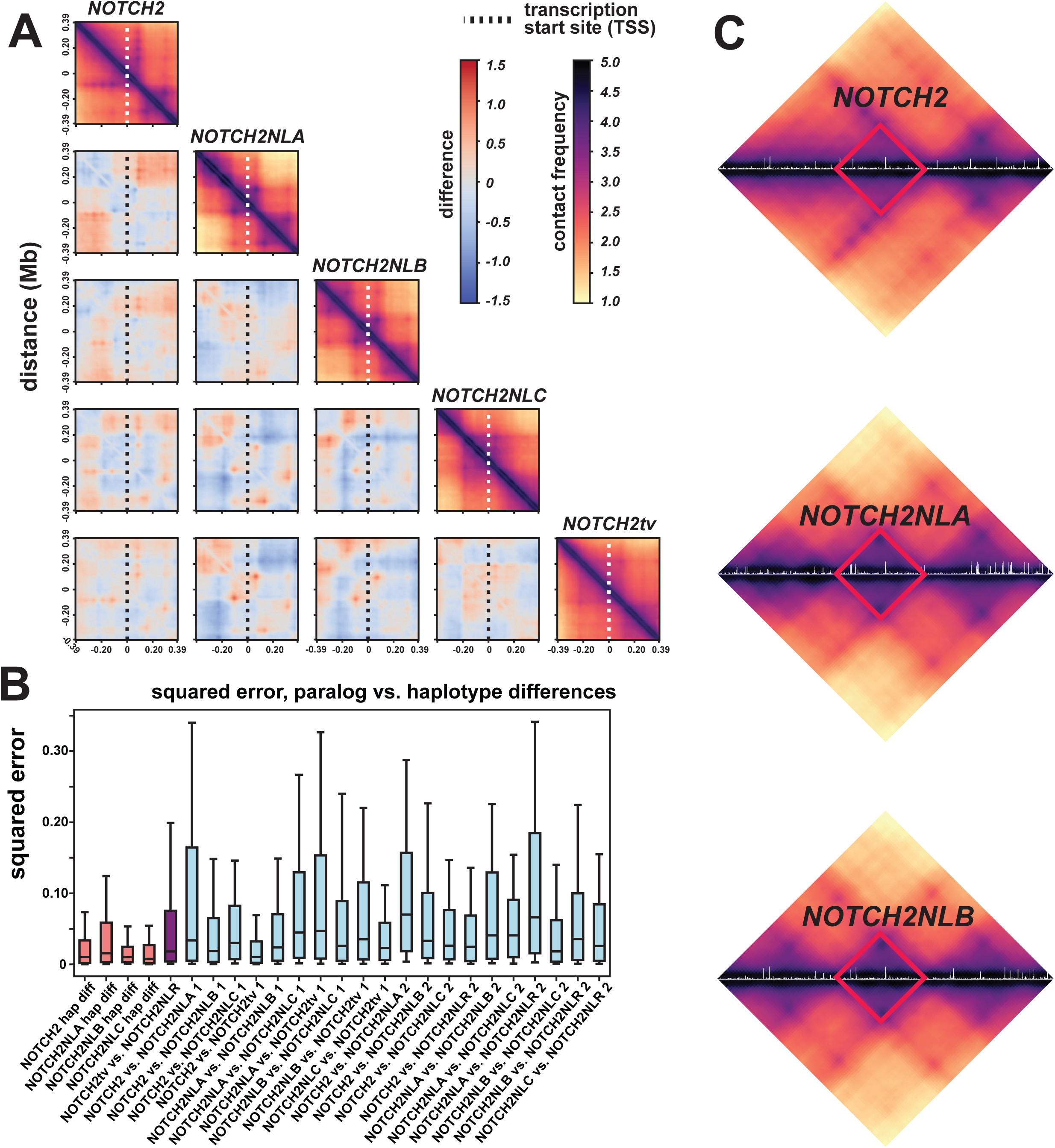

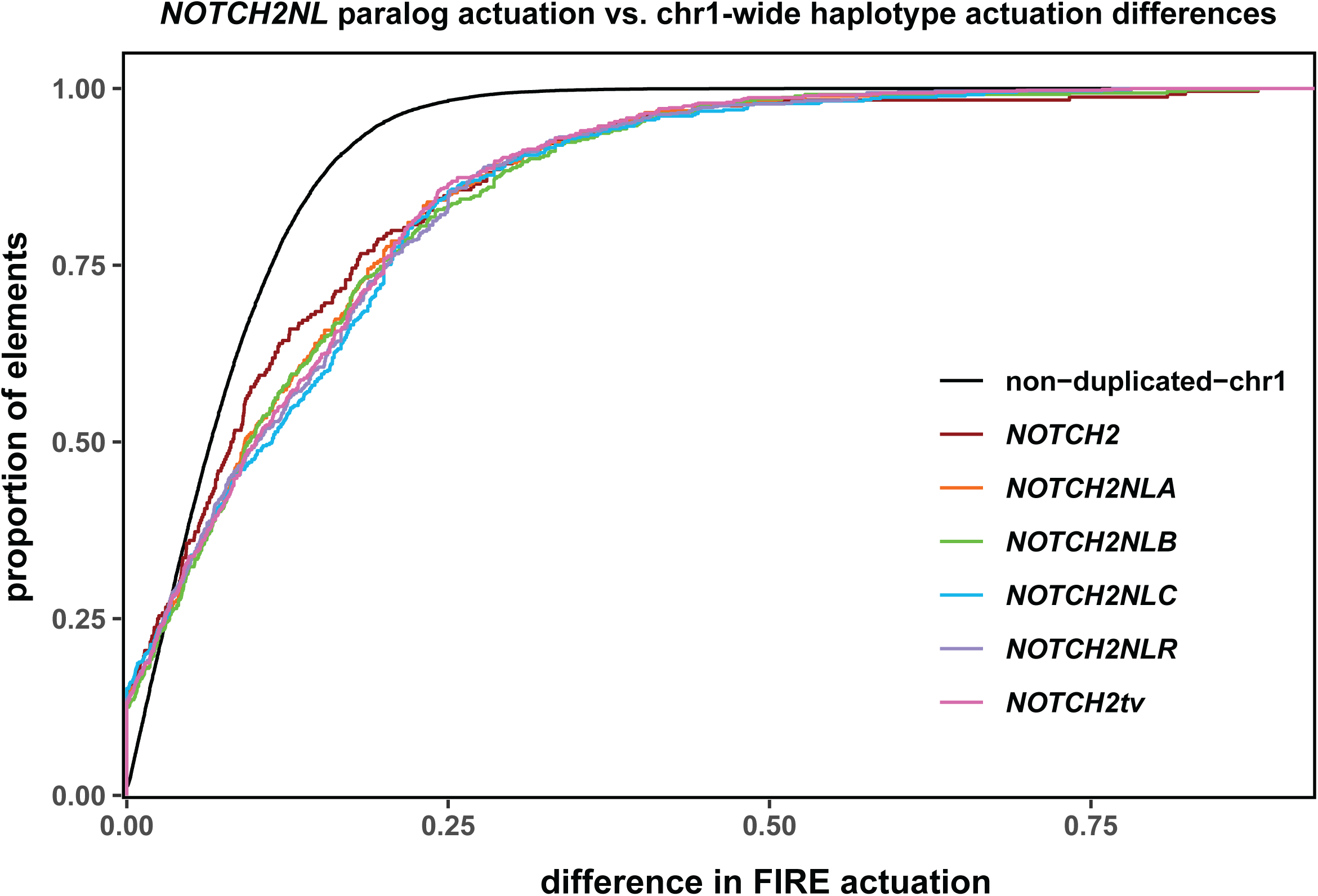

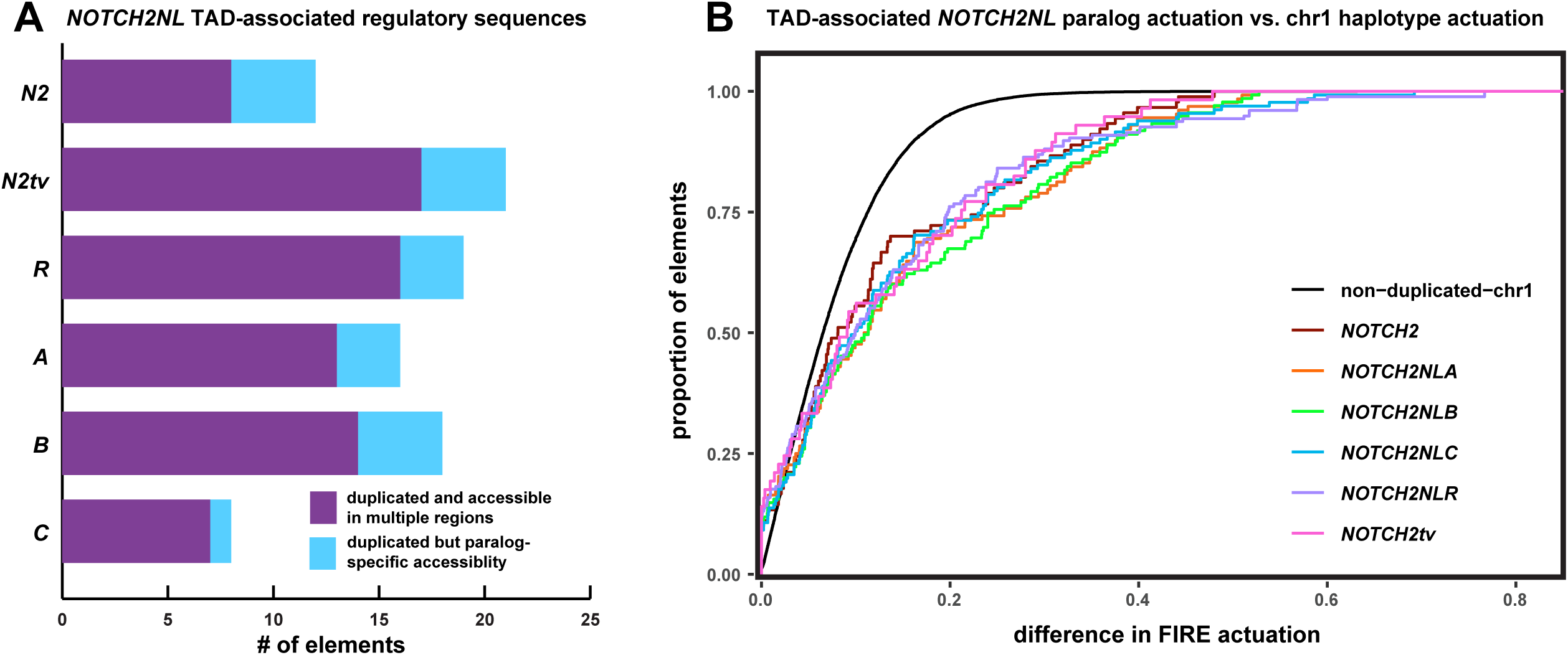

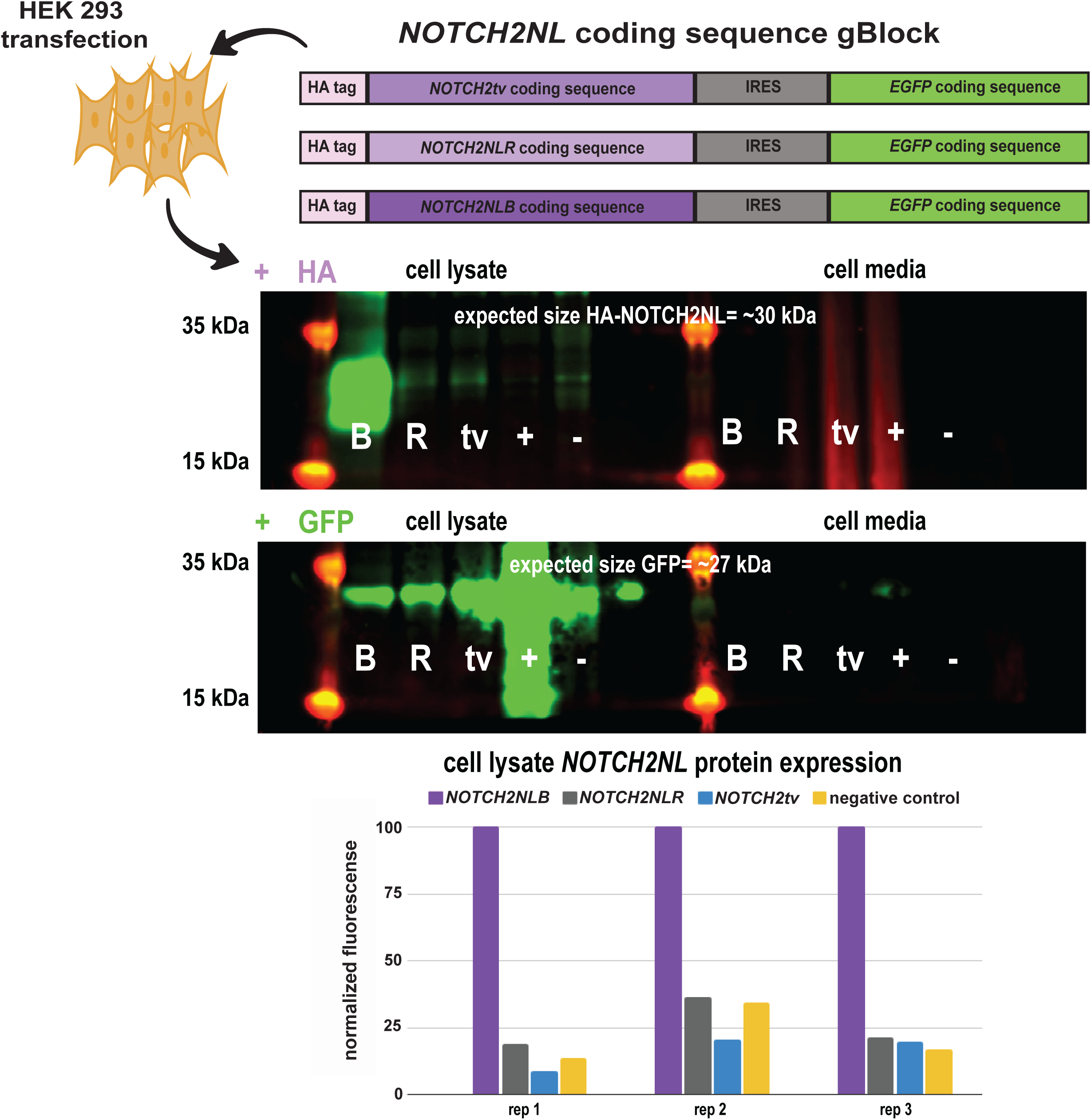

## Notes

https://www.ncbi.nlm.nih.gov/bioproject/1236375

https://zenodo.org/records/15022214

https://github.com/tdreal/NOTCH2NL-0325/tree/main

### Summary of Updates

This version of the manuscript has been edited to include an additional author, FiberFold analysis, as well as streamlined per comments during the revision process.

