## Supplementary Materials for "Genetic diversity and regulatory features of human-specific *NOTCH2NL* duplications"

### SUPPLEMENTAL MATERIAL

### ADDITIONAL TABLES & FIGURES

**Table S1. Structural variants of *NOTCH2NL* paralogs in T2T-CHM13, related to Figure 1C.**

| query paralog | ref paralog | query start | query end | mapq | structural variant size | in. (I) / del (D) |
|---|---|---|---|---|---|---|
| NOTCH2 | NOTCH2NLC | 119846347 | 119900735 | 60 | 54389 | I |
| NOTCH2 | NOTCH2NLC | 119582683 | 119582682 | 60 | 737 | D |
| NOTCH2 | NOTCH2NLC | 119541422 | 119547699 | 60 | 6278 | I |
| NOTCH2 | NOTCH2NLC | 119577460 | 119579275 | 60 | 1816 | I |
| NOTCH2 | NOTCH2NLB | 119542857 | 119555494 | 60 | 12638 | I |
| NOTCH2 | NOTCH2NLB | 119814488 | 119818620 | 60 | 4133 | I |
| NOTCH2 | NOTCH2NLB | 119461254 | 119461253 | 60 | 5629 | D |
| NOTCH2 | NOTCH2NLB | 119461388 | 119461387 | 60 | 4741 | D |
| NOTCH2 | NOTCH2NLB | 119462558 | 119462557 | 60 | 996 | D |
| NOTCH2 | NOTCH2NLB | 119677199 | 119677198 | 60 | 748 | D |
| NOTCH2 | NOTCH2NLB | 119857611 | 119857610 | 60 | 3875 | D |
| NOTCH2 | NOTCH2NLB | 119917835 | 119917834 | 60 | 1358 | D |
| NOTCH2 | NOTCH2NLB | 120135864 | 120137057 | 60 | 1194 | I |
| NOTCH2 | NOTCH2NLB | 120134461 | 120135831 | 60 | 1371 | I |
| NOTCH2 | NOTCH2NLA | 119814488 | 119818620 | 60 | 4133 | I |
| NOTCH2 | NOTCH2NLA | 119677199 | 119677198 | 60 | 748 | D |
| NOTCH2 | NOTCH2NLA | 119857611 | 119857610 | 60 | 3852 | D |
| NOTCH2 | NOTCH2NLA | 119917825 | 119917824 | 60 | 1360 | D |
| NOTCH2 | NOTCH2NLA | 119542836 | 119555473 | 60 | 12638 | I |
| NOTCH2 | NOTCH2NLA | 119462604 | 119462603 | 60 | 996 | D |
| NOTCH2 | NOTCH2NLA | 119461389 | 119461388 | 60 | 4741 | D |
| NOTCH2 | NOTCH2NLA | 119461253 | 119461252 | 60 | 5629 | D |
| NOTCH2 | NOTCH2NLA | 120134365 | 120134947 | 31 | 583 | I |
| NOTCH2 | NOTCH2NLA | 120135063 | 120135832 | 31 | 770 | I |
| NOTCH2 | NOTCH2NLA | 120135863 | 120137056 | 31 | 1194 | I |
| NOTCH2NLA | NOTCH2NLC | 145263860 | 145265217 | 60 | 1358 | I |
| NOTCH2NLA | NOTCH2NLC | 145426571 | 145428098 | 60 | 1528 | I |
| NOTCH2NLA | NOTCH2NLC | 145431274 | 145434427 | 60 | 3154 | I |
| NOTCH2NLA | NOTCH2NLC | 144857035 | 144858873 | 60 | 1839 | I |
| NOTCH2NLA | NOTCH2NLB | 145373282 | 145374823 | 60 | 1542 | I |
| NOTCH2NLA | NOTCH2NLB | 145426775 | 145428316 | 60 | 1542 | I |
| NOTCH2NLA | NOTCH2NLB | 145357941 | 145357940 | 60 | 6296 | D |
| NOTCH2NLA | NOTCH2NLB | 145452444 | 145455266 | 60 | 2823 | I |
| NOTCH2NLB | NOTCH2NLC | 146024204 | 146083108 | 60 | 58905 | I |
| NOTCH2NLB | NOTCH2NLC | 146100183 | 146101538 | 60 | 1356 | I |
| NOTCH2NLB | NOTCH2NLC | 146196656 | 146202946 | 60 | 6291 | I |
| NOTCH2NLB | NOTCH2NLC | 146270637 | 146273780 | 60 | 3144 | I |
| NOTCH2NLB | NOTCH2NLC | 146282918 | 146295060 | 60 | 12143 | I |
| NOTCH2NLB | NOTCH2NLC | 146295069 | 146303567 | 60 | 8499 | I |
| NOTCH2NLB | NOTCH2NLC | 146215843 | 146215842 | 60 | 1549 | D |
| NOTCH2NLB | NOTCH2NLC | 146461653 | 146461652 | 60 | 2066 | D |
| NOTCH2NLB | NOTCH2NLC | 145759226 | 145761064 | 60 | 1839 | I |
| NOTCH2NLC | NOTCH2NLR | 148580984 | 148584007 | 60 | 3024 | I |
| NOTCH2NLC | NOTCH2NLR | 148623116 | 148626328 | 60 | 3213 | I |
| NOTCH2NLC | NOTCH2NLR | 148627877 | 148631089 | 60 | 3213 | I |
| NOTCH2NLC | NOTCH2NLR | 148635700 | 148678456 | 60 | 42757 | I |
| NOTCH2NLC | NOTCH2NLR | 148608665 | 148608664 | 60 | 6274 | D |
| NOTCH2NLC | NOTCH2NLR | 148687503 | 148687502 | 60 | 11467 | D |
| NOTCH2NLC | NOTCH2NLR | 148687511 | 148687510 | 60 | 8199 | D |
| NOTCH2NLC | NOTCH2NLR | 148867048 | 148870884 | 60 | 3837 | I |
| NOTCH2NLC | NOTCH2NLR | 148443146 | 148443145 | 60 | 11467 | D |
| NOTCH2NLC | NOTCH2NLR | 148443154 | 148443153 | 60 | 8199 | D |
| NOTCH2NLC | NOTCH2NLR | 148335672 | 148335671 | 60 | 1816 | D |
| NOTCH2NLR | NOTCH2NLB | 120452239 | 120464876 | 60 | 12638 | I |

| | | | | | | |
|---|---|---|---|---|---|---|
| NOTCH2NLR | NOTCH2NLB | 120546480 | 120546479 | 60 | 5629 | D |
| NOTCH2NLR | NOTCH2NLB | 120546346 | 120546345 | 60 | 4741 | D |
| NOTCH2NLR | NOTCH2NLB | 120545176 | 120545175 | 60 | 996 | D |
| NOTCH2NLR | NOTCH2NLB | 120330535 | 120330534 | 60 | 748 | D |
| NOTCH2NLR | NOTCH2NLB | 120730144 | 120730143 | 60 | 1356 | D |
| NOTCH2NLR | NOTCH2NLB | 120798010 | 120798009 | 60 | 3025 | D |
| NOTCH2NLR | NOTCH2NLB | 120841724 | 120841723 | 60 | 3184 | D |
| NOTCH2NLR | NOTCH2NLB | 120843404 | 120843403 | 60 | 3172 | D |
| NOTCH2NLR | NOTCH2NLB | 120845053 | 120845052 | 60 | 3202 | D |
| NOTCH2NLR | NOTCH2NLB | 120849542 | 120849541 | 60 | 40814 | D |
| NOTCH2NLR | NOTCH2NLB | 120861687 | 120861686 | 60 | 654 | D |
| NOTCH2NLR | NOTCH2NLB | 120594342 | 120594341 | 60 | 748 | D |
| NOTCH2NLR | NOTCH2NLA | 120452260 | 120464897 | 60 | 12638 | I |
| NOTCH2NLR | NOTCH2NLA | 120545130 | 120545129 | 60 | 996 | D |
| NOTCH2NLR | NOTCH2NLA | 120546345 | 120546344 | 60 | 4741 | D |
| NOTCH2NLR | NOTCH2NLA | 120546481 | 120546480 | 60 | 5629 | D |
| NOTCH2NLR | NOTCH2NLA | 120594342 | 120594341 | 60 | 748 | D |
| NOTCH2NLR | NOTCH2NLA | 120820315 | 120826597 | 60 | 6283 | I |
| NOTCH2NLR | NOTCH2NLA | 120730144 | 120730143 | 60 | 1358 | D |
| NOTCH2NLR | NOTCH2NLA | 120798010 | 120798009 | 60 | 3025 | D |
| NOTCH2NLR | NOTCH2NLA | 120843404 | 120843403 | 60 | 3180 | D |
| NOTCH2NLR | NOTCH2NLA | 120844944 | 120844943 | 60 | 3168 | D |
| NOTCH2NLR | NOTCH2NLA | 120330535 | 120330534 | 60 | 748 | D |
| NOTCH2NLR | NOTCH2 | 120798010 | 120798009 | 60 | 3028 | D |
| NOTCH2NLR | NOTCH2 | 120838292 | 120846100 | 60 | 7809 | I |

**Table S2. Additional structural variation between human and NHA, related to Figure 2.**

| Variant information | | | | NOTCH2 mapping (primary hap) | | INV BP mapping (T2T-space; kbp) | | |
|---|---|---|---|---|---|---|---|---|
| Species | Variants | Count | Bases (kbp) | Count | Bases (kbp) | Count | Bases (kbp) | p-value |
| PPY | PAV-DEL | 816 | 752.8 | 3 | 0.7 | 71 | 99.5 | 1.000 |
| PPY | PAV-INS | 829 | 1025.1 | 6 | 9.1 | 82 | 140.0 | 1.000 |
| PPY | Syri/PAV-INV | 6 | 930.4 | - | - | - | - | - |
| PPY | Sedef-SD (h1/h2) | 98/97 | 2588.9/2588.9 | 1 | 10.6 | - | - | - |
| PAB | PAV-DEL | 810 | 751.7 | 3 | 0.7 | 72 | 106.1 | 1.000 |
| PAB | PAV-INS | 812 | 1016.1 | 5 | 9.0 | 74 | 123.7 | 1.000 |
| PAB | Syri/PAV-INV | 4 | 1338.3 | - | - | - | - | - |
| PAB | Sedef-SD (h1/h2) | 97/95 | 2611.4/2576.7 | 1 | 10.3 | - | - | - |
| PTR | PAV-DEL | 343 | 341.3 | 3 | 5.2 | 29 | 69.3 | 0.987 |
| PTR | PAV-INS | 312 | 1146.2 | 2 | 604.9 | 40 | 883.4 | **<0.001*** |
| PTR | Syri/PAV-INV | 3 | 8017.1 | - | - | - | - | - |
| PTR | Sedef-SD (h1/h2) | 109/111 | 6565.9/6582.8 | 10 | 3660.8 | - | - | - |
| GGO | PAV-DEL | 475 | 591.1 | 5 | 11.1 | 36 | 141.7 | 0.832 |
| GGO | PAV-INS | 422 | 1010.9 | 0 | 0.0 | 35 | 223.8 | **0.034*** |
| GGO | Syri/PAV-INV | 6 | 8885.6 | - | - | - | - | - |
| GGO | Sedef-SD (pri/alt) | 114/118 | 7103.1/7049.3 | 6 | 3043.9 | - | - | - |
| PPA | PAV-DEL | 353 | 359.9 | 3 | 5.2 | 19 | 51.0 | 0.999 |
| PPA | PAV-INS | 308 | 537.7 | 1 | 3.0 | 25 | 262.5 | **<0.001*** |
| PPA | Syri/PAV-INV | 6 | 9691.5 | - | - | - | - | - |
| PPA | Sedef-SD (pri/alt) | 110/111 | 6459.5/6059.9 | 10 | 3532.2 | - | - | - |
| HSA | Sedef-SD | 91 | 8315.6 | 3 | 3988.8 | 30 | 1918.8 | **<0.001*** |
| | | | **Average** | 3.9 | 931.0 | 46.6 | 365.4 | |

### Table S3. NHA *NOTCH2NL* FLNC characterization, related to Figure 3.

| Name | Iso-Seq transcript | Chr | Start | End | Strand | # of exons | Predicted AA Lengths | Long-read transcript support (NEC & testis) | Intergenic distance N2NL-NBPF (bp) | Species | Comments |
|---|---|---|---|---|---|---|---|---|---|---|---|
| PTR-NOTCH2 | NOTCH2 | chr1 | 117131242 | 117288384 | - | -- | 2432 | -- | -- | PTR/PPA/GGO/HSA | -- |
| PTR-NOTCH2NL-1 | NOTCH2NLR | chr1 | 106135723 | 106156373 | + | 4 | 246 | 26 | 11,622 | PPA/PTR | ORF has exons 1, 3-5 of NOTCH2NLR |
| | Fusion NOTCH2NL-NBPF | chr1 | 106135792 | 106185074 | | 20 | 851 | 69 | | | ORF has exons 1, 3, 4 of NOTCH2NLR & NBPF exons |
| PTR-NOTCH2NL-2 | Fusion MAGI3-NOTCH2NL | chr1 | 106365562 | 106516651 | + | 4 | 324 | 6 | 11,398 | PPA/PTR | ORF has MAGI3 exon 1 & exons 3-5 of NOTCH2NLR |
| PTR-NOTCH2NL-3 | NOTCH2NLR | chr1 | 108282469 | 108322382 | + | 4 | 235 | 16 | 11,838 | PPA/PTR/GGO | ORF has exons 2-5 of NOTCH2NLR |
| | Fusion NOTCH2NL-NBPF | chr1 | 108282319 | 108349249 | | 13 | 672 | 11 | | | ORF has exons 2, 3, 4 of NOTCH2NLR & NBPF exons |
| | Fusion PDE4DIP-NOTCH2NL-NBPF | chr1 | 108214852 | 108349197 | | -- | -- | 53 | | | No ORF goes through the whole transcript |
| PTR-NOTCH2NL-4 | Fusion MAGI3-NOTCH2NL | chr1 | 109104609 | 109227579 | - | 13 | 845 | 12 | 11,410 | PPA/PTR | ORF has exon 1 of NOTCH2NLR, 9 MAGI3, and 3-5 exons of NOTCH2NLR |
| | Fusion MAGI3-NOTCH2NL-NBPF | chr1 | 109078294 | 109227828 | | 17 | 1,411 | 53 | | | ORF has exon 1 of NOTCH2NLR, 9 MAGI3, NOTCH2NL exons 3, 4, and NBPF exons |
| PTR-NOTCH2NL-5 | NOTCH2NLR | chr1 | 109873983 | 109910441 | - | 3 | 146 | 9 | 11,295 | PPA/PTR | ORF has exons 2, 3 of NOTCH2NLR & partial exon 4 (TRUNCATION) - transcript goes through until NOTCH2NLR-like exon 5 |
| | Fusion NOTCH2NL-NBPF | chr1 | 109846910 | 109910766 | | 3 | 146 | 12 | | | ORF has exons 2, 3 of NOTCH2NLR & partial exon 4 (TRUNCATION) - transcript goes through to NBPF exons |
| | Fusion TXNIP-NOTCH2NL-NBPF | chr1 | 109846504 | 109953395 | | 4 | 230 | 62 | | | ORF has exon 1 of TXNIP, exons 2,3 of NOTCH2NLR, and partial exon 4 (TRUNCATION) - transcript goes through until NOTCH2NLR-like exon 5 |
| PTR-NOTCH2NL-6 | NOTCH2NLR | chr1 | 110315936 | 110336372 | + | 4 | 243 | 11 | 11,504 | PPA/PTR/GGO | ORF has exon 1, 3-5 of NOTCH2NLR |
| | Fusion NOTCH2NL-NBPF | chr1 | 110313408 | 110368080 | | 20 | 856 | 22 | | | ORF that has exons 1, 3, 4 of NOTCH2NLR & NBPF exons |
| | Fusion SORT1-LRIG2-NOTCH2NL-NBPF | chr1 | 110225344 | 110368328 | | -- | -- | 14 | | | No ORF goes through the whole transcript |
| PTR-NOTCH2NL-7 | Fusion with MAGI3 | chr1 | 117544406 | 117696496 | - | 4 | 324 | 7 | 11,693 | PPA/PTR | ORF has exon 1 of MAGI3 & exons 3-5 of NOTCH2NLR |
| PTR-NOTCH2NL-8 | NOTCH2NLR | chr1 | 118044677 | 118065257 | - | 4 | 246 | 9 | 11,295 | PTR | ORF has exons 1, 3-5 of NOTCH2NLR |
| | Fusion NOTCH2NL-NBPF | chr1 | 117960416 | 118065110 | | 58 | 3033 | 16 | | | ORF has exons 1, 3, 4 of NOTCH2NLR & NBPF exons |
| | Fusion LRIG2-NOTCH2NL | chr1 | 118044029 | 118124464 | | -- | -- | 5 | | | No ORF goes through the whole transcript |
| | Fusion LRIG2-NOTCH2NL-NBPF | chr1 | 117960416 | 118125899 | | -- | -- | 4 | | | No ORF goes through the whole transcript |
| PTR-NOTCH2NL-9 | NOTCH2NLR | chr1 | 121665898 | 121686392 | + | 4 | 245 | 11 | 11,486 | PPA/PTR | ORF has exons 1, 3-5 of NOTCH2NLR |
| | Fusion NOTCH2NL-NBPF | chr1 | 121665622 | 121801595 | | 88 | 4694 | 48 | | | ORF has exons 1, 3, 4 of NOTCH2NLR & NBPF exons |
| PPA-NOTCH2 | NOTCH2 | chr1 | 112555608 | 112716322 | - | -- | 2610 | -- | -- | PTR/PPA/GGO/HSA | -- |
| PPA-NOTCH2NL-1 | NOTCH2NLR | chr1 | 101669793 | 101703165 | - | 4 | 235 | 46 | 11,607 | PPA/PTR/GGO | ORF has exons 2-5 of NOTCH2NLR human |
| | Fusion NOTCH2NL-NBPF | chr1 | 101642931 | 101703233 | | 17 | 656 | 132 | | | ORF has exons 2, 3, 4 of NOTCH2NLR & NBPF exons |
| | Fusion PDE4DIP-NOTCH2NL-NBPF | chr1 | 101641810 | 101777882 | | -- | -- | 155 | | | No ORF goes through the whole transcript. |
| PPA-NOTCH2NL-2 | Fusion MAGI3-NOTCH2NL | chr1 | 103485748 | 103636324 | - | 4 | 237 | 6 | 11,837 | PPA/PTR | ORF has exon 1 of MAGI3 & exons 3-5 of NOTCH2NLR |
| PPA-NOTCH2NL-3 | NOTCH2NLR | chr1 | 103841187 | 103861631 | + | 4 | 246 | 11 | 11,672 | PPA/PTR | ORF has exons 1, 3-5 of NOTCH2NLR |
| | Fusion NOTCH2NL-NBPF | chr1 | 103815726 | 103861620 | | 15 | 757 | 98 | | | ORF has exons 1, 3, 4 of NOTCH2NLR & NBPF exons |
| PPA-NOTCH2NL-4 | Fusion MAGI3-NOTCH2NL | chr1 | 104697226 | 104820526 | | 13 | 845 | 201 | -- | PPA/PTR | ORF has exon 1 of NOTCH2NLR, 9 MAGI3, & 3-5 exons of NOTCH2NLR |
| PPA-NOTCH2NL-5 | NOTCH2NLR | chr1 | 105315877 | 105349528 | + | 4 | 235 | 11 | 11,690 | PPA/PTR | ORF has exons 2-5 of NOTCH2NLR |
| | Fusion NOTCH2NL-NBPF | chr1 | 105288382 | 105349479 | | 15 | 743 | 69 | | | ORF has exons 2, 3, 4 of NOTCH2NLR & NBPF exons |
| | Fusion TXNIP-NOTCH2NL-NBPF | chr1 | 105286533 | 105389706 | | 5 | 319 | 21 | | | ORF has exon 1 of TXNIP & exons 2, 3-5 of NOTCH2NLR |
| PPA-NOTCH2NL-6 | NOTCH2NLR | chr1 | 105752206 | 105772648 | - | 4 | 246 | 7 | 11,773 | PPA/PTR | ORF has exons 1, 3-5 of NOTCH2NLR |
| | Fusion NOTCH2NL-NBPF | chr1 | 105752087 | 105795054 | | 14 | 682 | 81 | | | ORF has exons 1, 3, 4 of NOTCH2NLR & NBPF exons |
| PPA-NOTCH2NL-7 | NOTCH2NLR | chr1 | 112968284 | 112977175 | - | 4 | 246 | 0 | 11,194 | PPA | ORF has exons 3-5 of NOTCH2NLR |
| PPA-NOTCH2NL-8 | Fusion MAGI3-NOTCH2NL | chr1 | 113432163 | 113583978 | - | 4 | 324 | 12 | 11,233 | PPA/PTR | ORF has exon 1 of MAGI3 & exons 3-5 of NOTCH2NLR |
| PPA-NOTCH2NL-9 | NOTCH2NLR | chr1 | 113892494 | 113913027 | - | 4 | 246 | 56 | 11,785 | PPA | ORF has exons 1, 3-5 of NOTCH2NLR |
| PPA-NOTCH2NL-10 | NOTCH2NLR | chr1 | 117521012 | 117542196 | + | 4 | 246 | 12 | 11,731 | PPA/PTR/GGO | ORF has exons 1, 3-5 of NOTCH2NLR |
| | Fusion NOTCH2NL-NBPF | chr1 | 117520465 | 117596479 | | 50 | 2399 | 35 | | | ORF has exons 1, 3, 4 of NOTCH2NLR & NBPF exons |

| | | | | | | | | | |
|---|---|---|---|---|---|---|---|---|---|
| | Fusion SORT1-LRIG2-NOTCH2NL-NBPF | chr1 117431222 117596326 | | -- | -- | 4 | | | No ORF goes through the whole transcript |
| GGO-NOTCH2 | NOTCH2 | chr1 124746752 124900741 | - | -- | 2611 | -- | -- | PTR/PPA/GGO/HSA | -- |
| GGO-NOTCH2NL-1 | NOTCH2NLR | chr1 116765261 116796179 | + | -- | -- | 2 | 11,618 | PPA/PTR/GGO | Has exon 3-5 of NOTCH2NLR but no exon 1/2 and no M start |
| | Fusion NOTCH2NL-NBPF | chr1 116765192 116847395 | | -- | -- | 13 | | | Has exon 3-4 of NOTCH2NLR & NBPF exons but no exon 1/2 & no M start |
| | Fusion SORT1-LRIG2-NOTCH2NL-NBPF | chr1 116693440 116846869 | | -- | -- | 6 | | | No ORF goes through the whole transcript. |
| GGO-NOTCH2NL-2 | NOTCH2NLR | chr1 117331132 117364496 | - | 4 | 235 | 5 | 11,825 | PPA/PTR/GGO | ORF has exons 2-5 of NOTCH2NLR |
| | Fusion NOTCH2NL-NBPF | chr1 117294017 117373312 | | 24 | 1114 | 11 | | | ORF has exons 2-4 of NOTCH2NLR & NBPF exons |
| | Fusion PDE4DIP-NOTCH2NL-NBPF | chr1 117294017 117447550 | | -- | -- | 16 | | | No ORF goes through the whole transcript. |
| GGO-NOTCH2NL-3 | Fusion LRIG2-NOTCH2NL-NBPF | chr1 125591450 125661487 | - | -- | 1662 | 5 | 5385 | GGO | Has exons of LRIG2, exon 3 of NOTCH2NLR, & NBPF exons but no M start |
| | Fusion NOTCH2NL-NBPF | chr1 125591876 125635112 | | -- | -- | 8 | | | Has exon 3 of NOTCH2NLR and NBPF exons but no exon 1/2 and no M start |
| GGO-NOTCH2NL-4 | Fusion LRIG2-NOTCH2NL-NBPF | chr1 129364798 129435830 | + | -- | -- | 19 | 5,608 | GGO | Has 2 LRIG2 exons, exon 3 of NOTCH2NLR, & NBPF exons but early stop in exon 2 |
| | Fusion NOTCH2NL-NBPF | chr1 129389184 129435600 | | -- | -- | 2 | | | Has 3-4 of NOTCH2NLR & NBPF exons but no exon 1/2 and no M start |
| GGO-NOTCH2NL-5 | Fusion MAGI3-NOTCH2NL-NBPF | chr1 129726316 129995372 | + | -- | 1487 | 2 | 5,588 | GGO | Has exons of MAGI3, exon 3 of NOTCH2NLR, & then NBPF exons but no M start |
| | Fusion NOTCH2NL-NBPF | chr1 129964606 129995476 | | -- | -- | 6 | | | Has 3-4 of NOTCH2NLR & NBPF exons but no exon 1/2 and no M start |
| GGO-NOTCH2NL-6 | Fusion NOTCH2NL-NBPF | chr1 130013152 130080198 | - | 13 | 783 | 72 | 39,879 | GGO | ORF has exon 1 of NOTCH2NLR & NBPF exons |
| GGO-NOTCH2NL-7 | NOTCH2NLR | chr1 130265287 130274296 | - | -- | -- | 14 | 11,779 | GGO | Has 3-5 of NOTCH2NLR but no exon 1/2 and no M start |
| | Fusion NOTCH2NL-NBPF | chr1 130222211 130280603 | | -- | -- | 15 | | | Has 3-4 of NOTCH2NLR & NBPF exons but no exon 1/2 and no M start |
| | Fusion BRD9-NOTCH2NL | chr1 130264240 130305169 | | -- | -- | 8 | | | No ORF that goes through the whole transcript. |
| | Fusion BRD9-NOTCH2NL-NBPF | chr1 130221951 130304952 | | -- | -- | 4 | | | No ORF that goes through the whole transcript. |
| HSA-NOTCH2 | NOTCH2 | chr1 119924936 120082923 | - | -- | 2471 | -- | -- | PTR/PPA/GGO/HSA | -- |
| HSA-NOTCH2NLA | NOTCH2NLA | chr1 145272197 145345902 | - | 5 | 236 | -- | 11,336 | HSA | |
| HSA-NOTCH2NLB | NOTCH2NLB | chr1 146108509 146181500 | + | 5 | 249 | -- | 11,375 | HSA | |
| HSA-NOTCH2NLC | NOTCH2NLC | chr1 148535272 148596912 | + | 5 | 236 | -- | 11,726 | HSA | |
| HSA-NOTCH2NLR | NOTCH2NLR | chr1 120737165 120807117 | + | 5 | 274 | -- | 11,635 | HSA | |
| | [a]Fusion NOTCH2NL-NBPF | chr1 120737168 120851627 | | 36 | 1673 | -- | | | |

[a]Fusion exists in all human *NOTCH2NL* but only *NOTCH2NLR* fusion is used for comparison.

**Table S4. Population origin of HPRC[a] haplotypes resolved[b] across the *NOTCH2NL* locus, related to Figures 4 and 5.**

| superpopulation | superpopulation total | population | population total |
|---|---|---|---|
| | | African Caribbean in Barbados | 11 |
| | | Gambian in Western Divisions - Mandinka | 15 |
| AFR | 34 | Mende in Sierra Leone | 6 |
| | | Yoruba in Ibadan, Nigeria | 2 |
| | | African Ancestry in Southwest USA | 2 |
| | | Maasai in Kinyawa, Kenya | 1 |
| | | Puerto Rican in Puerto Rico | 14 |
| AMR | 23 | Colombian in Medellin, Colombia | 2 |
| | | Peruvian in Lima, Peru | 7 |
| | | Kinh in Ho Chi Minh City, Vietnam | 1 |
| EAS | 7 | Southern Han Chinese in Hu Nan Province, China | 5 |
| | | Chinese Ancestry in USA | 1 |
| EUR | 1 | Ashkenazim Jewish | 1 |
| SAS | 1 | Punjabi in Lahore, Pakistan | 1 |

[a]HPRC: Human Pangenome Reference Consortium

[b]This includes haplotypes that are not completely assembled across the centromere but have no gaps or collapses between *NOTCH2* and *NOTCH2NLR*, and *NOTCH2NLA/B/C*, respectively.

| Table S5. ***NOTCH2NL/GFP* gBlocks for measuring protein expression, related to STAR Methods.** | |
|---|---|
| **Sequence name** | **5' – sequence – 3'** |
| HA-NOTCH2tv-IRES-NheI-FseI gBlock | GGTCTAGAGCTAGCGAATTCGCCGGTGCCACCATGTACCCATACGATGTTCCAGA<br>TTACGCTCCCGCCCTGCGCCCCGCTCTGCTGTGGGCGCTGCTGGCGCTCTGGCT<br>GTGCTGCGCGGCCCCCGCGCATGCATTGCAGTGTCGAGATGGCTATGAACCCTG<br>TGTAAATGAAGGAATGTGTGTTACCTACCACAATGGCACAGGATACTGCAAATGTC<br>CAGAAGGCTTCTTGGGGGAATATTGTCAACATCGAGACCCCTGTGAGAAGAACCG<br>CTGCCAGAATGGTGGGACTTGTGTGGCCCAGGCCATGCTGGGGAAAGCCACGTG<br>CCGATGTGCCTCAGGGTTTACAGGAGAGGACTGCCAGTACTCAACATCTCATCCA<br>TGCTTTGTGTCTCGACCCTGCCTGAATGGCGGCACATGCCATATGCTCAGCCGGG<br>ATACCTATGAGTGCACCTGTCAAGTCGGGTTTACAGGTAAGGAGTGCCAATGGAC<br>GGATGCCTGCCTGTCTCATCCCTGTGCAAATGGAAGTACCTGTACCACTGTGGCC<br>AACCAGTTCTCCTGCAAATGCCTCACAGGCTTCACAGGGCAGAAATGTGAGACTG<br>ATGTCAATGAGTGTGACATTCCAGGACACTGCCAGCATGGTGGAACCTGCCTCAA<br>CCTGCCTGGTTCCTACCAGTGCCAGTGCCCTCAGGGCTTCACAGGCCAGTACTGT<br>GACAGCCTGTATGTGCCCTGTGCACCCTCACCTTGTGTCAATGGAGGTACCTGTC<br>GGCAGACTGGTGACTTCACTTTTGAGTGCAACTGCCTTCCAGAAACAGTGAGAAAT<br>AAGAGGAACAGAGCTCTGGGAAAGGGACAGGCAAGTCTGGAATGGAAAAGAACA<br>TGAGTCGACCGCTTGGAATAAGGCCGGTGTGCGTTTGTCTATATGTTATTTTCCAC<br>CATATTGCCGTCTTTTGGCAATGTGAGGGCCCGGAAACCTGGCCCTGTCTTCTTG<br>ACGAGCATTCCTAGGGGTCTTTCCCCTCTCGCCAAAGGAATGCAAGGTCTGTTGA<br>ATGTCGTGAAGGAAGCAGTTCCTCTGGAAGCTTCTTGAAGACAAACAACGTCTGTA<br>GCGACCCTTTGCAGGCAGCGGAACCCCCCACCTGGCGACAGGTGCCTCTGCGGC<br>CAAAAGCCACGTGTATAAGATACACCTGCAAAGGCGGCACAACCCCAGTGCCACG<br>TTGTGAGTTGGATAGTTGTGGAAAGAGTCAAATGGCTCTCCTCAAGCGTATTCAAC<br>AAGGGGCTGAAGGATGCCCAGAAGGTACCCCATTGTATGGGATCTGATCTGGGG<br>CCTCGGTGCACATGCTTTACATGTGTTTAGTCGAGGTTAAAAAAACGTCTAGGCCC<br>CCCGAACCACGGGGACGTGGTTTTCCTTTGAAAAACACGATGATAAGATCTGCGA<br>TCTAAGTAAGCTTGGCATTCCGGTACTGTTGGTAAAGCCACCATGGAATCCGGCC<br>GGCCCGAATTCGGC |
| HA-NOTCH2NLB pEF1A Gibson gBlock | GATGTTCCAGATTACGCTTGTCGAGATGGCTATGAACCCTGTGTAAATGAAGGAAT<br>GTGTGTTACCTACCACAATGGCACAGGATACTGCAAATGTCCAGAAGGCTTCTTG<br>GGGGAATATTGTCAACATCGAGACCCCTGTGAGAAGAACCGCTGCCAGAATGGTG<br>GGACTTGTGTGGCCCAGGCCATGCTGGGGAAAGCCACGTGCCGATGTGCCTCAG<br>GGTTTACAGGAGAGGACTGCCAGTACTCGACATCTCATCCATGCTTTGTGTCTCGA<br>CCTTGCCTGAATGGCGGCACATGCCATATGCTCAGCCGGGATACCTATGAGTGCA<br>CCTGTCAGGTCGGGTTTACAGGTAAGGAGTGCCAATGGACCGATGCCTGCCTGTC<br>TCATCCCTGTGCAAATGGAAGTACCTGTACCACTGTGGCCAACCAGTTCTCCTGCA<br>AATGCCTCACAGGCTTCACAGGGCAGAAGTGTGAGACTGATGTCAATGAGTGTGA<br>CATTCCAGGACACTGCCAGCATGGTGGCATCTGCCTCAACCTGCCTGGTTCCTAC<br>CAGTGCCAGTGCCTTCAGGGCTTCACAGGCCAGTACTGTGACAGCCTGTATGTGC<br>CCTGTGCACCCTCGCCTTGTGTCAATGGAGGCACCTGTCGGCAGACTGGTGACTT<br>CACTTTTGAGTGCAACTGCCTTCCAGAAACAGTGAGAAGAGGAACAGAGCTCTGG<br>GAAAGAGACAGGGAAGTCTGGAATGGAAAAGAACACGATGAGAATTAGGTCGACC<br>GCTTGGAATAA |
| HA-NOTCH2NLR pEF1A Gibson gBlock | GATGTTCCAGATTACGCTCCCGCCCTGCGTCCCGCTCTGCTGTGGGCGCTGCTG<br>GCGCTCTGGCTGTGCTGGGCGGCCCCCGCGCATGCATTGCAGTGTCGAGATGGC<br>TATGAACCCTGTGTAAATAAAGGAATGTGTGTTACCTACCACAGTGGCACAGGATA<br>CTGCAAATGTCCAGAAGGCTTCTTGGGGGAATATTGTCAACATCGAGACCCCTGT<br>GAGAAGAACCGCTGCCAGAATGGTGGGACTTGTGTGGCCCAGGCCATGCTGGGG<br>AAAGCCACGTGCCGGTGTGCCTCAGGGTTTACAGGAGAGGACTGCCAGTACTCG<br>ACACCTCATCCATGCTTTGTGTCTCGACCTTGCCTGAATGGCGGCACATGCCATAT<br>GCTCAGCCGGGATACCTATGAGTGCACCTGTCAAGTCGGGTTTACAGGTAAGGAG<br>TGCCAATGGACCGATGCCTGCCTGTCTCATCTCTGTGCAAATGGAAGTACCTGTA<br>CCACTGTGGCCAACCAGTTCTCCTGCAAATGCCTCACAGGCTTCACAGGGCAGAA<br>GTGTGAGACTGATGTCAATGAGTGTGACATTCCAGGACACTGCCAGCATGGTGGC<br>ACCTGCCTCAACCTGCCTGGTTCCTACCAGTGCCAGTGCCTTCAGGGCTTCACAG<br>GCCAGTACTGTGACAGACTGTATGTGCCCTGTGCACACTCGCCTTGTGTCAATGG<br>AGGCACCTGTCGGCAGACTGGTGACTTCACTTTTGAGTGCAACTGCCTTCCAGAA<br>ACAGTGAGAAATAAGAGGAACAGAGCTCTGGGAAAGAGACAGGCAAGTCTGGAAT<br>GGAAAAGAACACGATGAGTCGACCGCTTGGAATAA |

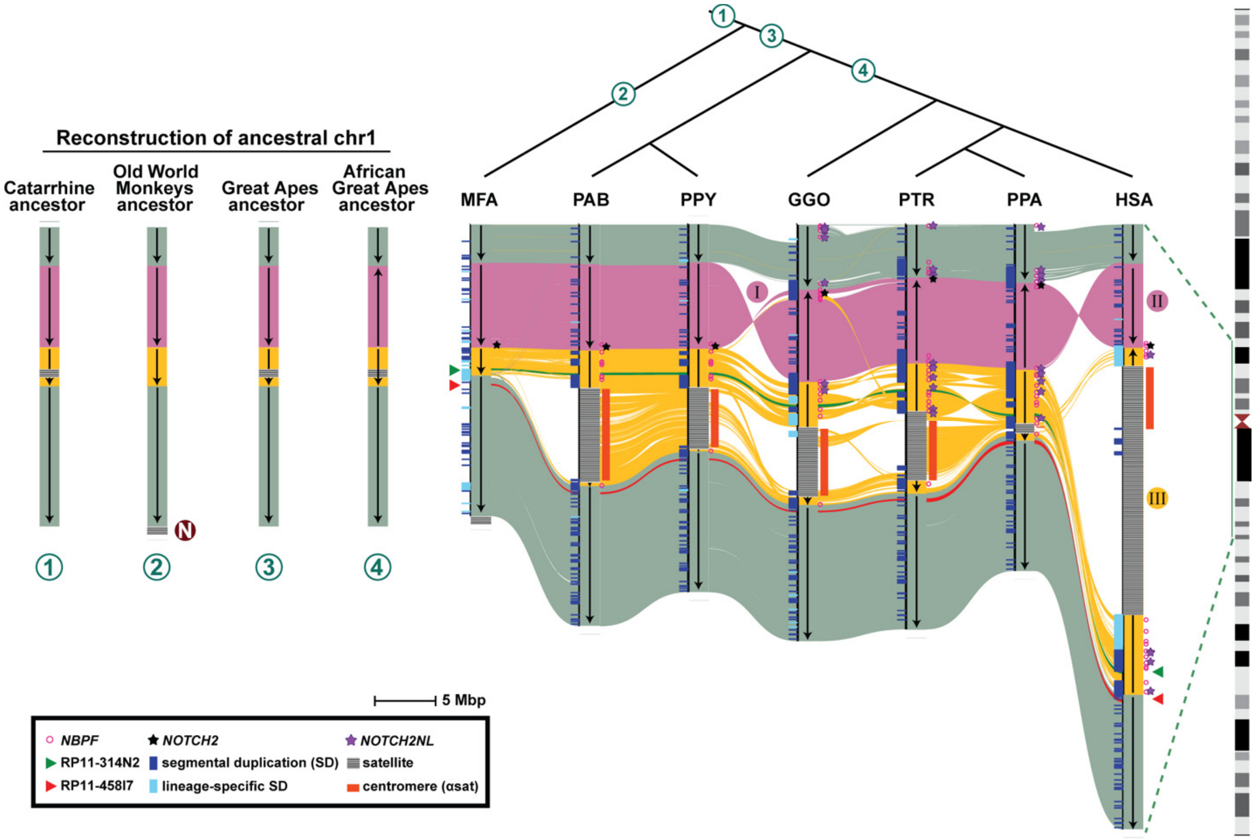


**Figure S1. Ape evolutionary rearrangement and expansion of human chromosome 1p21.2-q23.2, related to Figure 2.** The genomic structure of chromosome 1p21.2-q23.2 region is compared among macaque (MFA), Sumatran orangutan (PAB), Bornean orangutan (PPY), gorilla (GGO), chimpanzee (PTR), bonobo (PPA), and human (HSA) with annotations that include ancestral *NOTCH2* (black stars), *NOTCH2NL* duplications (purple stars), *NBPF* duplications (pink circles), and the centromere (orange bars). The circled numbers represent previous ancestral states of chromosome 1. The circled N represents a centromere repositioning event (N, neocentromere). Three distinct evolutionary inversions are predicted (I, II, III). Two probes (RP11-314N2, green, and RP11-458I7, red) used in FISH analyses from Szamalek et al. (2006)[S1] are shown (green and red triangles). Both probes map to the q-arm in humans, with the green probe located inside the inverted region and the red probe outside. FISH data from Szamalek et al. (2006)[S1] revealed that in chimpanzee the green probe maps to the region homologous to the human p-arm, while the red probe maps to the q-arm. Sequence analysis supports the FISH mapping and shows that in great apes the sequence of the two probes (represented as red and green lines in the SVbyEye) map on opposite sides of the centromere.

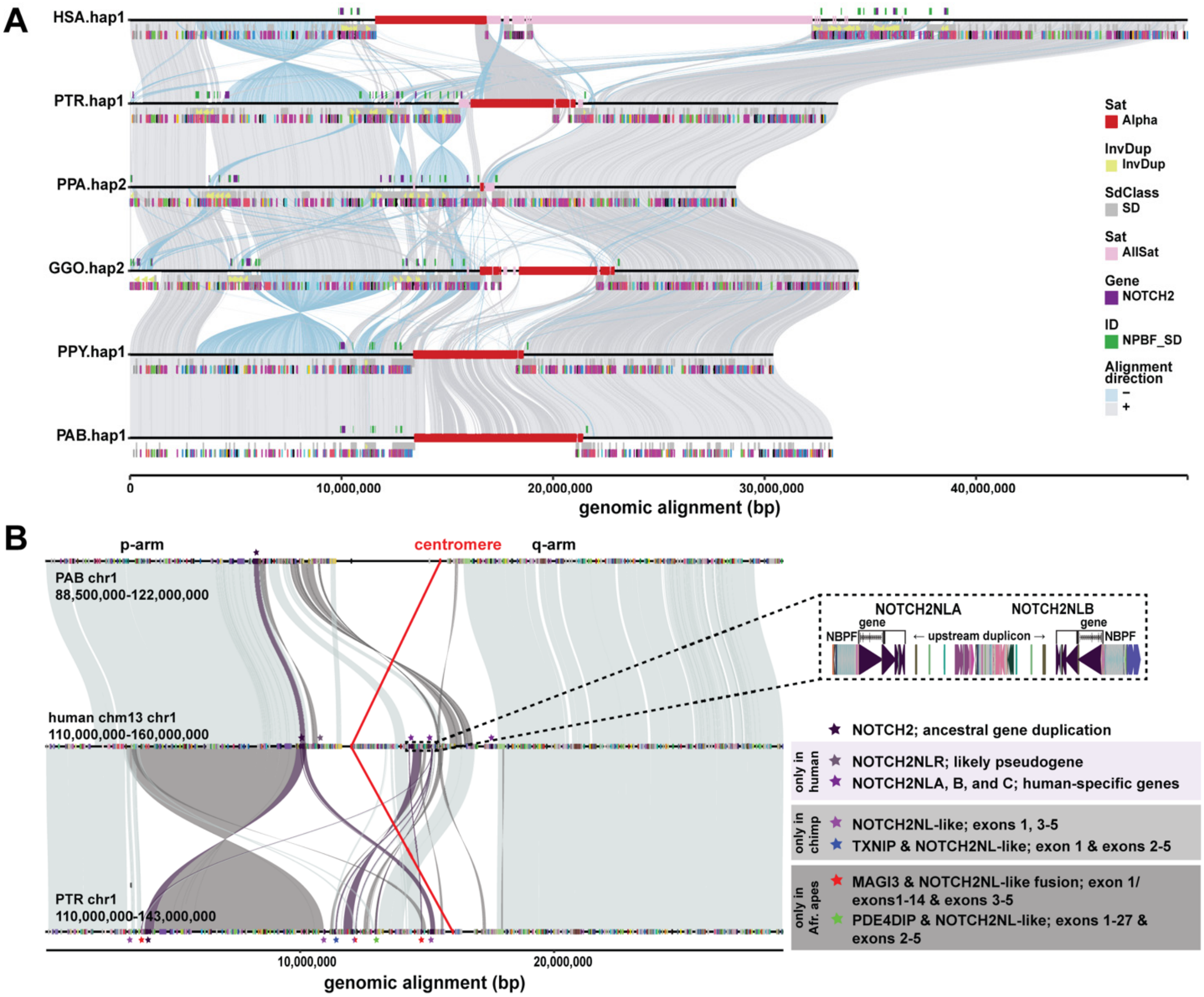


**Figure S2. NHP and human chromosome 1 evolutionary rearrangements, related to Figure 2.**
**A)** Stacked SVbyEye plot of human chromosome 1 compared to syntenic chromosomes in other apes over 30-50 Mbp of the pericentromeric region. Directly aligned sequence (gray) is compared to inversion rearrangements (light blue) with various annotations including *NOTCH2NL* and *NBPF* genes, satellites, and others. **B)** A zoomed-in comparison including only chimpanzee, orangutan, and human where centromeric sequence has been masked to highlight the non-orthologous location of NHA *NOTCH2NL* transcripts (colored stars) as described by Fiddes et al. (2018)[S2].

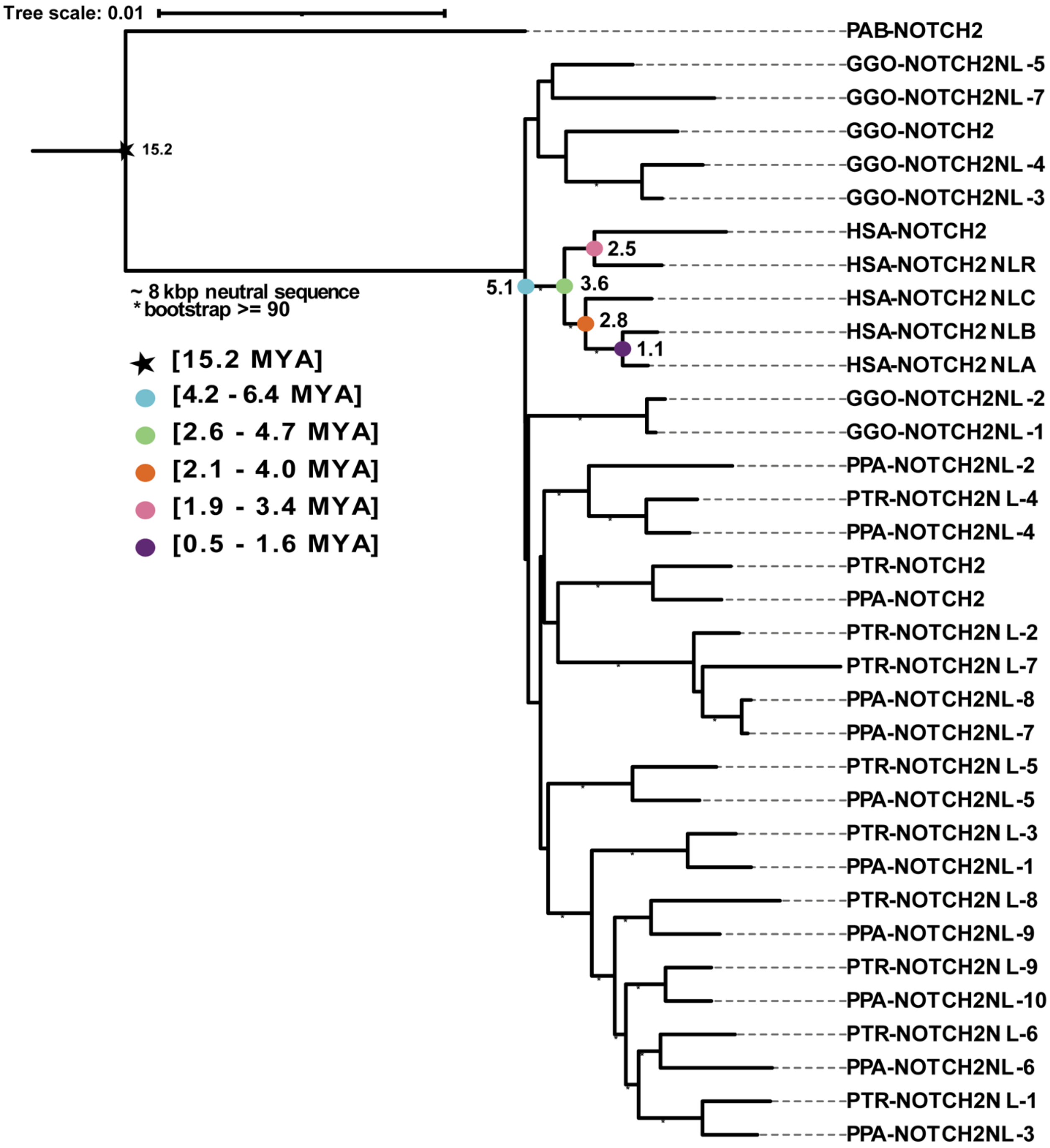


**Figure S3. Extended NHA phylogeny, related to Figure 3.** An alternate maximum likelihood phylogeny based on an MSA of 8 kbp from intron 3 of *NOTCH2/NL* sequence from paralogs of five ape species, using Sumatran orangutan as an outgroup. Boots representing 25/26 NHA homologs. Bootstrap support (>90%) is indicated (asterisk) and is less robust than the phylogeny in Figure 3A but contains more taxa. Estimated divergence times of human paralogs and their confidence intervals are indicated (multicolored dots). Timings were based on human–orangutan divergence time of 15.2 MYA (Methods).

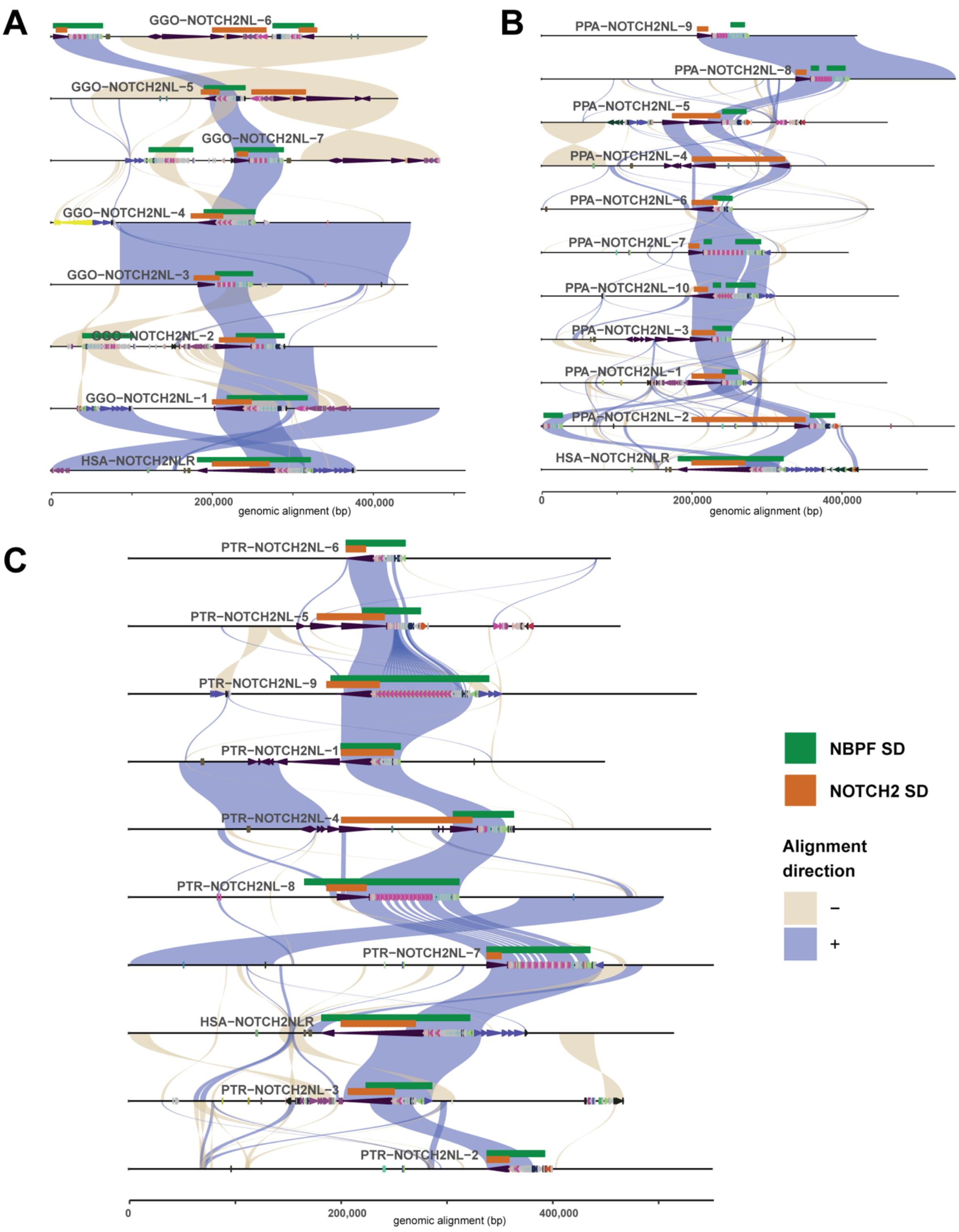


**Figure S4. Ladder alignment of NHA *NOTCH2NL* homologs, related to Figure 3.** Gorilla **(A)**, bonobo **(B)**, and chimpanzee **(C)** self-alignments (with human *NOTCH2NLR*) show a consistent association between the *NOTCH2/NL* duplication (purple) and the core duplicon *NBPF* (green). However, the breakpoints of these alignments are largely different not just from humans but also each other.

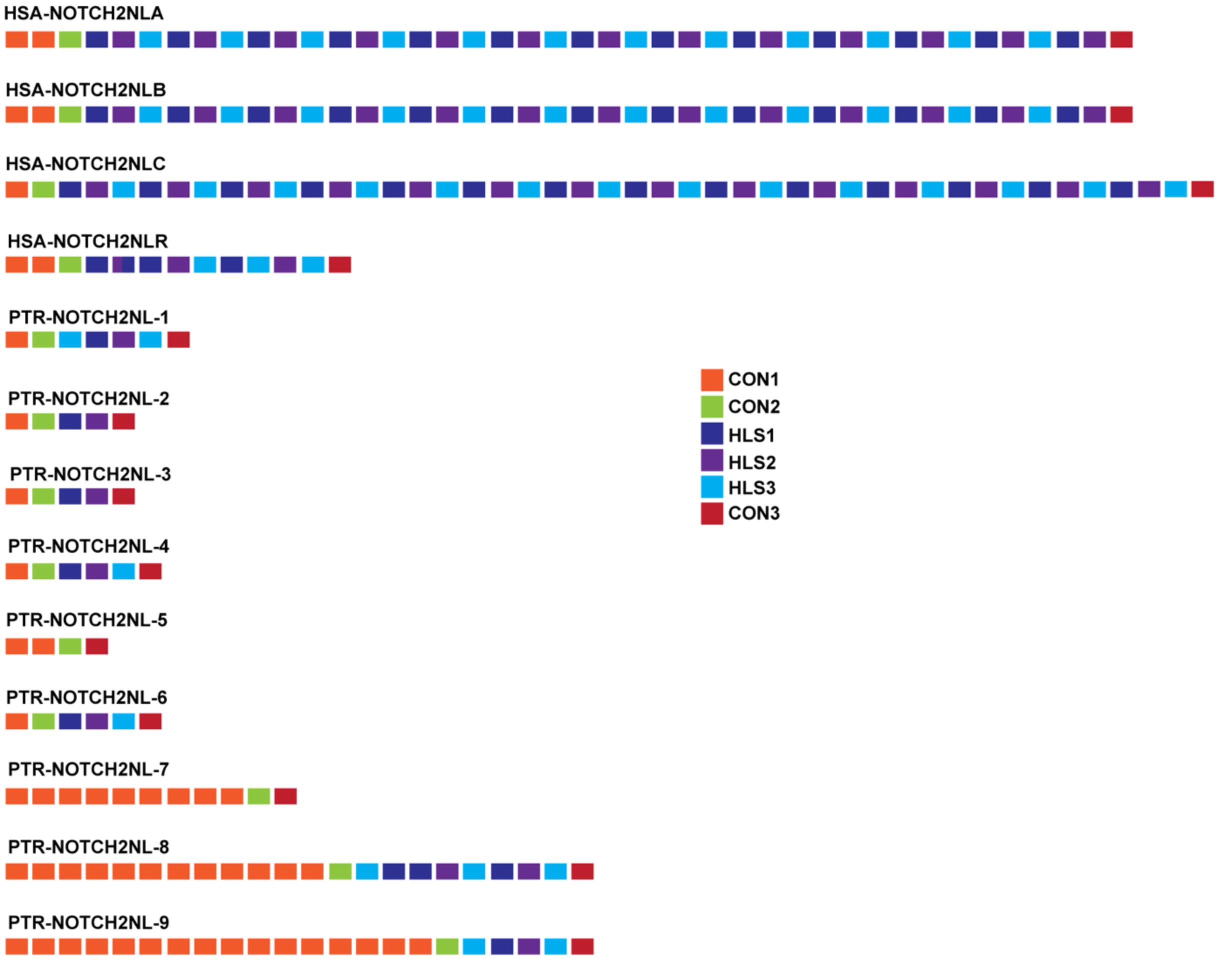


**Figure S5. NBPF duplicon analysis, related to Figure 3.** For each of the HSA and PTR *NOTCH2NL* copies, the *NBPF* directly downstream is represented in terms of the DUF1220 domains present—CON1, CON2, CON3, HLS1, HLS2, HLS3, based on Fiddes et al. (2019)[S3]. The *NBPF* copies downstream of *NOTCH2NL* copies in PPA and GGO show similar patterns of DUF1220 repeats as well.

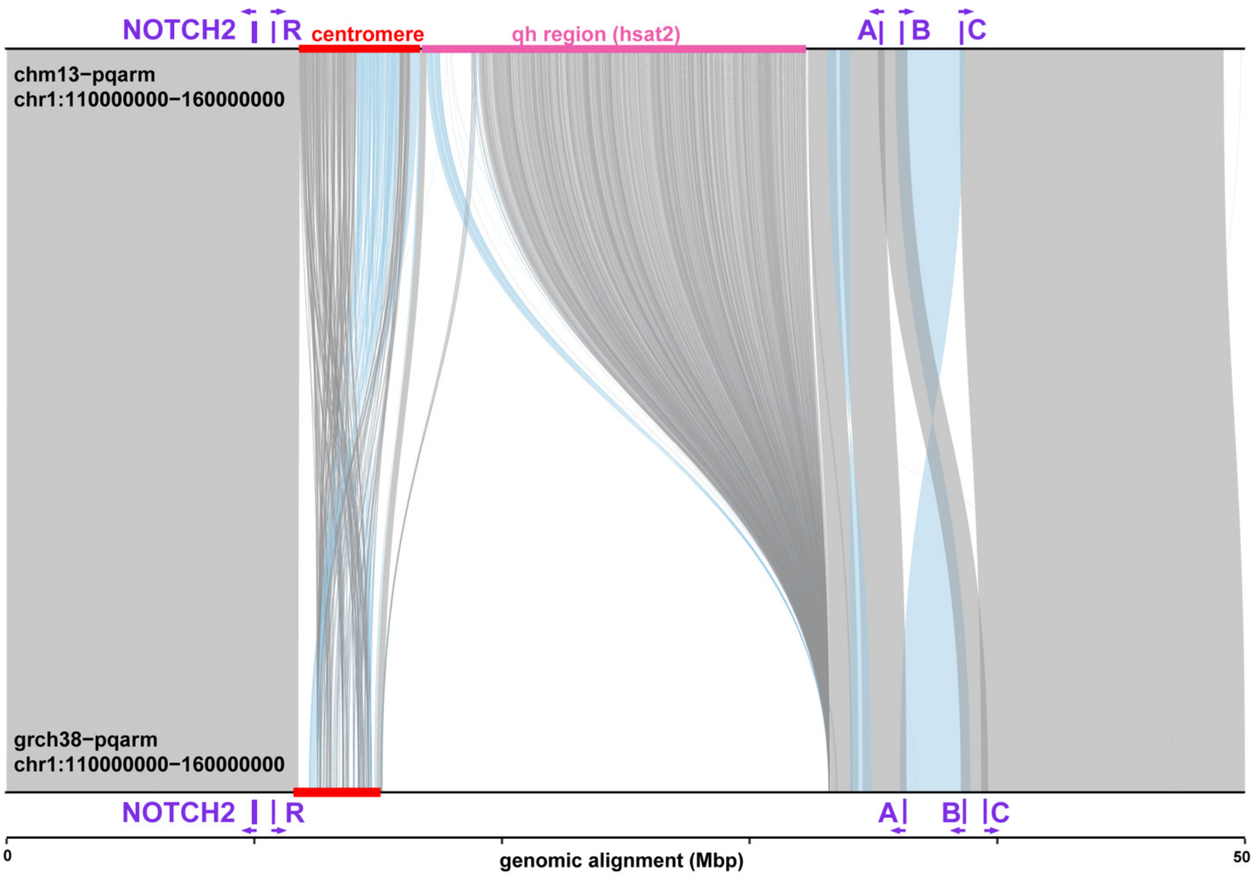


**Figure S6. T2T-CHM13 comparison to GRCh38, related to STAR Methods.** Organization of the *NOTCH2/NL* region in the completed T2T haploid assembly CHM13 against the previous reference, GRCh38. The T2T-CHM13 assembly has expanded the sequence and annotations that cover the centromere (red) and qh region (pink), which are two large satellite sequences that separate the two *NOTCH2/NL* (purple) loci. A long inversion around *NOTCH2NLB* that may include both *NOTCH2NLA/C* on either side changes the gene's orientation between the two assemblies (relevant alignments have increased color opacity).

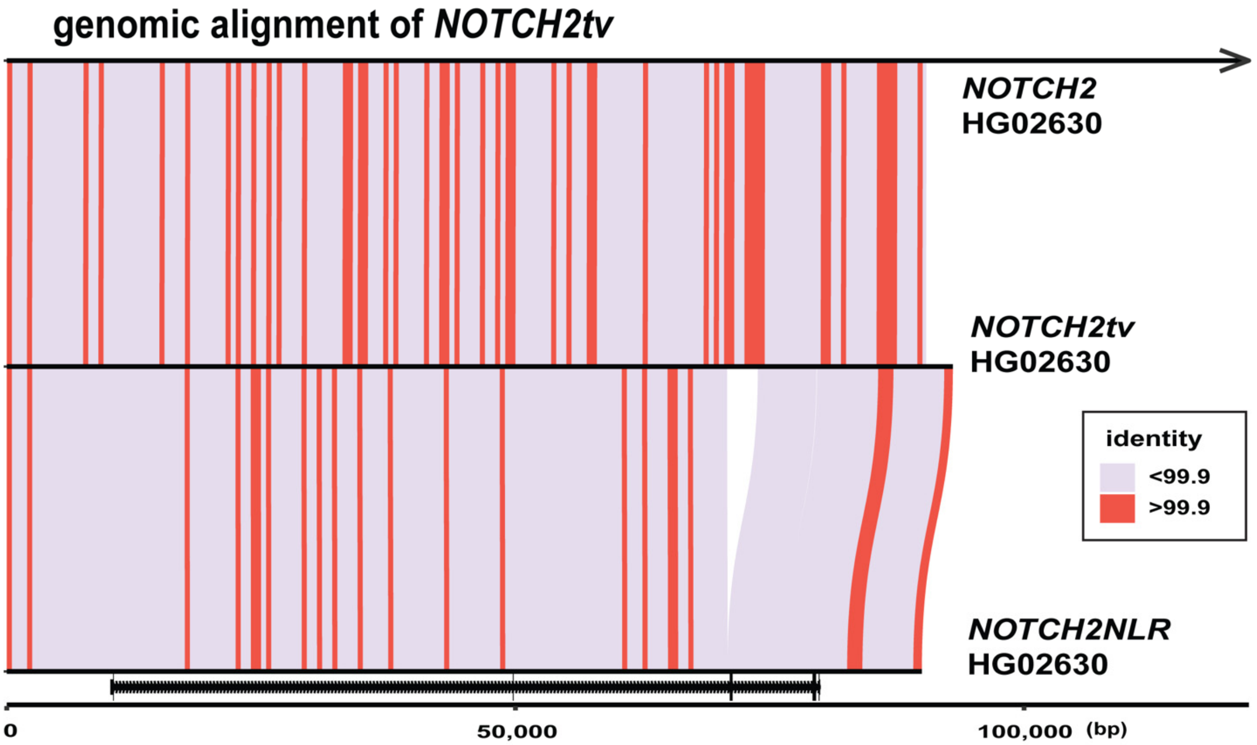


**Figure S7. NOTCH2NLR to NOTCH2tv gene conversion, related to Figure 5.** Nucleotide alignment of *NOTCH2tv* (middle) to ancestral *NOTCH2* (top) and *NOTCH2NLR* (bottom) confirms larger stretches of near perfect sequence identity (red≥99.9%) between *NOTCH2tv and NOTCH2*, consistent with IGC.

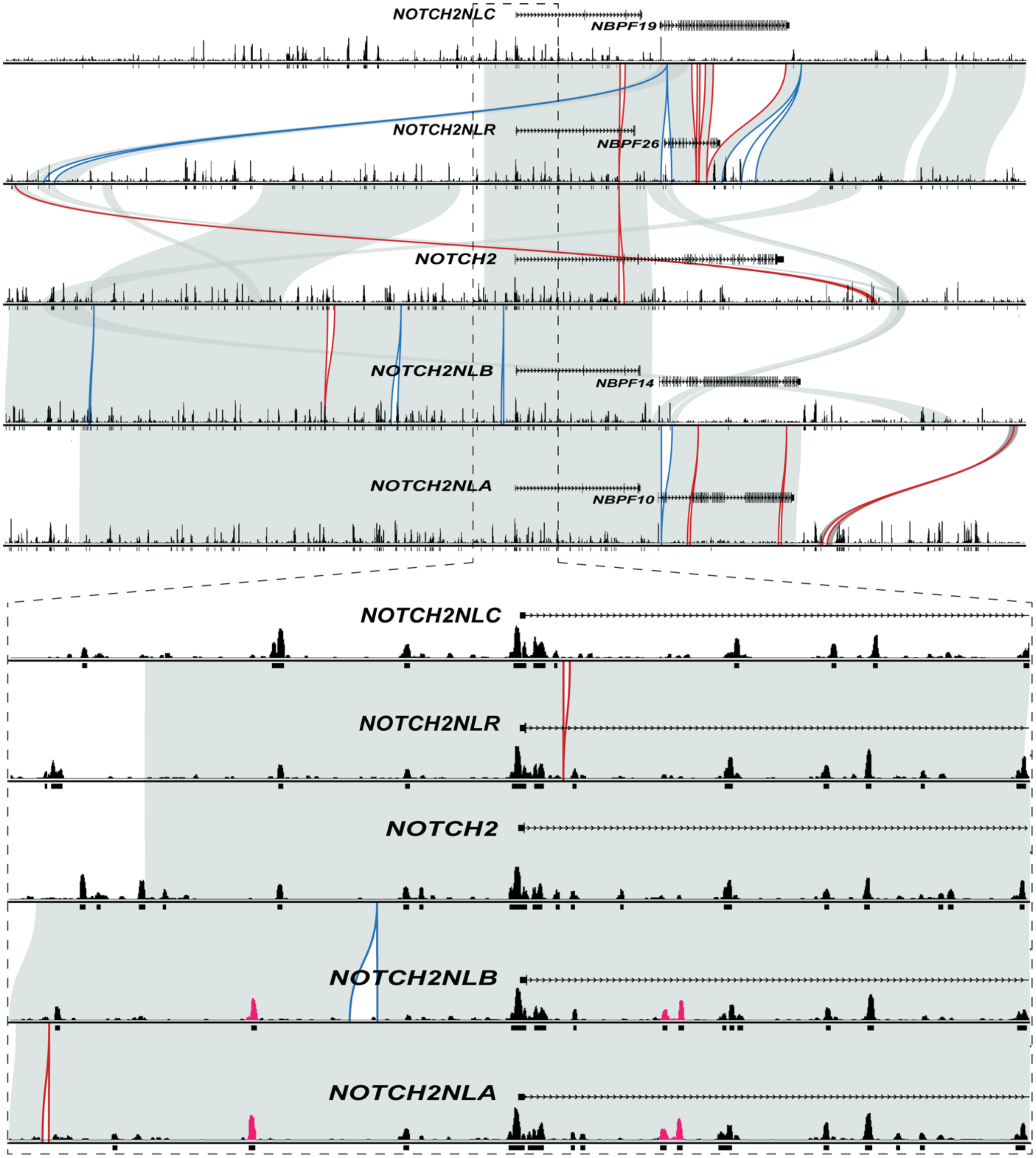


**Figure S8. Regulatory architecture of T2T-CHM13, related to STAR Methods.** Fiber-seq chromatin actuation peaks for each T2T-CHM13 *NOTCH2/NL* paralog in the context of homology (gray) and gene model 300 kbp on either side of the TSS. Examples of signals of paralog-specific actuation (pink) are shown in pop-out panel (25 kbp on either side of TSS) in *NOTCH2NLA* and *NOTCH2NLB*. Structural variants like insertions (blue) and deletions (red) are outlined.

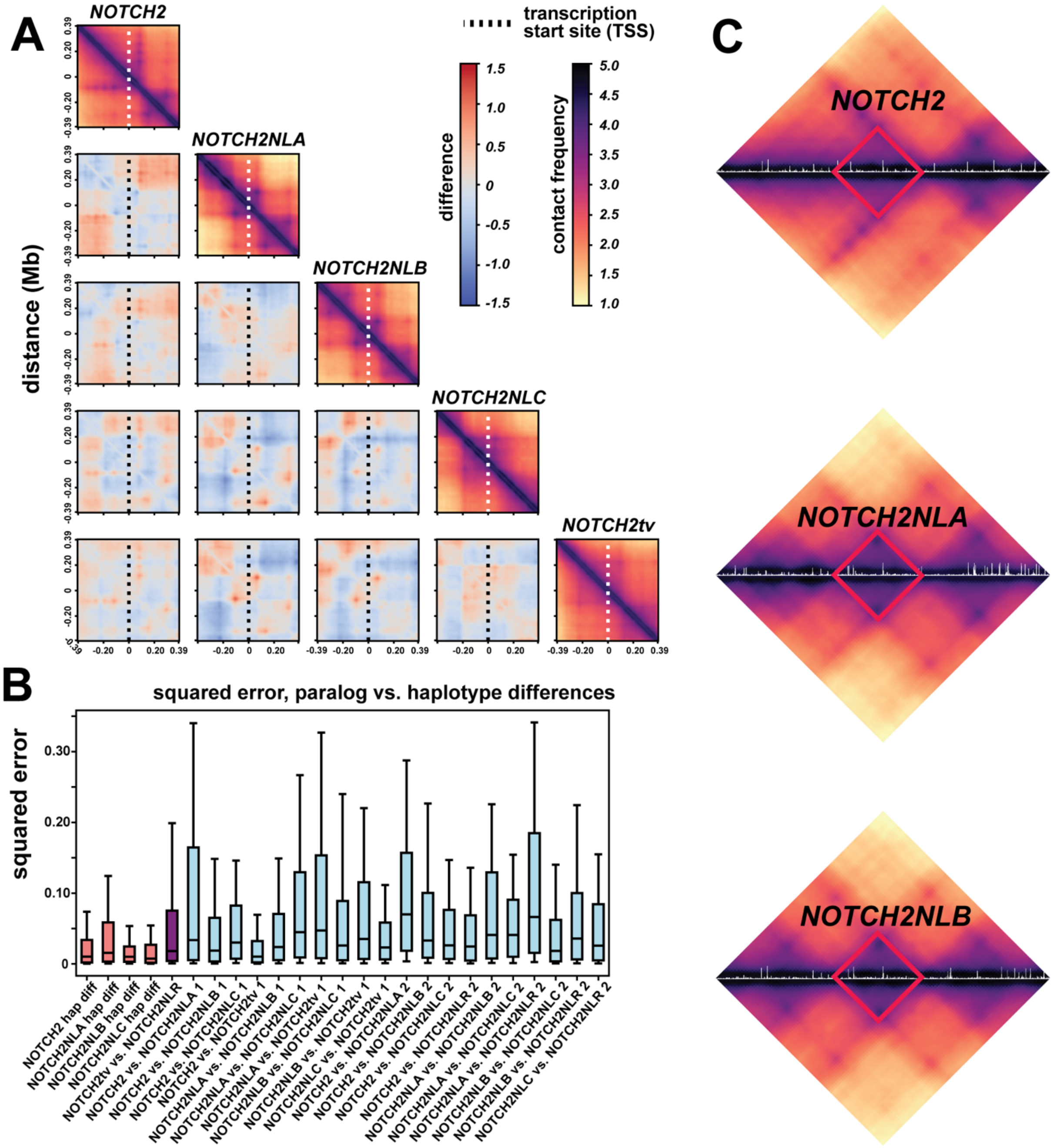


**Figure S9. Predicted TADs and associated regulatory architectures in *NOTCH2NL* paralog regions, related to Figure 6. A)** FiberFold TAD *NOTCH2NL* paralog predictions and pairwise heatmap of contact similarities and differences. Dotted lines represent gene TSS. **B)** Squared error chromatin accessibility differences between haplotypes and pairwise paralog comparisons. **C)** Fiber-seq chromatin actuation overlaid on TAD predictions for *NOTCH2*, *NOTCH2NLA*, and *NOTCH2NLB* to show which regulatory peaks are included in contact regions.

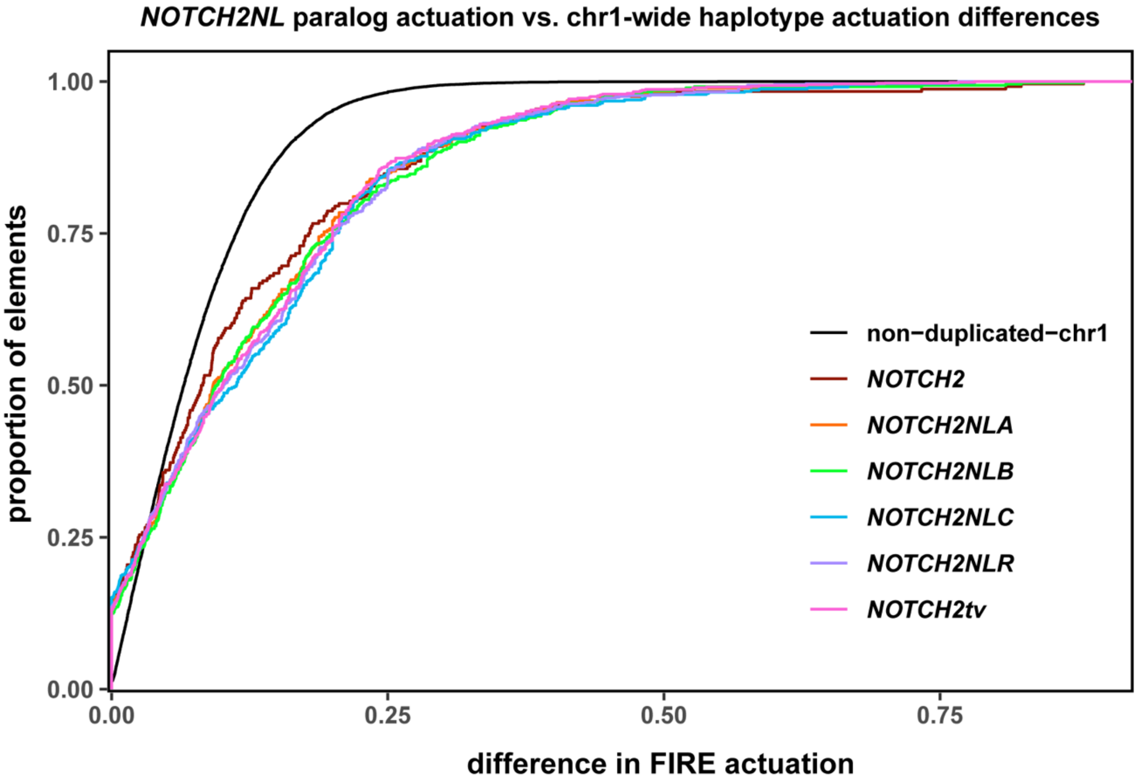


**Figure S10. Comparison of *NOTCH2NL* paralog and haplotype actuation, related to Figure 6.** The model of expected haplotype differences on chromosome 1 using brain organoid data mapped to T2T-CHM13 (black) shows a higher proportion of regulatory elements with less actuation difference than what is seen for the different *NOTCH2NL* paralogs.

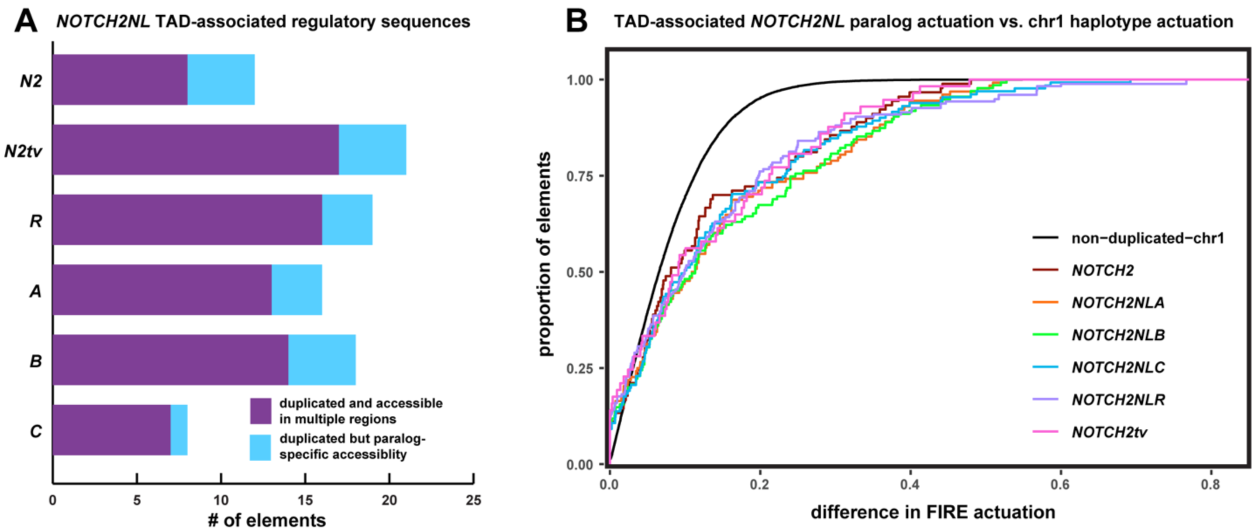


**Figure S11. TAD-restricted regulatory analysis, related to Figure 6. A)** Bar graph showing categorization of the TAD-restricted accessible elements surrounding each *NOTCH2NL* paralog based on the presence of duplicate sequence and accessibility at that sequence on the different paralogs.
**B)** A TAD-restricted model of expected haplotype differences on chromosome 1 using brain organoid data mapped to T2T-CHM13 (black) also shows a higher proportion of regulatory elements with less actuation difference than what is seen for the different *NOTCH2NL* paralogs.

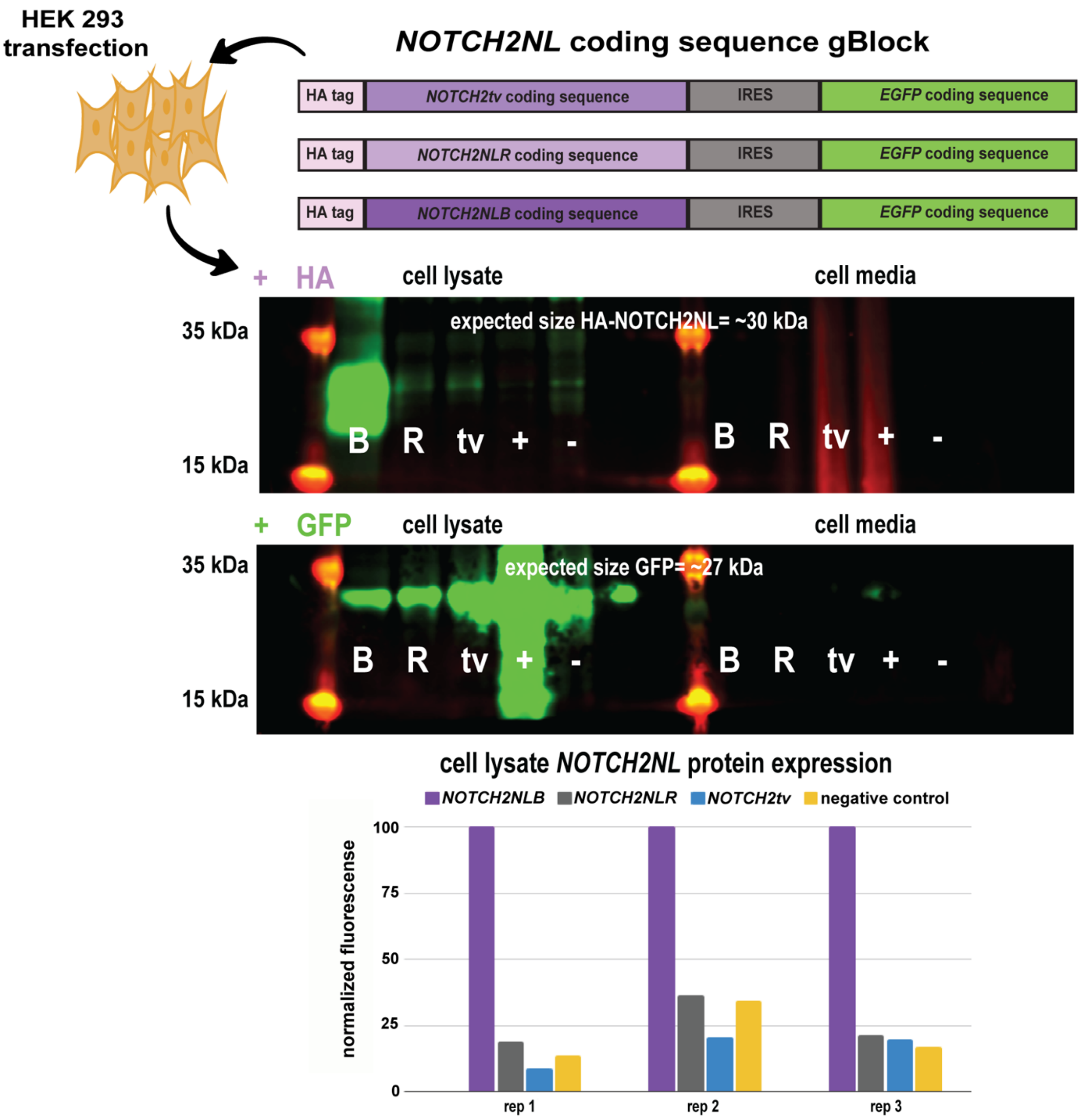


**Figure S12. NOTCH2tv protein expression, related to STAR Methods.** gBlocks containing an HA antibody tag, *NOTCH2NL* CDS, and *EGFP* CDS were cloned into vector DNA and transfected into HEK293 cells. Antibody staining for HA (NOTCH2NL) and GFP were done on both the cell lysate and media. HA expression fluorescence was normalized using GFP for cell lysate, which shows stable expression of NOTCH2NLB only.

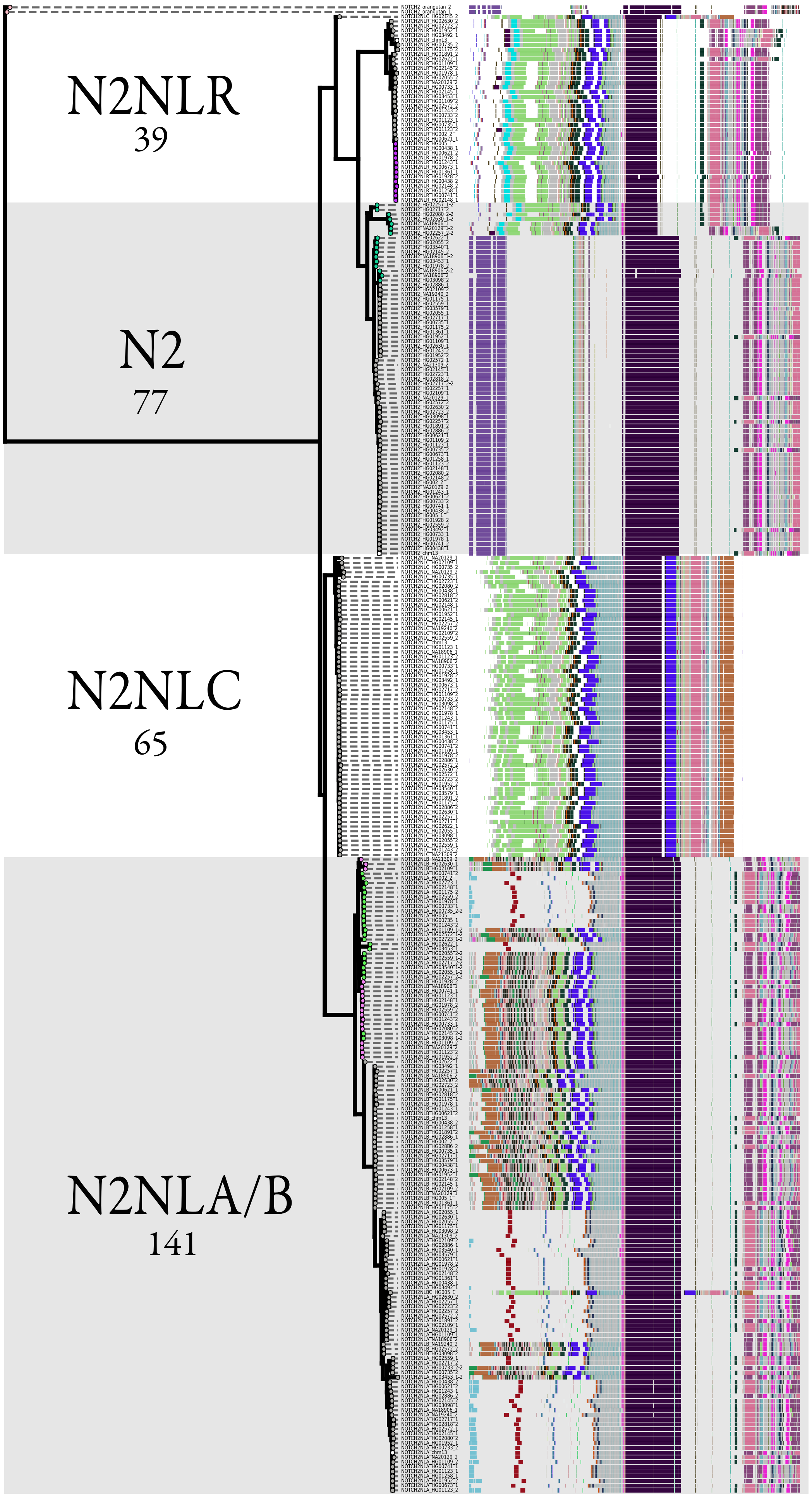

N2NLR
39
N2
77
N2NLC
65
N2NLA/B
141

**Figure S13. Phylogeny of human *NOTCH2NL* genetic variation, related to Figures 4 and 5.**
(figure on previous page)
